# Integrative network modeling reveals mechanisms underlying T cell exhaustion

**DOI:** 10.1101/582312

**Authors:** Hamid Bolouri, Mary Young, Joshua Beilke, Rebecca Johnson, Brian Fox, Lu Huang, Cristina Costa Santini, Christopher Mark Hill, Anne-Renee van der Vuurst de Vries, Paul Shannon, Andrew Dervan, Pallavur Sivakumar, Matthew Trotter, Douglas Bassett, Alexander Ratushny

**Affiliations:** Division of Human Biology, Fred Hutchinson Cancer Research Center, Seattle, WA 98109, USA.; Celgene Corporation, Seattle, WA 98102, USA.; Celgene Institute for Translational Research Europe (CITRE), Seville 41092, Spain.; Institute for Systems Biology, Seattle, WA 98109, USA.

**Author notes:** Correspondence should be addressed to: H.B. or A.R.

## Abstract

Failure to clear antigens causes CD8^+^ T cells to become increasingly hypo-functional, a state known as exhaustion. We combined manually extracted information from published literature with gene expression data from diverse model systems to infer a set of molecular regulatory interactions that underpin exhaustion. Topological analysis and simulation modeling of the network suggests CD8^+^ T cells undergo 2 major transitions in state following stimulation. The time cells spend in the earlier proliferative/pro-memory (PP) state is a fixed and inherent property of the network structure. Transition to the second state is necessary for exhaustion. Combining insights from network topology analysis and simulation modeling, we predict the extent to which each node in our network drives cells towards an exhausted state. We demonstrate the utility of our approach by experimentally testing the prediction that drug-induced interference with EZH2 function increases the proportion of proliferative/pro-memory cells in the early days post-activation.

## Introduction

Diverse mechanisms can limit T cell responses to tumors and immunotherapies^1^ (summarized in Supplementary Fig. 1). In particular, CD8^+^ T cells stimulated without the appropriate co-stimulatory signals can become anergic, while telomere erosion and/or DNA damage can result in T cell senescence^2^ over periods spanning *months/years*. T cell exhaustion (TCE) occurs in spite of physiologically appropriate stimulation when stimulation is prolonged to periods of *weeks*^2^, as is often the case for tumor-infiltrating lymphocytes and engineered T cells. TCE is defined by high expression of inhibitory receptors, and lowered capacity for proliferation, cytokine production, cytotoxic activity, and memory formation^2^. To date, immune checkpoint inhibitors targeting inhibitory receptors and other immunotherapies have shown great success in only *subsets* of patients across multiple cancers^3^. The limited efficacy of checkpoint inhibitors and their adverse effects in some patients^4,5^ suggest a need for a better understanding of TCE and the development of new TCE inhibitors. A better understanding of the mechanisms underlying TCE may improve the efficacy and reduce the adverse effects of immune checkpoint inhibitors, chimeric antigen receptor (CAR), engineered T cell receptor (TCR), and T cell engager immunotherapies^3^.

In response to acute infections, naïve and memory CD8^+^ T cells both undergo a stereotyped series of transcriptional state changes, summarized in Fig. 1a. CD8^+^ T cells receiving prolonged stimulation by chronic infections or cancer antigens undergo a qualitatively similar set of state changes (Fig. 1b), but become increasingly exhausted over time.

**Fig. 1.**
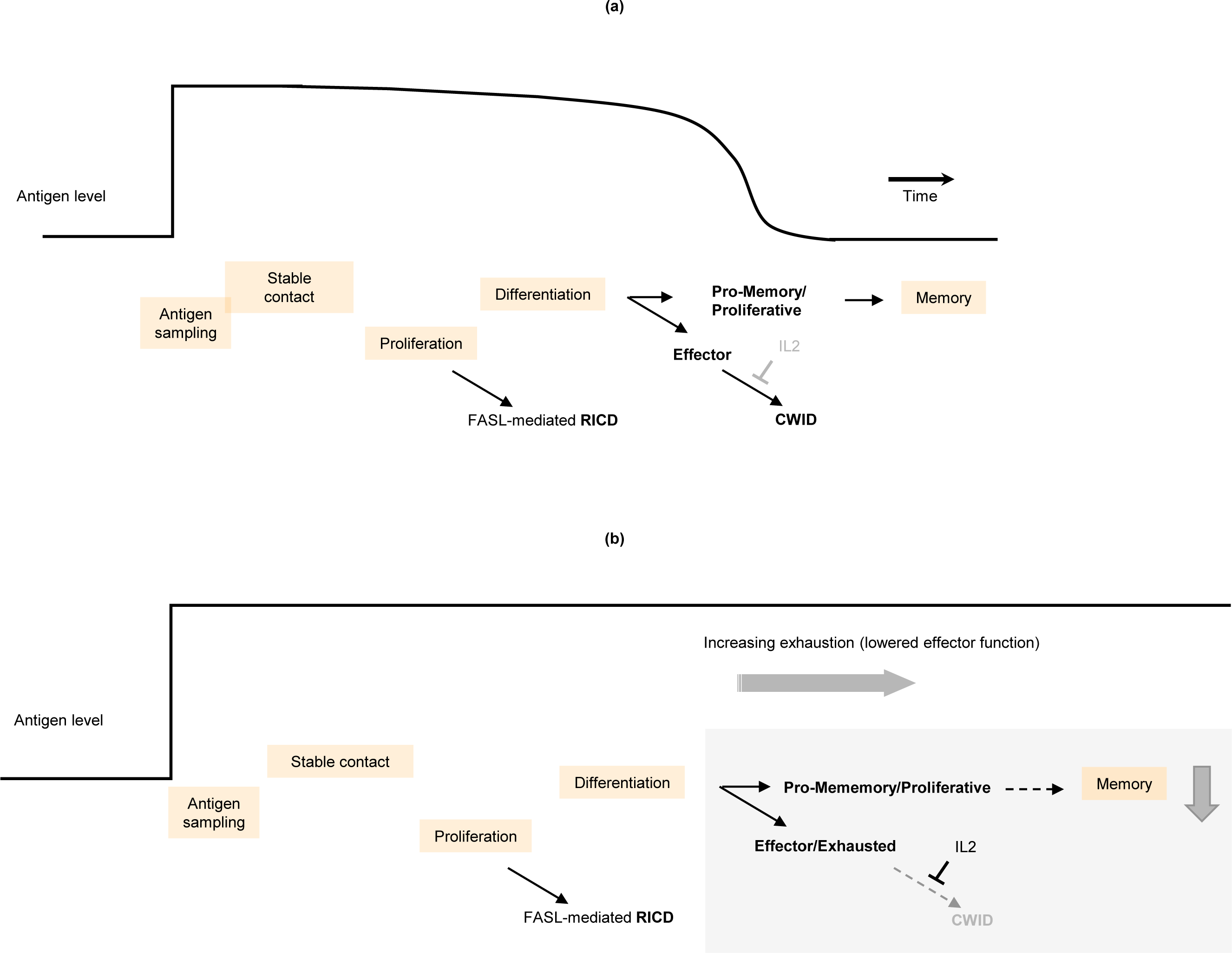
Multiple cellular state transitions underlie CD8^+^ T cell responses to acute and chronic co-stimulation. (**a**) Literature-based model of the acute response. CD8^+^ T cell activation starts with a complex and highly regulated process of antigen sampling (brief cell-cell contacts) that minimizes the risk of a response to self-antigens while sensitively detecting matching non-self antigens. Identification of a target cell results in the formation of a stable immune synapse and the activation of the T cell receptor (TCR) and its associated co-receptors (known as signals 2 and 3). TCR co-stimulation in turn leads to rapid and robust T cell proliferation. The magnitude of T cell amplification is controlled via a FAS/FASL mediated apoptotic signaling process known as restimulation-induced cell death (RICD). Activated CD8^+^ T cells differentiate into memory precursor and short-lived effector cells. Effector cells require cytokines such as IL2 for survival, and undergo cell death when antigen clearance leads to loss of cytokine signaling, a process known as cytokine withdrawal-induced cell death (CWID). After antigen clearance, memory CD8^+^ T cells are formed from the promemory/proliferative (PP) population in a process that takes several weeks. (**b**) In response to chronic co-stimulation, CD8^+^ T cells undergo state changes similar to (**a**). Prolonged activation reduces the proportion of memory-precursor cells, while persistence of cytokine signaling antagonizes CWID and leads to a hypo-functional effector/exhausted state.

**Fig. 2.**
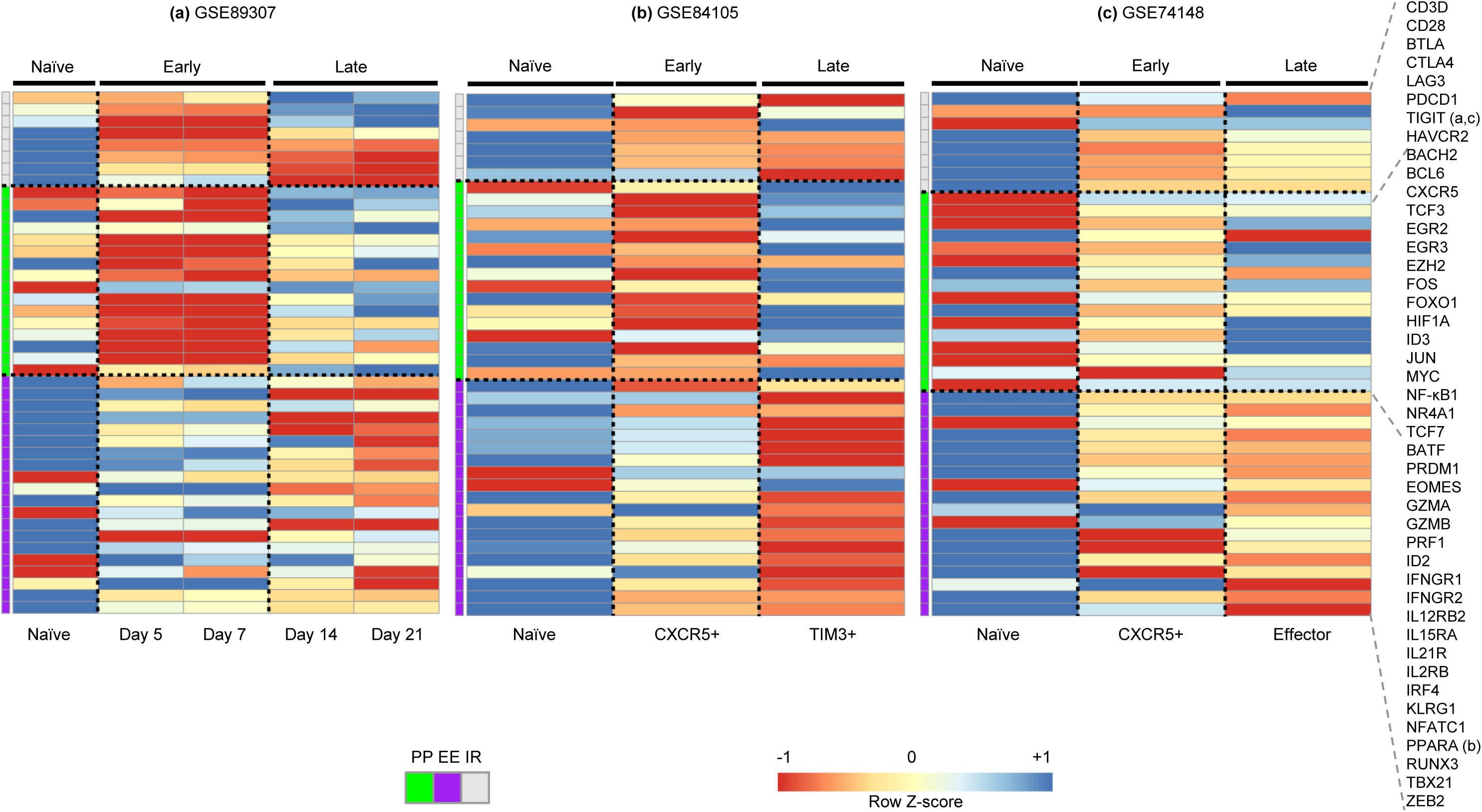
A two-state view of the key gene expression during CD8^+^ T cell activation and exhaustion. Shown are heatmaps comparing expression profiles of T cell exhaustion (TCE)-associated genes in CD8^+^ T cells from 3 published studies: (**a**) **GSE89307** data are for CD8^+^ cells in a murine liver cancer model. (**b**) **GSE84105** and (**c**) **GSE74148** are from mouse chronic infection models, in which CXCR5^+^ cells were previously identified as being reversibly exhausted, while TIM3^+^ cells were identified as being irreversibly exhausted. For clarity, genes primarily regulated post-transcriptionally in TCE are not shown. The heatmap color scale represents per gene log2 expression Z-scores truncated at +/-1. Each heatmap shows 3 groups of genes, as indicated by the color bar at the left of the heatmaps. Genes marked ‘IR’ (gray) represent immune/inhibitory receptors. Genes marked ‘PP’ (green) have high expression early on following stimulation, and are associated with proliferative and memory-precursor states. Genes marked ‘EE’ (purple) have high expression at later CD8^+^ T cell activation periods, and are associated with effector function and irreversible exhaustion. Note the similarities between Tumor Day 5 and Day 7 CD8^+^ cells, and between these states and CXCR5^+^ cells in chronic infection, as well as similarities between Tumor Day 14 and Day 21 CD8^+^ T cells and TIM3^+^ and committed effector cells in chronic infection.

Importantly, the early stages of T cell activation and exhaustion are not steady states (i.e. self-maintaining), but require continued stimulation by antigen and associated co-stimulatory signals (hereinafter referred to as Ag for brevity). Prolonged stimulation eventually drives T cells into irreversible (steady state) exhaustion, maintained by epigenetic changes^6^. How a single regulatory input (Ag presence / absence) can *reliably* drive the multiple transitory state changes that occur during prolonged stimulation (Fig. 1) is currently poorly understood.

In recent years, a number of groundbreaking studies have identified key regulatory genes and interactions underlying CD8^+^ TCE^7^-^12^. In spite of the differences in model systems and experimental protocols, these studies reveal a common set of regulatory interactions underlying TCE. However, all known genes and gene products whose expression or modification state have been associated with TCE also play a role in normal (acute) T cell activation^13^. We hypothesized that TCE may arise from changes in diverse gene regulatory interactions rather than the dysregulation of a single gene.

Here, we present a manually curated, literature-based, and data-driven network of gene regulatory interactions underpinning TCE in CD8^+^ cells. Using diverse published data, we show that the TCE network accurately captures the gene expression states of CD8^+^ T cells in both chronic infection and tumor settings.

Analysis of functionally distinct network motifs reveals multiple overlapping interactions, implementing overlying sets of functional modules (system building blocks) that appear to reinforce each other’s function, and lead to robust, highly stereotypic cellular state changes following activation. A simple mathematical model derived from the network’s functional modules suggests exhaustion arises because the duration of time that activated CD8^+^ T cells spend in a memory-progenitor state is independent of the duration of stimulation, whereas differentiation is prolonged with stimulation. We use our 2-state TCE network model to predict the phenotypic effects of targeted drugs and provide experimental evidence in support of this approach.

## Results

We initially used manual curation of the literature to establish a network of key molecular interactions that are believed to underlie TCE (Supplementary Table 1 and Supplementary Fig. 2). We then superimposed onto this network expression levels of each gene at various stages of T cell activation and exhaustion from published datasets (see Methods; example visualizations shown in Supplementary Figs. 3-8).

**Fig. 3.**
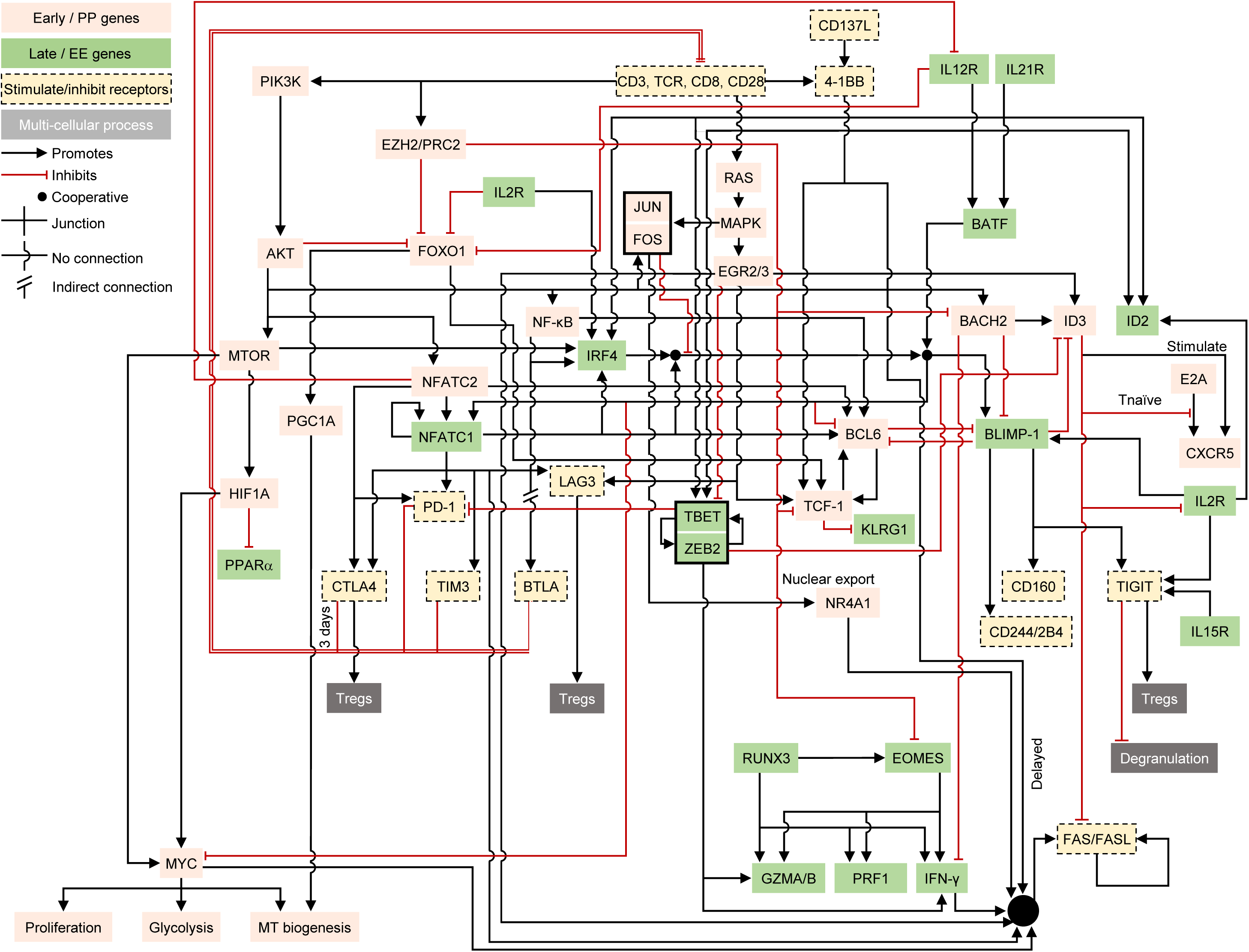
A curated, literature-based network of key regulatory interactions underlying CD8^+^ T cell exhaustion (TCE). Red lines ending in bars indicate inhibition. Black lines ending in arrows indicate activation. Pink network nodes indicate molecules and processes associated with the early pro-memory/proliferative (PP) state. Nodes on a green background represent molecules and processes associated with the late effector/irreversibly exhausted (EE) state. For simplicity, nodes in the network represent both genes and their products and are labeled by their commonly used names across all figures. Inhibitory and related immune receptors are shown on a tan background with dashed borders. In a few cases (e.g. IL2R), a node appears more than once in the network in order to reduce line clutter. The black box around JUN/FOS indicates their cooperative role as AP-1 dimers. Likewise, T-BET and ZEB2 can act cooperatively and are grouped together. The double-edged repressive input onto the T cell receptor (TCR) complex (labeled ‘CD3, TCR, CD8, CD28’) bundles the actions of the inhibitory receptors PD-1, BTLA, CTLA-4, TIM3, and LAG3.

**Fig. 4.**
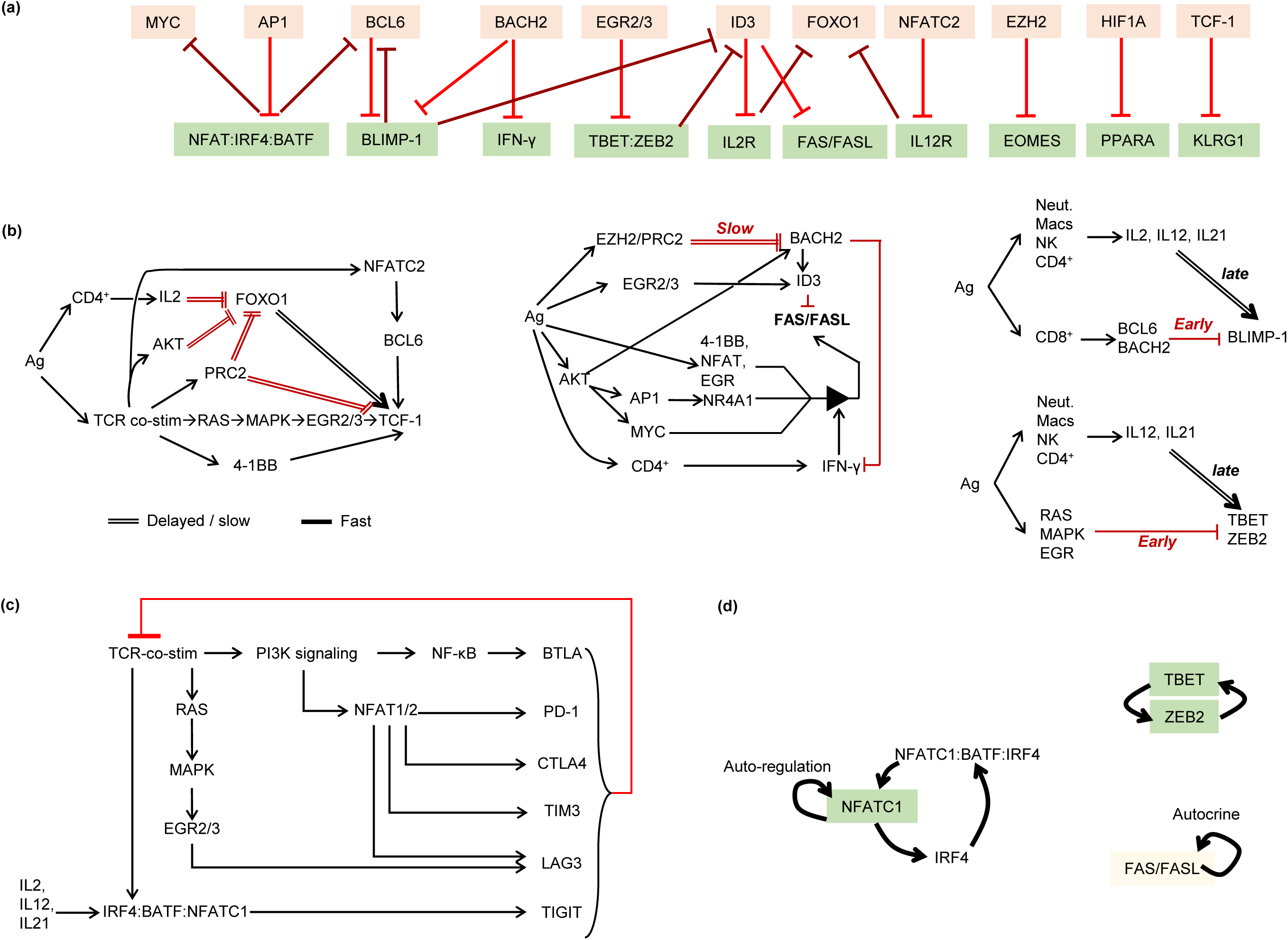
The TCE network can be decomposed and understood as a collection of overlapping functional building blocks. (**a**) There is widespread mutual inhibition between early (PP) and late (EE) state genes, suggesting the 2 states mutually exclude each other. (**b**) The expression of the PP-state driver gene *TCF-1*, the onset of restimulation-induced cell death (RICD) via FAS/FASL signaling, and the time of activation of the key EE state genes *BLIMP-1* (aka *PRDM1*), *TBET*, and *ZEB2* are each controlled by multiple, overlapping incoherent feed-forward loop network motifs. Double lines mark slow/delayed processes. The triangular symbol in the FAS/FASL network indicates that IFN-γ signaling is required for upstream factors such as 4-1BB, NFATs, EGRs, and NR4A1 to activate the transcription of FAS and FASL. (**c**) Inhibitory immune receptors implement overlapping negative feedback inhibition of T cell activation. It should be noted that CTLA4, LAG3 and TIGIT additionally exert inhibitory activity via regulatory T cells not shown here to emphasize the negative feedback on CD8/TCR signaling exerted by inhibitory receptors. (**d**) Multiple positive feedback loops reinforce and maintain the late effector/irreversible exhaustion (EE) state.

**Fig. 5.**
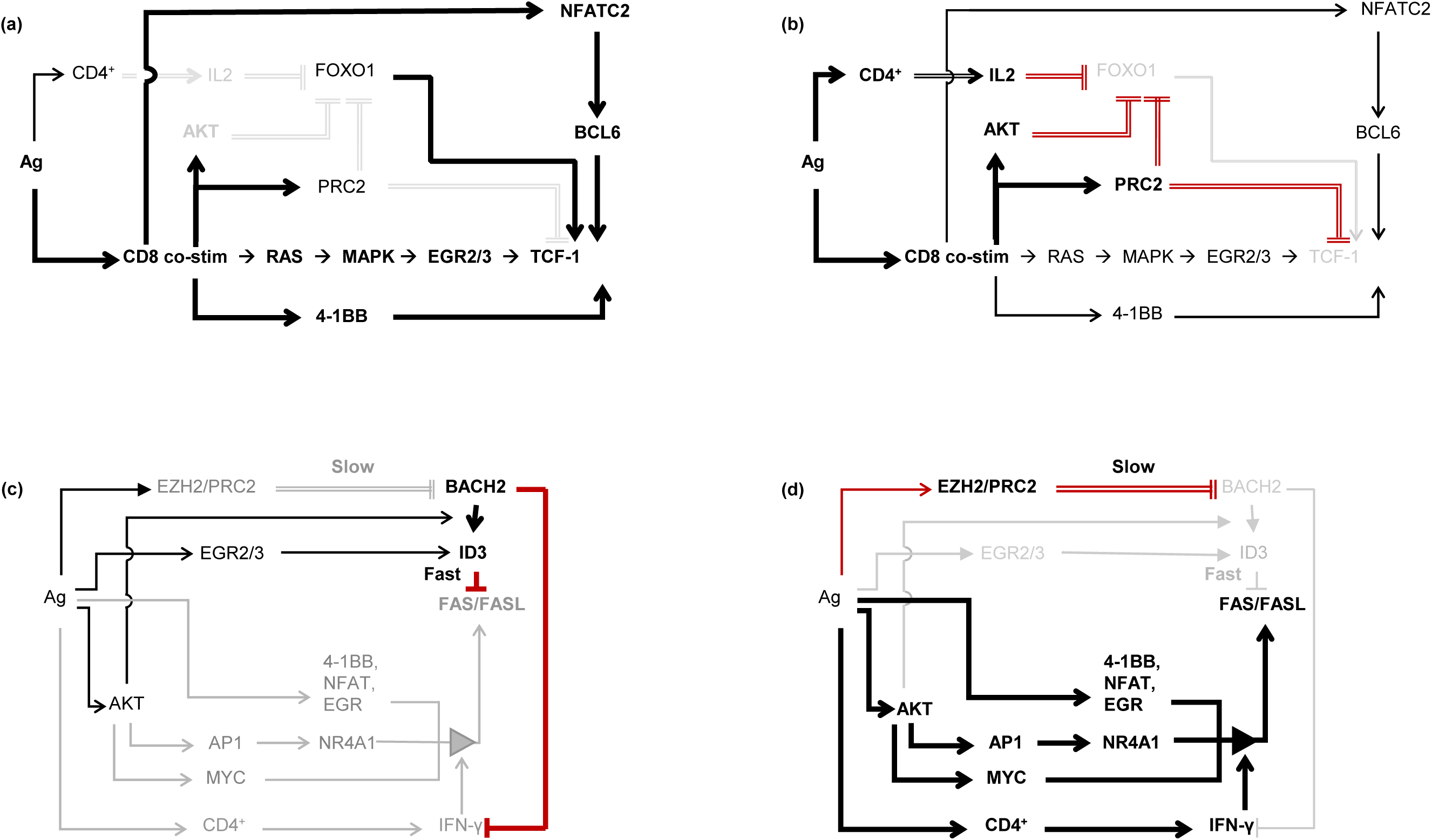
Two-state models of the incoherent feed-forward loops (iFFLs) regulating TCF-1 and FAS/FASL activity. (**a**) The early state of the TCF-1 iFFL. Thick black lines indicate active regulatory interactions. Light gray lines and text indicate inactive interactions and genes. TCF-1 and FOXO1 are expressed in naïve CD8^+^ T cells. TCF-1 expression during early CD8^+^ T cell activation is maintained via direct regulation by FOXO1, and via MAPK signaling activation of ERG2/3, via 4-1BB signaling, and through calcium-activated NFATC2. The 2 repressive regulators of TCF-1, EZH2/PRC2 signaling and PI3K/AKT signaling, are inactive at this stage. AKT activation in stimulated CD8^+^ cells peaks at ∼day 5 post-infection^52^. EZH2/PRC2 activity peaks at around 4 days post-infection^30^. (**b**) At around 3 days post-infection, AKT represses FOXO1 activity, while PRC2 initiates a chain of events culminating in DNA-methylation and transcriptional shut-down of *FOXO1* and *TCF-1* genes. IL2 signaling also becomes active around this time and suppresses FOXO1 activity. Thin black lines indicate activating inputs overridden by dominant inhibitory inputs. (**c**) and (**d**) illustrate the early and late states of the FAS/FASL signaling iFFL using the same notation as in (**a**) and (**b**). In (**c**), BACH2 and ID3 are active in naïve and early activated CD8^+^ cells, and block FAS/FASL expression and signaling. At later time points (**d**), EZH2/PRC2 activity suppresses BACH2/ID3. IFN-γ signaling becomes activated, and facilitates activation of FAS/FASL by 4-1BB, NFATs, EGRs, and NR4A1.

**Fig. 6.**
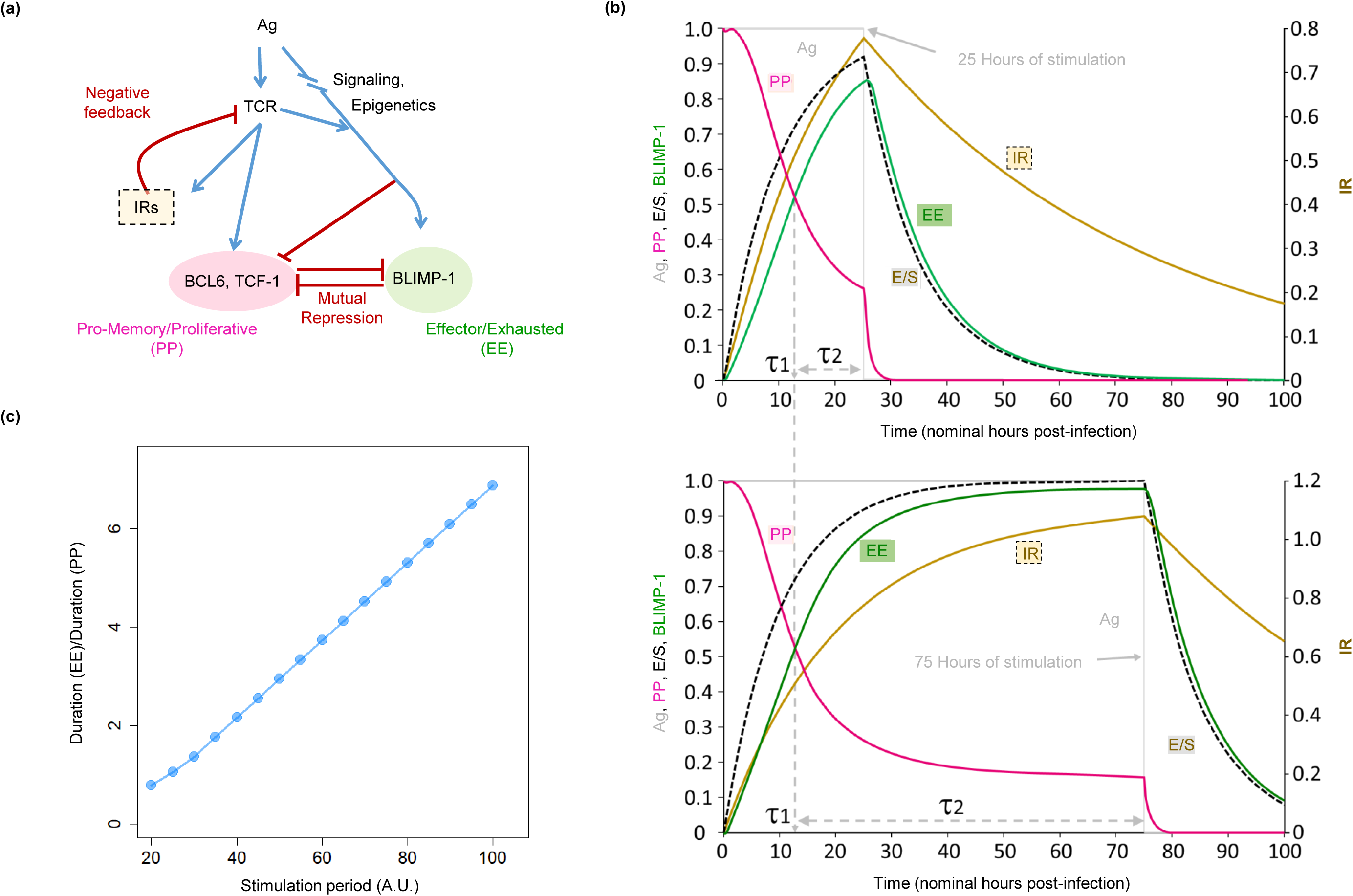
Functional building block-based modeling reveals a mechanism driving T cell exhaustion (TCE). (**a**) Abstract representation of the overlapping incoherent feed-forward loops (iFFLs) that regulate the timing of expression/repression of key PP and EE state genes *TCF-1/BCL6* and *BLIMP-1*. The 2 states are modeled as being mutually repressive (cf. Fig. 4a). Additionally, inhibitory immune receptors (IRs) exert negative feedback on T cell receptor (TCR) activity. The overlapping iFFLs create a brief period of fixed length shortly after TCR stimulation, during which PP state driver genes (*TCF-1* and *BCL6*) are active. At the end of this period, PP state genes become repressed, while EE state genes such as *BLIMP-1* are simultaneously activated. A set of ordinary differential equations (ODEs) implementing this model are described in Methods. (**b**) Simulation results for the model in (**a**) under nominal ‘acute’ and ‘chronic’ stimulation conditions. The gray line shows the duration of antigen availability (nominally 25 hours for the ‘acute’ model and 75 hours for the ‘chronic’ model). The time at which the red PP-activity curve crosses the green EE-activity curve (τ1) marks the duration of time the cells spend in the PP state. The period from the end of τ1 until antigen clearance (labeled τ2) marks the duration cells spend in the effector/exhausted EE state. Comparison of the 2 simulation runs shows that τ1 remains constant irrespective of the duration of stimulation, while τ2 increases with the duration of stimulation. (**c**) The ratio (τ2 / τ1) as a function of stimulation period grows in approximate proportion to the total duration of stimulation, suggesting that the proportion of cells in the EE state will increase with prolonged stimulation.

**Fig. 7.**
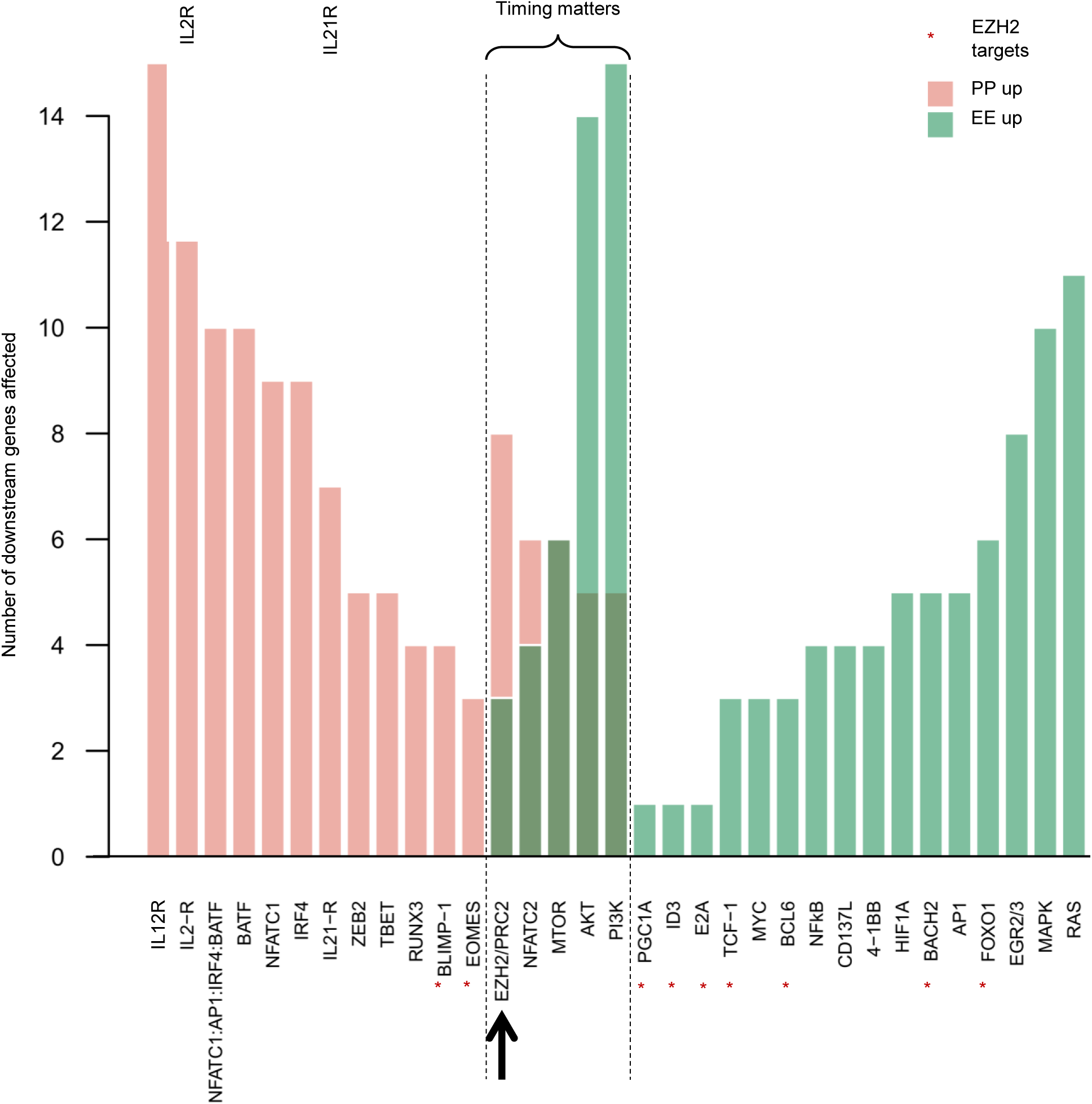
Model-based prediction of the effects of interfering with the activities of genes in the TCE network. For each gene listed, bar heights show the number of downstream genes that are affected by perturbation of the regulator gene. Affected gene counts are grouped into pro-PP (PP gene up-regulated or EE gene down-regulated) and pro-EE effects (EE gene up-regulated or PP gene down-regulated). Leaf nodes (i.e. genes with no downstream targets in the network), and genes within global feedback paths (e.g. the checkpoint receptors) are excluded. Of note, inhibiting the activity of a large majority of the genes in the TCE network results in exclusively pro-PP or pro-EE effects. But a number of genes, exemplified by *EZH2*, appear to have both pro-PP and pro-EE effects (as an example, EZH2 targets are marked by *). As illustrated in Fig. 8a-c, these apparently contradictory effects are resolved when the activities of these genes are explored at finer time-resolutions.

**Fig. 8.**
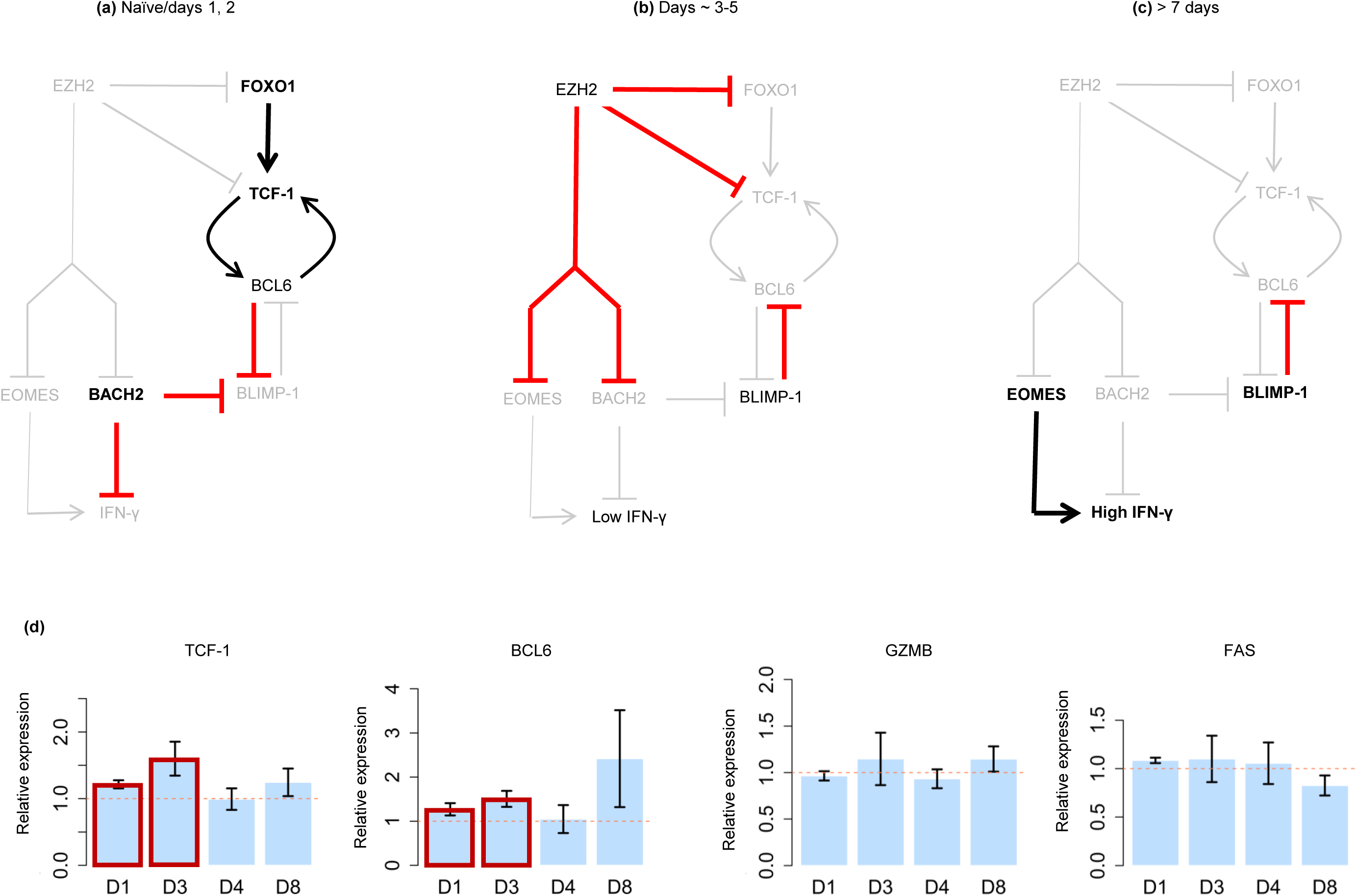
Prediction of the effect of EZH2 inhibition on T cell exhaustion (TCE). The networks in panels (**a-c**) show 3 distinct states that the primary targets of EZH2/PRC2 adopt following TCR stimulation, and reveal that EZH2 has pro-PP effects at all times. Although *EZH2* transcription peaks at about 1 day post-stimulation, its repressive effects (via PRC2) are not apparent until around Day 3 post-infection. PRC2 is not active in naïve CD8^+^ cells, or during the earliest days post-stimulation. During this time, FOXO1 drives TCF-1 (pro-PP state) expression, and BACH2 represses the pro-EE state gene *BLIMP-1*. (**b**) At around 3-5 days post-infection, EZH2/PRC2 inhibit FOXO1, BACH2, and TCF-1, allowing the expression of the pro-EE gene *BLIMP-1*. In turn, BLIMP-1 represses the pro-PP gene *BCL6*. Concurrent with these events, EZH2 inhibition blocks early expression of the EE gene *EOMES*. Thus, inhibition of EZH2 during this period will block/delay the onset of the EE state. (**c**) *EZH2* transcription returns to basal levels by about 7 days post-infection, allowing EOMES to become activated and drive high (EE state) IFN-γ expression. (**d**) Experimental evidence supporting the pro-PP predicted effect of EZH2 inhibition in human peripheral blood T cells (average of 3 donors and 3 technical replicates per donor). Shown are the relative mRNA levels (treatment relative to no-treatment) of the pro-PP genes *TCF-1* and *BCL6*, and the EE marker genes *GZMB* and *FAS* as measured by RT qPCR at days 1, 3, 4, and 8 post-infection (D1 to D8). As predicted, both TCF-1 and BCL6 are up-regulated in the early days post-infection (red boxes), while GZMB and FAS expression levels show no significant change. Error bars mark ± 1 standard error.

Next, we calculated fold changes in expression compared to the naïve state and evaluated the fraction of times each interaction exhibited fold changes in source and target genes consistent with the reported sense of the interaction (promoting or inhibitory). At a false discovery rate of ∼15% (estimated via permutation testing), only 17 interactions (∼3.6%) in the network did not have supportive expression data (see Methods and Supplementary Fig. 9).

Consistent with the observation that all driver genes implicated in TCE are common to, and play similar roles in, exhausted and acute responses, principal component analysis (PCA) of expression data for the genes in the TCE network exhibited similar trajectories for acute and exhausted gene expression profiles. TCE gene sets from previous studies also showed similar trajectories and further support our findings (Supplementary Fig. 10).

In addition to known regulators of TCR stimulation and co-receptor engagement, T cells subjected to repeat stimulation and exhaustion undergo multiple metabolic changes^14,15^. Indeed, metabolic deprivation can ameliorate T cell activation^16^. Consistent with our TCE network findings, analysis of gene expression changes driving metabolic switching during acute and chronic stimulation also suggested similar expression state trajectories for CD8^+^ cells undergoing acute and chronic stimulation (Supplementary Figs. 11 and 12).

Considering that the genes involved in both acute and chronic stimulation are largely the same (as revealed by the literature-based network), and that the trajectories of their expression changes are also similar (as revealed by our gene expression and metabolic analyses), we hypothesized that the differences between acute and chronic T cell responses may lie in the timing of gene expression changes.

To find differences in the relative timing of gene expression changes between acute and chronic responses, we carried out time course gene expression clustering (Supplementary Figs. 13-15), and looked for transcription factors and signaling genes that switch cluster membership (i.e. timing) between acute and chronic settings. This approach identified metabolic changes downstream of *TP53* as differentially regulated in acute and chronic settings. Of the three *TP53* regulatory genes whose expression changes correlated with the TCE network genes, only one gene (CD200-R1, a known T cell suppressor^17-19^) showed consistently large fold change differences between chronic and acute settings in multiple datasets (Supplementary Fig. 16). However, recent research suggests CD200-R1 activity inhibits T cell responses primarily through CD8^+^ T cell independent pathways^20-22^.

### Network analysis

In contrast to the above findings, a series of alternate logic models of the TCE network (see Methods) all required specific gene activation delays in order to recapitulate the observed sequences of CD8^+^ cell gene expression changes during acute and chronic stimulation (see Supplementary Figs. 17-20 and Methods for examples). Feed-forward and feedback loops (Supplementary Fig. 21) are widely used to control timing and activity in both biological and engineered systems^23,24^. Thus, to address how antigen availability can determine a sequence of specifically timed gene expression changes, we searched for feed-forward and feedback motifs in our literature-derived TCE network.

To facilitate loop detection, the initial literature-based TCE network was simplified by collapsing all isoforms of each gene into a single gene symbol (e.g. *NF-*κ*B* instead of *RELB, cREL*, etc.) unless a specific isoform was known to play a distinct role in TCE. Similarly, chains of interactions with no incoming or outgoing branches were collapsed into single nodes, and (where possible), members of protein complexes were grouped into single nodes. Finally, genes with no reported downstream targets and unknown regulatory significance were removed. To maximize loop detection, we then carried out a second, more detailed literature review to identify all known interactions of the genes in our reduced literature-based TCE network. The resulting network has 64 nodes and 120 interactions (Supplementary Fig. 22, and Supplementary Table 2).

### Genes in the TCE network can be divided into early and late activity classes

Multiple recent studies of TCE have noted the existence of distinct ‘reversible’ and ‘irreversible’ subpopulations of exhausted CD8^+^ T cells^10-12^. Reversibly exhausted CD8^+^ T cells have been variously defined by high expression levels of *CXCR5, TCF-1*, and *BCL6*, and concurrent low expression of *KLRG1, BLIMP-1 (PRDM1)*, and *TIM3 (HAVCR2)*^25^. They have relatively high proliferative potential and can produce fully functional memory cells^25^. In contrast, irreversibly exhausted CD8^+^ T cells are defined by the opposite expression pattern and have low proliferative and memory-forming potential.

While performing expression clustering and time course analysis of the TCE network genes, we noticed that markers of ‘reversible exhaustion’ (e.g. high *CXCR5* in LCMV infections, high *TCF-1* in tumor-infiltrating lymphocytes, accompanied with low levels of *KLRG1* and *TIM-3*) overlap with and include pro-memory genes, and exclude genes associated with effector function and ‘irreversible exhaustion’ (Fig. 2). Moreover, many genes associated with the reversible exhaustion state are also associated with proliferation (notably *TCF-1, MYC, NF-*κ*B*, and genes enabling glycolysis). These findings led us to hypothesize that the processes underlying CD8^+^ T cell activation and exhaustion may fall into 2 broad classes: pro-memory and proliferative, versus pro-effector and exhaustion.

Using clustered gene expression time course profiles (Supplementary Figs. 13-15) and consistent with literature reports, we were able to assign the regulatory interactions underlying TCE into 2 groups defining an early, pro-memory and proliferative state (PP), and a later state associated with effector differentiation and function ultimately leading to irreversible exhaustion (EE). The resulting network model is illustrated in Fig. 3. It is important to note that in terms of nodes (i.e. genes/gene products) and interactions, the network in Fig. 3 is identical to that in Supplementary Fig. 22 (i.e. both are visualization of the citations listed in Supplementary Table 2). The key differences are that the layout and coloring of the network have been manually adjusted to highlight the PP and EE network components and their interactions.

### Network motifs identify modular functional building blocks

We next used the 2-state TCE network of Fig. 3 to search for regulatory network motifs and functional building blocks that may explain the state changes of CD8^+^ cells undergoing TCE. A summary of the functional network motifs/building blocks discovered is presented in Fig. 4.

#### Mutual inhibition between early and late activation states

Excluding the negative-feedback interactions of the inhibitory receptors (discussed below), a remarkable 18 out of 23 inhibitory interactions in our TCE network (78%) are between ‘early’ (PP) and ‘late’ (EE) activation genes (Fig. 4a). Such inhibition can enable the mutual exclusion of these 2 essential activation states in either a graded or bistable manner^24^. Consistent with this hypothesis, recently published single-cell RNA-seq data^8^ suggest *BCL6* and *BLIMP-1* repress each other in a mutually exclusive manner (Supplementary Fig. 23).

#### Overlapping incoherent feed-forward loops fix the duration of the pro-memory PP state

In an incoherent feed-forward loop (iFFL), an upstream regulator activates and represses a downstream target via pathways that operate on different timescales^26^. One characteristic behavior of such regulation is that the downstream target will be turned on (or off) for a fixed duration corresponding to the difference in the timescales of the activating and inhibitory pathways.

As summarized in Fig. 4b, a set of overlapping iFFLs ensure delayed loss of nuclear FOXO1 protein activity, and concomitant loss of *TCF-1* gene expression. FOXO1 and TCF-1 are both expressed in naïve and memory T cells^27,28^. Following T cell stimulation, their expression is abrogated by delayed Polycomb repressor complex 2 (PRC2)-mediated inhibition downstream of TCR signaling (see also Supplementary Table 3).^29,30^. An additional set of iFFLs involving repression of FOXO1 by signaling downstream of the IL2 and IL12 receptors further reinforces delayed repression of TCF-1. Taking these observations together, we hypothesize that the duration of *FOXO1/TCF-1* transcription following stimulation is fixed by the time it takes for TCR-activated PRC2 and signaling pathways to silence the 2 genes. The sequences of early and late regulatory interactions governing *TCF-1* expression are illustrated schematically in Fig. 5a, b.

Focusing on interactions among the ‘early’ genes and cell surface receptors in the TCE network revealed an additional iFFL that regulates the timing of FAS/FASL signaling (Fig. 4b). Interferon gamma (IFN-γ) signaling, which is required for FAS/FASL activity^31^, is repressed early on by BACH2^32^, which is expressed in naïve CD8^+^ T cells^33^ and activated by TCR-mediated signaling (Fig. 5c). At later time points, TCR-activated PRC2 epigenetically represses *BACH2* expression^29^, thus enabling FAS/FASL activation by IFN-γ (Fig. 5d).

A third set of overlapping iFFLs ensure the delayed activation of the late/effector state genes *TBET, ZEB2* and *BLIMP-1* (Fig. 4b). Thus, the duration of activity of early/progenitor state (PP) genes, the timing of the activation of the late effector/exhausted state (EE) genes, and the timing of FAS/FASL signaling to limit proliferation, are all controlled by overlapping iFFLs that share many genes and reinforce each other’s function in a coordinated, redundantly robust manner.

With the exception of *TCF-1/BCL6* mutual activation, the regulatory interactions of the early-phase genes in the TCE network are mediated entirely by iFFLs. We hypothesize that these iFFLs create a fixed duration time window during which activated T cells proliferate, are in a memory precursor state, and are capable of reinvigoration.

#### Negative feedback by inhibitory receptors

Negative feedback (Supplementary Fig. 21 and Fig. 4c) is a well described mechanism for homeostasis. At least 2 distinct sets of negative feedbacks appear to regulate TCE. Firstly, negative feedback via FAS/FASL signaling results in restimulation-induced cell death^34^ (RICD, Fig. 1) and is thought to limit T cell numbers following activation-induced proliferation^34^.

In addition, and in contrast to population control via FAS-mediated RICD, negative feedback via inhibitory receptors primarily acts by down-regulating signal transduction downstream of the TCR^35-40^, and can guard against over-activation of individual T cells. In this context, negative feedback can allow a vigorous early response that is actively down-regulated at later times to avoid over-reacting^23^ (Supplementary Figs. 24 and 25).

#### Positive feedback loops maintain the late EE state

In contrast to the early PP state, the later EE state has no iFFLs and is instead self-reinforced via multiple positive feedback loops (Fig. 4d). Importantly, such feedback loops stabilize the expression of not only the nodes directly involved in each loop (NFATC1, IRF4, TBET, ZEB2, and BLIMP-1), but also their regulatory targets, which include CTLA4, PD-1, TIM3, LAG3, IFN-γ, GZMA/B, PRF1, FAS/FASL, 2B4, CD160, and TIGIT.

### Known interactions are sufficient to explain T cell exhaustion

As noted above (Figs. 4 and 5), the onset of the late EE state leads to the repression of key drivers of the early PP state. Moreover, the transition between the 2 states occurs at a fixed time post-stimulation dictated by delayed/slow regulatory interactions within iFFLs, and independent of the duration of stimulation. Thus, the longer that T cell stimulation continues, the more time a given CD8^+^ T cell will spend reinforcing its EE state at the cost of pro-memory and proliferative states. Fig. 6 presents simulation results illustrating this principle (see Methods for details).

### The 2-state TCE model allows prediction of drug effects

To identify optimal targets for counteracting exhaustion, we first computed all downstream targets of each gene in the TCE network (see Methods for details). Depending on whether an interaction is inhibitory or activating, lowered activity of an upstream regulator will increase or decrease the activity of target genes. To estimate the overall phenotypic effect of an upstream gene, we calculated a pro-PP score by adding the number of up-regulated PP genes to the number of down-regulated EE genes, and a complementary pro-EE score.

As shown in Fig. 7, blocking or reducing the activity of most genes in the TCE network results in exclusively pro-PP or pro-EE predicted effects. A small number of genes appear to impact both PP and EE states because they play distinct pro-PP or pro-EE roles at different times post-stimulation. For example, transcription of *EZH2* (the core subunit of PRC2) is up-regulated following TCR co-stimulation, peaks at ∼1 day post-infection, and is largely back to basal levels by day 7 post-infection^30^. At the protein level, PRC2 activity peaks around 3-5 days post-infection^41,42^, at which time EZH2/PRC2 suppress the already active pro-memory/proliferation genes *FOXO1* and *TCF-1* while suppressing the pro-effector/exhaustion gene EOMES (Fig. 7 and Fig. 8a-c). Thus, based on the network topology alone, it appears as though EZH2 represses both the PP and EE network states, but time step analysis resolves the apparent contradiction and suggests reduction of EZH2 activity should have exclusively pro-PP effects (because EZH2 is inactive at the times when it could have pro-EE effects).

To test the hypothesis that EZH2 inhibition will increase the early expression of PP state genes without impacting EE state genes, we stimulated blood-derived T cells from 3 volunteers with equal quantities of CD3 and CD28 antibodies in vitro for 5 days with and without drug-mediated EZH2 inhibition (see Methods for details). As shown in Fig. 8d, quantitative RT-PCR revealed up-regulation of the pro-PP state genes *TCF-1* and *BCL6* in EZH2-inhibited samples, while the EE state markers *FAS* and *GZMB* showed no significant change.

## Discussion

The network analysis presented here suggests CD8^+^ TCE arises from an inherent topological property of the network of regulatory interactions that underlie CD8^+^ T cell responses to both acute and prolonged stimulation. Specifically, we presented computational and experimental evidence suggesting that the duration of time CD8^+^ T cells spend in a pro-memory state is fixed by network properties that limit the duration of activity of the pro-memory gene *TCF-1* in a manner independent of the duration of stimulation.

A recently published study used mass cytometry to classify exhausted CD8^+^ T cells into multiple subtypes^43^. Consistent with our model, Bengsch et al.^43^ report that in HIV infections with a lower viral load and in HIV patients on anti-retroviral therapy (i.e. cases with greater immune activity), exhausted T cells have higher expression levels of TCF-1 and/or CXCR5, and appear more functional. Further supporting our model, drug-induced activation of TCF-1 has been shown to block CD8^+^ T cell differentiation, and increase proliferation^44^.

The analyses presented here focused on molecules and interactions generally accepted by the research community to play important roles in CD8^+^ T cell responses to acute and chronic stimulation. It is important to note that the approach presented here can be easily extended to predict novel molecules and interactions by extending our literature-based network to include known and predicted protein-protein and protein-DNA interactors of the current network nodes.

Analysis of our literature-based, data-driven, and manually curated TCE network suggests TCE is a highly robust process mediated by multiple redundant and overlapping feedback and feed-forward loops. In addition, diverse, overlapping molecular mechanisms contribute to TCE, including combinatorial regulation by transcription factors, post-translational modifications, protein localization, chromatin state changes, and metabolic reprogramming. Taken together, our analyses are consistent with the view that TCE is not a dysfunctional state, but rather an adaptive response to situations where the immune system fails to clear antigen rapidly. Clinical remediation of TCE will need to overcome multiple, overlapping and redundant regulatory mechanisms, requiring combinatorial interventions.

## Methods

### Datasets

The following published datasets were downloaded from the NCBI Gene Expression Omnibus (GEO) repository (https://www.ncbi.nlm.nih.gov/geo/). The “GSE” codes given below are the unique IDs for each dataset. The relevant subsets of the conditions used in this study are indicated individually below. Raw microarray data were normalized using the ‘normalizeBetweenArrays’ function of the R/Bioconductor ‘limma’ package (https://bioconductor.org/packages/release/bioc/html/limma.html). All microarray analyses are based on probes with the highest interquartile range (IQR).

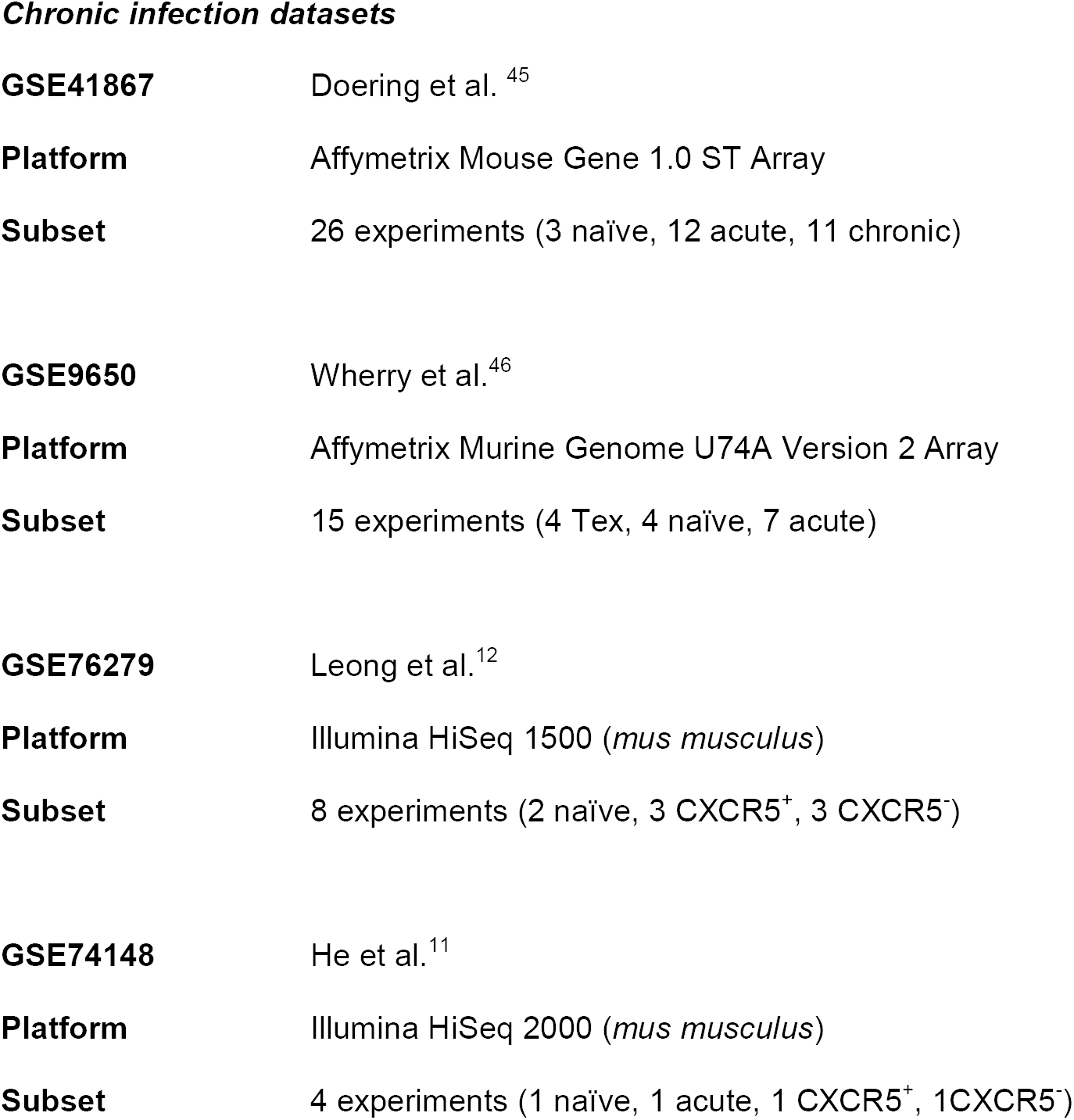
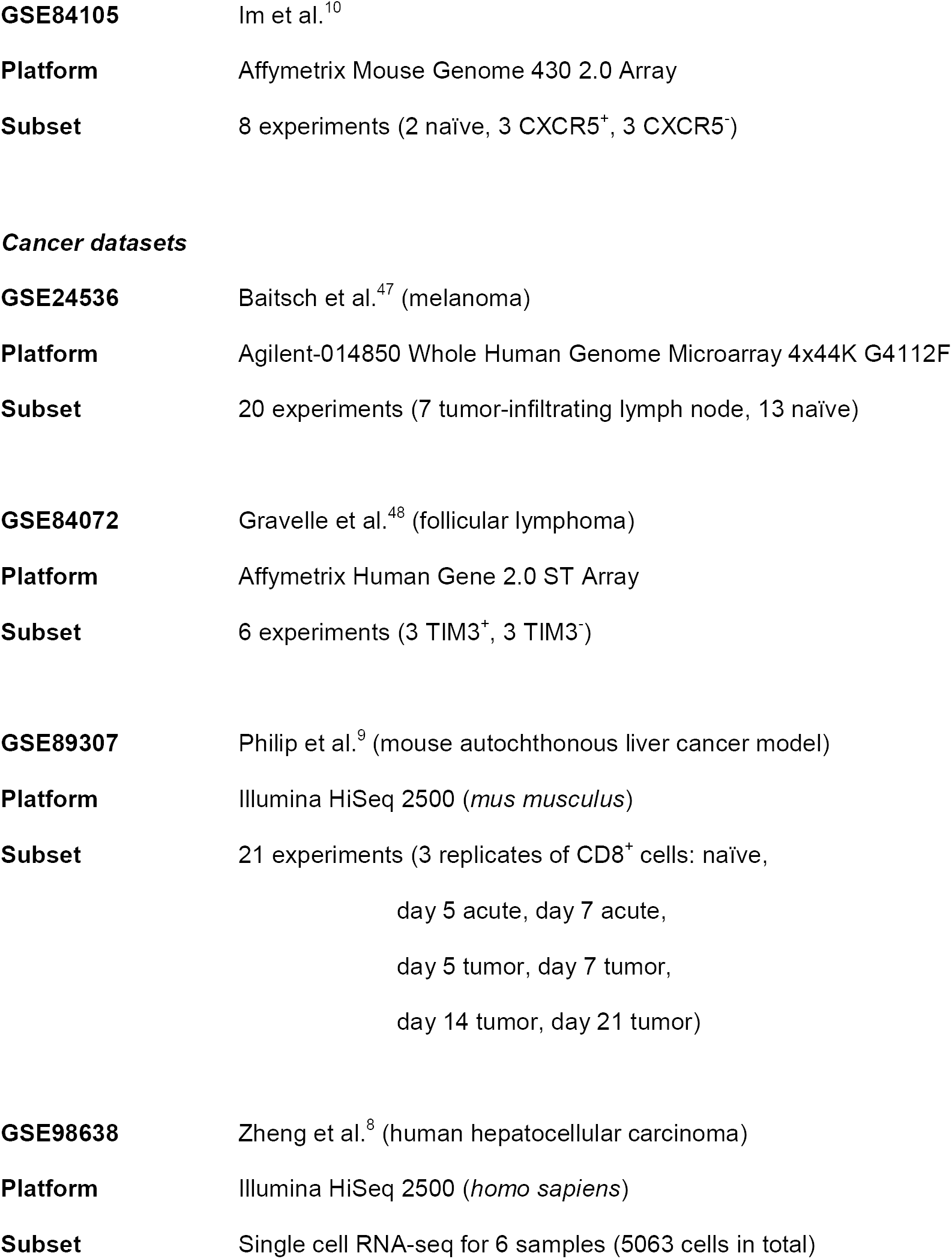
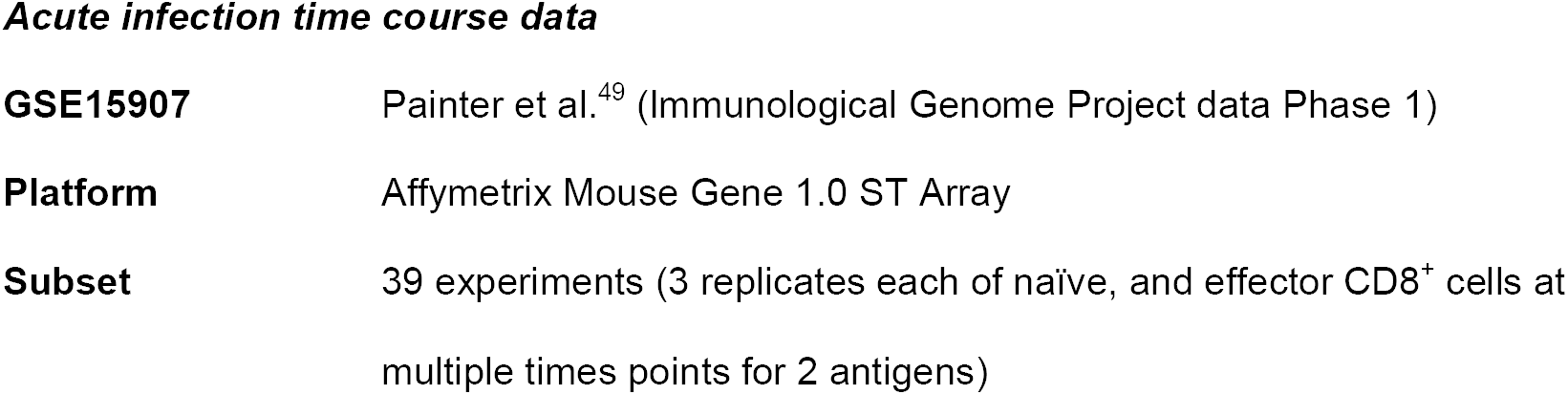

### Construction of the literature-based TCE network

The large-scale network of regulatory interactions underlying CD8^+^ TCE was manually curated from reports in publications released before August 2017, focusing on genes reported by multiple publications, as well as genes with small-scale (non-‘omics’) experimental support in a single paper. For brevity, only key publications are cited. In situations where different transcripts or protein isoforms may be involved, all isoforms were included with a view to evaluating them using expression data.

The 64-node reduced/simplified TCE network was extracted and curated manually by one co-author, and then independently verified on an interaction-by-interaction and citation-by-citation level by a second co-author.

### Network visualization

Both the large-scale TCE network, and its simplified version were visualized using the RCyJS (http://bioconductor.org/packages/release/bioc/html/RCyjs.html).

Expression data were superimposed onto the large-scale TCE network using RcyJS. Fold changes were calculated with respect to naïve CD8^+^ T cells and mapped to a color-scale as shown. Edge (line) thicknesses are proportional to the fraction of observations in which the source and target genes in an edge have fold changes concordant with the sense of the interaction (‘promoting’ or ‘inhibiting’). Thus, the auto-regulatory loop of NFATC1 – by definition – has maximum thickness. In some instances, edges were highly concordant (good agreement between the network model and data) in one dataset or condition, and not in another.

### Network evaluation by concordance matching

For each edge (interaction) in the TCE network, the fraction of times when the source and target genes in an edge had fold changes concordant with the sense of the interaction (‘promoting’ or ‘inhibiting’) were calculated. To account for variability in the data, fold changes were calculated *per replicate* data (instead of averaging data across all replicates). Thus, data for 2 replicates across 2 conditions generated 4 possible comparisons with potential average concordance values of (0.00, 0.25, 0.50, 0.75, 1.00).

To explore the statistical significance of the observed edge concordance values, 500,000 edges were generated with randomly assigned gene labels and interaction types. Next, the fraction of times an edge was concordant in a given dataset for the TCE and randomized (control) networks was calculated. Concordant edges occurred by chance in 14.8% of randomized controls, suggesting a false discovery rate of ∼15%.

### State tracking by PCA

To compare the TCE network state changes among different CD8^+^ T cell subsets, PCA was performed for each dataset using the normalized expression levels of all the genes in the TCE network. The relative positions of any 2 CD8^+^ cell subtypes on a plot of the first 2 components reflect their similarities/differences in expression of the TCE network genes and show similar state trajectories for responses to acute and chronic stimulation.

### Evaluation of cellular metabolic activity using transcriptional signatures

A set of marker regulatory genes per pathway were manually derived to explore metabolic changes in activated and exhausted CD8^+^ cells. Metabolic pathways typically comprise 3 topological features: metabolic transformation chains, incoming (joining) paths, and outgoing (forking) paths. We focused on marker genes that regulate metabolic transformation chains, because such genes are more likely to be highly correlated.

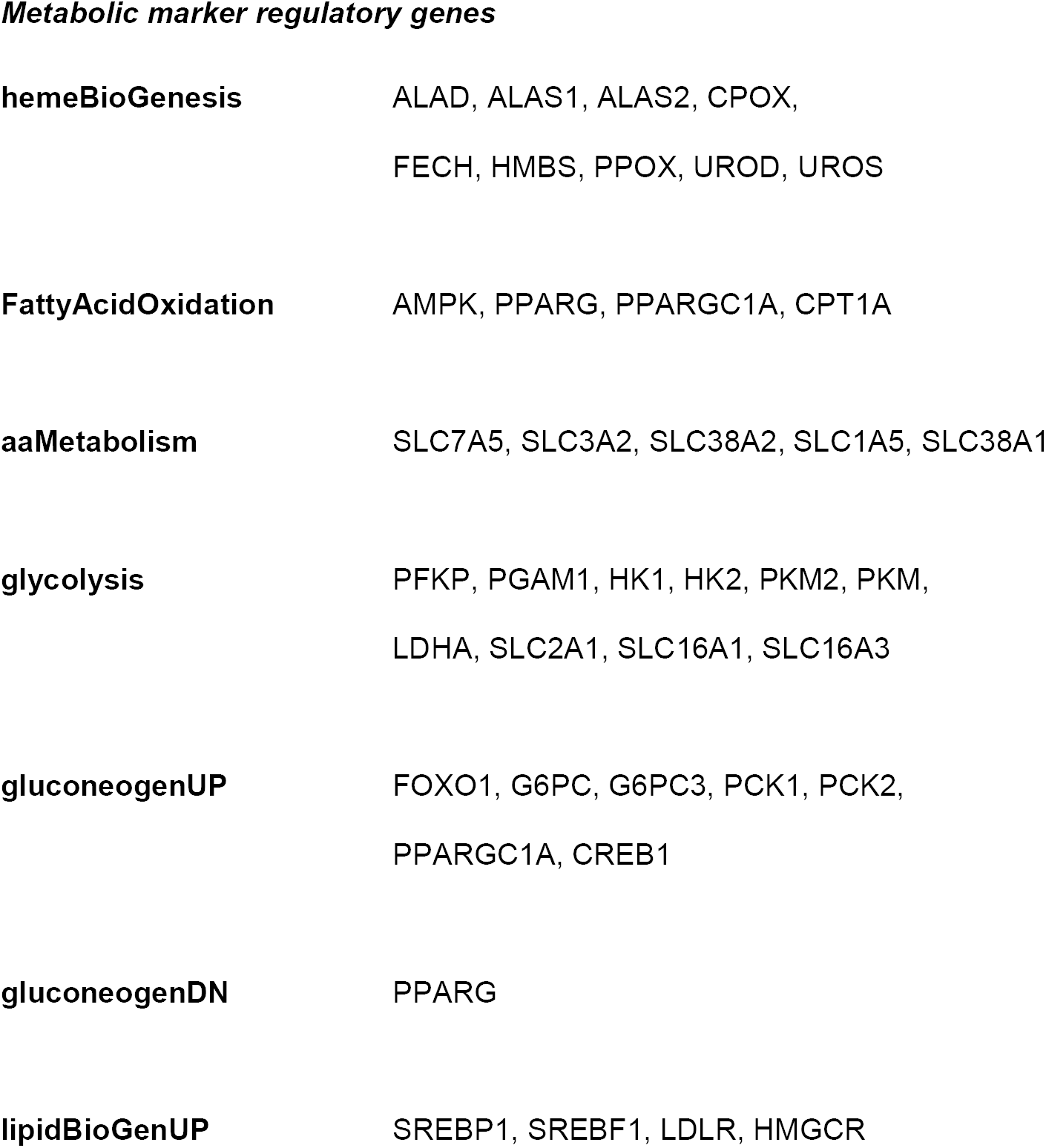
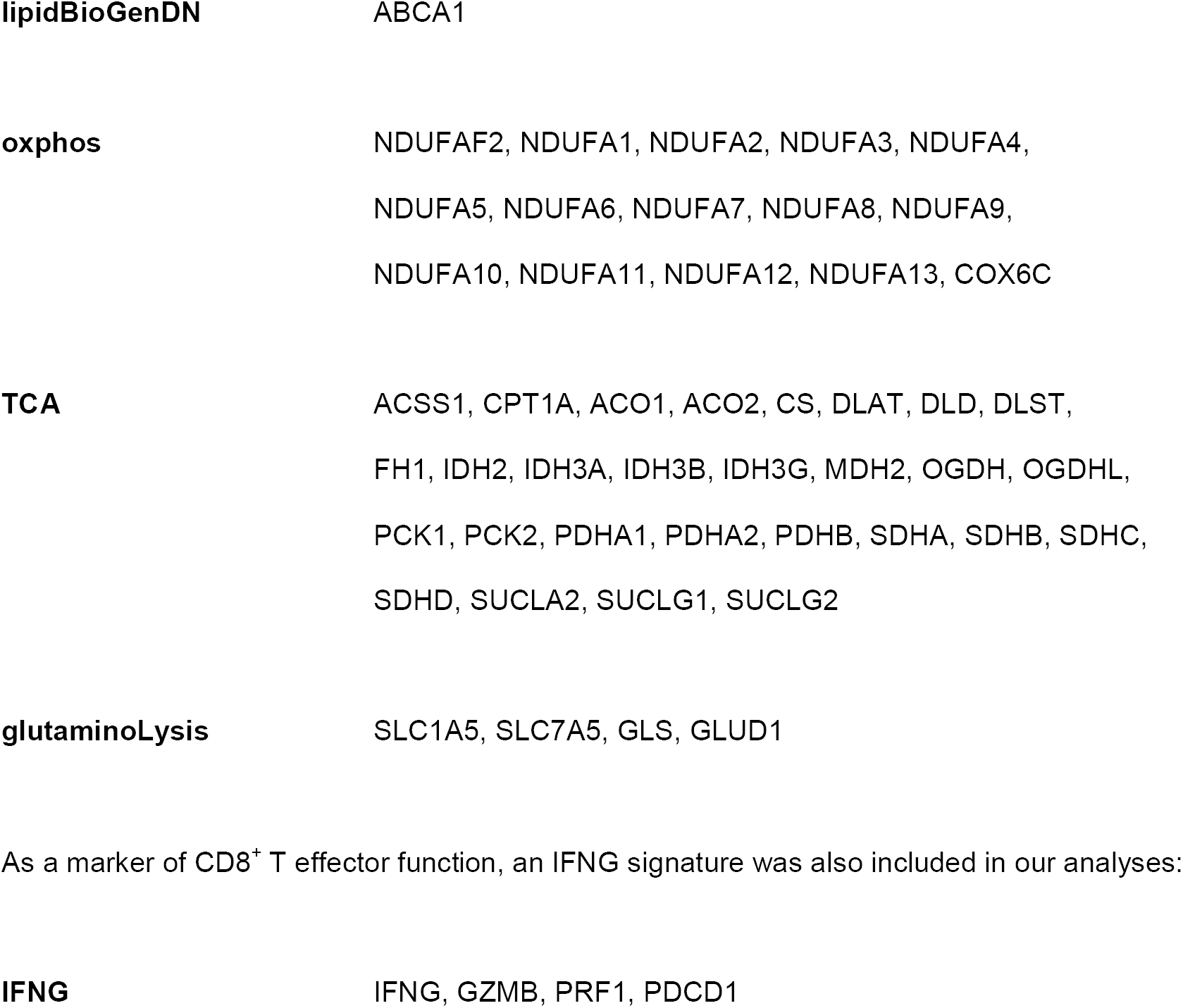

*Metabolic marker regulatory genes*
|  |  |
| --- | --- |
| **hemeBioGenesis** | ALAD, ALAS1, ALAS2, CPOX,<br>FECH, HMBS, PPOX, UROD, UROS |
| **FattyAcidOxidation** | AMPK, PPARG, PPARGC1A, CPT1A |
| **aaMetabolism** | SLC7A5, SLC3A2, SLC38A2, SLC1A5, SLC38A1 |
| **glycolysis** | PFKP, PGAM1, HK1, HK2, PKM2, PKM,<br>LDHA, SLC2A1, SLC16A1, SLC16A3 |
| **gluconeogenUP** | FOXO1, G6PC, G6PC3, PCK1, PCK2,<br>PPARGC1A, CREB1 |
| **gluconeogenDN** | PPARG |
| **lipidBioGenUP** | SREBP1, SREBF1, LDLR, HMGCR |
| **lipidBioGenDN** | ABCA1 |
| **oxphos** | NDUFAF2, NDUFA1, NDUFA2, NDUFA3, NDUFA4,<br>NDUFA5, NDUFA6, NDUFA7, NDUFA8, NDUFA9,<br>NDUFA10, NDUFA11, NDUFA12, NDUFA13, COX6C |
| **TCA** | ACSS1, CPT1A, ACO1, ACO2, CS, DLAT, DLD, DLST,<br>FH1, IDH2, IDH3A, IDH3B, IDH3G, MDH2, OGDH, OGDHL,<br>PCK1, PCK2, PDHA1, PDHA2, PDHB, SDHA, SDHB, SDHC,<br>SDHD, SUCLA2, SUCLG1, SUCLG2 |
| **glutaminoLysis** | SLC1A5, SLC7A5, GLS, GLUD1 |
| As a marker of CD8<sup>+</sup> T effector function, an IFNG signature was also included in our analyses: |  |
| **IFNG** | IFNG, GZMB, PRF1, PDCD1 |

### Calculating metabolic pathway signatures

Expression data were first averaged per gene across replicates, and then averaged across marker genes per pathway. For pathways with a known repressor, the average expression score was normalized by the average expression of the known repressors. To facilitate visual comparisons across datasets, all activity scores were scaled to the range (0, 1) across time points/conditions. Finally, metabolic activity heatmaps were generated using the R package ‘pheatmap’ (https://cran.r-project.org/web/packages/pheatmap/index.html).

### Expression cluster analysis

Expression clustering of time course data was performed using the ‘soft clustering’ method of the Bioconductor/R package ‘Mfuzz’ (https://bioconductor.org/). Soft clustering is an unsupervised approach that allows a single gene to potentially be a member of multiple clusters. In initial explorations, the number of clusters generated by Mfuzz was varied to find the number of clusters that was large enough to result in at least 1 cluster with few members and with at least 2 clusters showing visually similar profiles. These criteria ensure that each expression cluster is tightly defined with a highly correlated set of genes while keeping the number of clusters low. Forty-nine clusters met these requirements across all datasets.

Mfuzz performs clustering on mean-centered and scaled expression profiles (Z-scores). Thus, genes are grouped by their relative change in expression, rather than absolute level of expression. In a post-processing step, we confirmed that genes of interest identified through Mfuzz clustering (e.g. CD200R1) had significant absolute expression levels and fold changes.

### Logic simulation

Logic simulation was initially explored using the well-established GINsim (http://ginsim.org/) and booleannet (https://github.com/ialbert/booleannet) simulators. However, logic simulators are designed to explore network steady states, and certain characteristics of the trajectories among these states. As previously noted, the acute and chronic CD8^+^ T cell responses of interest to us comprise specific sequences of *transitory* states with defined transition trajectories. To enable more flexible exploration of such state changes in diverse alternate models, we implemented our network models as a series of logic statements that are executed in a specific order derived from published data and reports.

The R code corresponding to the simulation results shown in Supplementary Figs. 17-20 is given below. This model is one of many alternate models that were explored, and is presented only as an illustrative example. Here, ‘S’ (representing the network state) is a vector of gene activity levels. The symbols ‘&’, ‘|’ and ‘!’ represent logical AND, OR and NOT operators respectively. ‘prevS1’ and ‘prevS2’ are the states of the network at 1 and 2 steps earlier. Genes whose activity depends on earlier network states undergo delayed state changes. The R language symbols “<-“, “->”, and “=” indicate value assignments, and “==” tests equivalence.

~~~
allStates = S; TIME = 0
while(!identical(prevS2, S)) {
 

 TIME <- TIME + 1
 if (TIME == 2) prevS1 = S else if (TIME > 2) {
 prevS2 <- prevS1
 prevS1 <- S
 }

 AKA
 
 S[‘TCR’] <- (S[‘TCR’] & !S[‘PD1’])
 S[‘IL2’] = S[‘IL21’] = S[‘IL12’] <- S[‘TCR’]
 
 S[‘FOXO1’] <- (!S[‘IL2’] & !S[‘AKTAKT’]) & (prevS2[‘TCR’] | prevS2[‘STAT3’])
 S[‘AKT’] <- (S[‘TCR’] & !S[‘PD1’])
 
 S[‘NR4A1’] <- S[‘NFATC1’] | S[‘PD1’]
 S[‘PD1’] <- (((prevS2[‘AP1’] & prevS1[‘AP1’] & S[‘AP1’]) | [‘FOXO1’]) | S[‘NFATC1’])
 
 S[‘AP1.DNA’] <- (prevS1[‘AP1’] | prev S2[‘AP1’] | S[‘AP1’]) & !S[‘BCL6’]
 S[‘BLIMP1’] <- (S[‘BATF.IRF4’] | S[‘IL2’] | S[‘AP1.DNA’])
 
 S[‘IRF4’] <- (S[‘NFkB’] & !S[‘NR4A1’])
 S[‘BCL6’] <- ((S[‘FOXO1’] | S[‘BATF’])) & ! S[‘IRF4’]
 S[‘BCL6’] -> S[‘TCF1’]
 
 S[‘BATF.IRF4’] <- (S[‘BATF’] & S[‘TCR’])
 S[‘BATF’] <- (S[‘IL12’] | S[‘IL21’])
 
 S[‘IL2’] <- (S[‘IL2’] & !S[‘FOXO1’])
 
 S[‘NFATC1’] <- (prevS2[‘NFATC1.med’] | prevS1[‘NFATC1’])
 S[‘NFATC1.med’] <- prevS2[‘NFATC1.lo’]
 S[‘NFATC1.lo’] <- prevS2[‘NFATC2’]
 
 S[‘IFNg’] <- (S[‘NFATC2’] & S[‘AP1’] & !S[‘TCF1’])
 
 S[‘TCR’] -> S[c(‘NFkB’, ‘NFATC2’, ‘AP1’)]
 S[‘AP1’] <- S[‘TCR’] | S[‘AP1’]
 S[‘AKT’] -> S[‘mTOR’] -> S[‘glycolysis’]
 S[‘STAT3’] <- prevS1[‘IL21’]

 allStates <- rbind(allStates, S)

}
~~~

To assess the robustness of the logic models, we assessed how often a model (set of logic functions) passed through the same set of transitory states (and in the same order) when node update assignments were randomized. As an example, the above model passed through the same set of transitory states, in the same order as the above order of statements, in 28 randomized update runs out of 10,000, suggesting the model’s state trajectory is highly dependent on the specified update sequence and time delays.

To visualize the simulation results, the gene states at the end of each update cycle of the simulation were mapped to a network diagram generated using the online tool ‘PathwayMapper’ (http://pathwaymapper.org/).

### Network motif analysis

Combined results from 4 motif detection tools were manually curated to extract network motifs from the network:

i. mfinder (http://www.weizmann.ac.il/mcb/UriAlon/download/network-motif-software)
ii. The Cytoscape ‘Motif-Discovery’ app (http://apps.cytoscape.org/apps/motifdiscovery)
iii. FANMOD (http://theinf1.informatik.uni-jena.de/motifs/)
iv. MAVisto (http://mavisto.ipk-gatersleben.de/).

The above tools provide comprehensive lists of all motif occurrences, but become increasingly unwieldy for motifs with more than a handful of genes. Feed-forward and feedback loops in our network involving multiple steps (gene-gene interactions/network edges) were added manually to the list of found motifs by one author, and then independently verified by two co-authors.

### Differential equation modeling

The robustness and speed up properties of negative feedback illustrated in Supplementary Fig. 24 are adapted from analyses reported by Rosenfeld et al.^50^, and in Uri Alon’s book^23^.

The ordinary differential equation (ODE) model of the network kinetics (Fig. 6) was simulated using Berkeley Madonna (https://berkeley-madonna.myshopify.com/). The model listing is given below (statements following the “;” symbols are descriptive comments not executed by the simulator). Model parameters were selected arbitrarily for purely illustrative purposes.

~~~
METHOD RK4                ; ODE solver
STARTTIME  = 0
STOPTIME  = 100           ; duration of simulation
DT     = 0.01             ; solver timestep

k      = 100              ; pseudo parameter for rapid TCR kinetics

init TCR  = 0             ; TCR is inactive at start
init PD1  = 0             ; PD-1 is inactive at start
init M   = 1              ; BCL6/TCF-1/pro-MPEC activity is on in resting cells
init Blimp1 = 0           ; BLIMP-1 pro-differentiation/effector activity is off
init C   = 0              ; IL2/12/21/IFNg cytokine signaling is off at start

Ag = SQUAREPULSE(0, 75)   ; Antigen activity duration (set to 25 or 75 ‘hours’)

TCR = k * Ag / (1 + PD1^2) - k * TCR   ; TCR complex activity
PD1 = 0.05 * TCR - 0.02 * PD1          ; Inhibitory receptor activation
M = TCR / (1 + Blimp1^2 + C^3) - M     ; pro-memory drivers BCL6 & TCF-1
C = 0.2*Ag^2 / (1 + Ag^2) - 0.1*C      ; IL2/12/21/IFNg signaling & PRC2
Blimp1= C / (1 + M^2) - Blimp1         ; Blimp-1 (effector state marker)
~~~

### Model-based prediction of perturbation effects

To quantify the effects of knocking down the activity of individual genes in our network (e.g. via targeted drugs), we used the R package iGraph (http://igraph.org/r/) to calculate all paths through our network model. Feedback loops that impact TCR-signaling activity (e.g. inhibitory immune receptors) were excluded from this analysis because they affect all downstream genes.

Prediction of impacts using only network topology can be misleading for genes that perform distinct functions at different times (see Fig. 8 and related text). In particular, a target gene can appear to be both activated and repressed by an upstream regulator. For example, in naïve CD8^+^ T cells, ID3 represses *CXCR5* activation by E2A. But following stimulation, ID3 activates *CXCR5* expression. Regulatory interactions that could not be time-resolved due to lack of time course data were excluded from our analyses.

### In vitro T cell cultures

Total T cells were isolated from peripheral blood mononuclear cells from 3 human donors via negative selection (Stem Cell Technologie cat. no. 19051) and plated in anti-CD3 (BD Pharmingen clone UCHT1, cat. no. 555329) pre-coated (10 μg/ml in PBS overnight at 4°C) 96- well round bottom plates at 100,000 cells per well in RPMI1640, 10% FBS, 0.1 mM NEAA, 1 mM Na pyruvate, 5 ng/ml anti-CD28 (eBiosciences clone CD28.2, cat no. 16-0289-85), alone or with 5 μM (reported), 0.5 μM (not shown) EZH2 inhibitor, CPI-169 (APExBio B4678) or DMSO. The same volume of EZH2 inhibitor and DMSO was added to each well to control for any DMSO effect. T cells were cultured at 37°C, 5% CO^2^. T cells were harvested on days 1, 3, 4, 8, washed in PBS and re-suspended in 350 μl RLT (Qiagen cat. no. 74136) and stored at −80°C for future RNA isolation (Qiagen cat. no. 74136). Total T cells were also sampled and processed as other cultures on the day of isolation (day 0) and after 24 hours of incubation with 10 ng/ml human interleukin-7 (RnD Systems, cat. no. 207-IL-025) as controls for gene expression analysis.

### RNA isolation and cDNA reverse transcription

RNA was isolated from all samples using qiagen kit (Qiagen cat. no. 74136) and quantitated on a Nanodrop 2000 spectrophotometer (ThermoScientific). 1.5 μg RNA was reverse transcribed into cDNA as per protocol (Applied Biosystems cat. no. 4368814).

### Quantitative real-time PCR

qRT-PCR was performed using TaqMan Fast Advanced Master Mix (Applied Biosystems cat. no. 4444557) in a ViiA7 system (Applied Biosystems) using Applied Biosystems primers. Gene expression was quantified as per Livak & Schmittgen^51^ normalized to GUSB. All measurements were performed in triplicate.

### Data availability

All data analyzed in this manuscript have been previously published and are available publicly, as described in the Methods.

## Acknowledgements

The authors received editorial support, provided by Excerpta Medica, supported by Celgene Corporation. The authors are fully responsible for all content and editorial decisions.

## Author contributions

Except for bifurcation/sensitivity analyses, which were performed by L.H. and C.C.S., all data processing and analyses were performed by H.B. in consultation with all other authors. The initial version of the manuscript was drafted by H.B. and then edited by all co-authors.

