## Supplemental Tables for "Integrative network modeling reveals mechanisms underlying T cell exhaustion"

### **Supplementary Table 1. Large-scale TCE network specifications**

| **Source** | **Target** | **Reference** | **EdgeType** |
| --- | --- | --- | --- |
| AKT1 | BACH2 | ^1^ | Inhibits |
| AKT1 | FOXO1 | ^2,3^ | Inhibits |
| AKT1 | MTOR | ^4^ | Promotes |
| AKT1 | PPARGC1A | ^5^ | Inhibits |
| AKT1 | SLC2A1 | ^6^ | Promotes |
| AKT1 | SLC2A2 | ^6^ | Promotes |
| AKT1 | SLC2A3 | ^6^ | Promotes |
| AKT1 | SLC2A4 | ^6^ | Promotes |
| AKT1 | SLC2A5 | ^6^ | Promotes |
| AKT2 | BACH2 | ^1^ | Inhibits |
| AKT2 | FOXO1 | ^2,3^ | Inhibits |
| AKT2 | MTOR | ^4^ | Promotes |
| AKT2 | PPARGC1A | ^5^ | Inhibits |
| AKT2 | SLC2A1 | ^6^ | Promotes |
| AKT2 | SLC2A2 | ^6^ | Promotes |
| AKT2 | SLC2A3 | ^6^ | Promotes |
| AKT2 | SLC2A4 | ^6^ | Promotes |
| AKT2 | SLC2A5 | ^6^ | Promotes |
| AKT3 | BACH2 | ^1^ | Inhibits |
| AKT3 | FOXO1 | ^2,3^ | Inhibits |
| AKT3 | MTOR | ^4^ | Promotes |
| AKT3 | PPARGC1A | ^5^ | Inhibits |
| AKT3 | SLC2A1 | ^6^ | Promotes |
| AKT3 | SLC2A2 | ^6^ | Promotes |
| AKT3 | SLC2A3 | ^6^ | Promotes |
| AKT3 | SLC2A4 | ^6^ | Promotes |
| AKT3 | SLC2A5 | ^6^ | Promotes |
| BACH2 | GATA3 | ^7^ | Inhibits |
| BACH2 | ID3 | ^7^ | Inhibits |
| BACH2 | JUN | ^7^ | Inhibits |
| BACH2 | JUNB | ^1,7^ | Inhibits |
| BACH2 | JUND | ^1,7^ | Inhibits |
| BACH2 | PRDM1 | ^7,8^ | Inhibits |
| BATF | BCL6 | ^9^ | Promotes |
| BATF | PRDM1 | ^10^ | Promotes |
| BATF | EOMES | ^11^ | Promotes |
| BATF | IRF4 | ^11^ | Inhibits |
| BATF | NFATC2 | ^11^ | Promotes |
| BATF | NF-κB1 | ^11^ | Promotes |
| BATF | RUNX3 | ^11^ | Promotes |
| BATF | TBX21 | ^11^ | Promotes |
| BCL6 | GZMB | ^12^ | Inhibits |
| BCL6 | ID2 | ^13^ | Inhibits |
| BCL6 | LEF1 | ^13^ | Promotes |
| BCL6 | PRDM1 | ^13,14^ | Inhibits |
| BCL6 | TCF7 | ^13^ | Promotes |
| BTLA | AKT1 | ^15^ | Inhibits |
| BTLA | AKT2 | ^15^ | Inhibits |
| BTLA | AKT3 | ^15^ | Inhibits |
| BTLA | NF-κB1 | ^15^ | Inhibits |
| BTLA | NF-κB2 | ^15^ | Inhibits |
| BTLA | PIK3CA | ^15^ | Inhibits |
| BTLA | PIK3CB | ^15^ | Inhibits |
| BTLA | PIK3CG | ^15^ | Inhibits |
| BTLA | PRKCQ | ^15^ | Inhibits |
| BTLA | REL | ^15^ | Promotes |
| BTLA | RELA | ^15^ | Inhibits |
| BTLA | RELB | ^15^ | Inhibits |
| CA | NFATC1 | ^16^ | Promotes |
| CA | NFATC2 | ^16^ | Promotes |
| CD160 | IFN-γ | ^17^ | Inhibits |
| CD160 | IL2 | ^17^ | Inhibits |
| CD244 | CA | ^18^ | Promotes |
| CD244 | MAPK1 | ^18^ | Promotes |
| CD244 | MAPK3 | ^18^ | Promotes |
| CD244 | NFATC1 | ^18^ | Promotes |
| CD244 | NFATC2 | ^18^ | Promotes |
| CD244 | NF-κB1 | ^19^ | Promotes |
| CD244 | NF-κB2 | ^19^ | Promotes |
| CD244 | REL | ^19^ | Promotes |
| CD244 | RELA | ^19^ | Promotes |
| CD244 | RELB | ^19^ | Promotes |
| CD247 | AKT1 | ^20^ | Promotes |
| CD247 | AKT2 | ^20^ | Promotes |
| CD247 | AKT3 | ^20^ | Promotes |
| CD247 | DNMT3A | ^21^ | Promotes |
| CD247 | FOS | ^22^ | Promotes |
| CD247 | JUN | ^22^ | Promotes |
| CD247 | JUNB | ^22^ | Promotes |
| CD247 | JUND | ^22^ | Promotes |
| CD247 | NFATC1 | ^16^ | Promotes |
| CD247 | NFATC2 | ^16^ | Promotes |
| CD247 | NF-κB1 | ^23^ | Promotes |
| CD247 | NF-κB2 | ^23^ | Promotes |
| CD247 | REL | ^23^ | Promotes |
| CD247 | RELA | ^23^ | Promotes |
| CD247 | RELB | ^23^ | Promotes |
| CD247 | ZAP70 | ^24^ | Promotes |
| CD274 | ZAP70 | ^25^ | Inhibits |
| CD28 | DNMT3A | ^21^ | Promotes |
| CD28 | HRAS | ^15^ | Promotes |
| CD28 | KRAS | ^15^ | Promotes |
| CD28 | NF-κB1 | ^26^ | Promotes |
| CD28 | NRAS | ^15^ | Promotes |
| CD28 | PIK3CA | ^26^ | Promotes |
| CD28 | PIK3CB | ^26^ | Promotes |
| CD28 | PIK3CG | ^26^ | Promotes |
| CD28 | PRKCQ | ^26,27^ | Promotes |
| CD28 | RELA | ^26^ | Promotes |
| CD3D | AKT1 | ^20^ | Promotes |
| CD3D | AKT2 | ^20^ | Promotes |
| CD3D | AKT3 | ^20^ | Promotes |
| CD3D | DNMT3A | ^21^ | Promotes |
| CD3D | FOS | ^22^ | Promotes |
| CD3D | JUN | ^22^ | Promotes |
| CD3D | JUNB | ^22^ | Promotes |
| CD3D | JUND | ^22^ | Promotes |
| CD3D | NFATC1 | ^16^ | Promotes |
| CD3D | NFATC2 | ^16^ | Promotes |
| CD3D | NF-κB1 | ^23^ | Promotes |
| CD3D | NF-κB2 | ^23^ | Promotes |
| CD3D | REL | ^23^ | Promotes |
| CD3D | RELA | ^23^ | Promotes |
| CD3D | RELB | ^23^ | Promotes |
| CD3D | ZAP70 | ^24^ | Promotes |
| CD3E | AKT1 | ^20^ | Promotes |
| CD3E | AKT2 | ^20^ | Promotes |
| CD3E | AKT3 | ^20^ | Promotes |
| CD3E | DNMT3A | ^21^ | Promotes |
| CD3E | FOS | ^22^ | Promotes |
| CD3E | JUN | ^22^ | Promotes |
| CD3E | JUNB | ^22^ | Promotes |
| CD3E | JUND | ^22^ | Promotes |
| CD3E | NFATC1 | ^16^ | Promotes |
| CD3E | NFATC2 | ^16^ | Promotes |
| CD3E | NF-κB1 | ^23^ | Promotes |
| CD3E | NF-κB2 | ^23^ | Promotes |
| CD3E | REL | ^23^ | Promotes |
| CD3E | RELA | ^23^ | Promotes |
| CD3E | RELB | ^23^ | Promotes |
| CD3E | ZAP70 | ^24^ | Promotes |
| CD3G | AKT1 | ^20^ | Promotes |
| CD3G | AKT2 | ^20^ | Promotes |
| CD3G | AKT3 | ^20^ | Promotes |
| CD3G | DNMT3A | ^21^ | Promotes |
| CD3G | FOS | ^22^ | Promotes |
| CD3G | JUN | ^22^ | Promotes |
| CD3G | JUNB | ^22^ | Promotes |
| CD3G | JUND | ^22^ | Promotes |
| CD3G | NFATC1 | ^16^ | Promotes |
| CD3G | NFATC2 | ^16^ | Promotes |
| CD3G | NF-κB1 | ^23^ | Promotes |
| CD3G | NF-κB2 | ^23^ | Promotes |
| CD3G | REL | ^23^ | Promotes |
| CD3G | RELA | ^23^ | Promotes |
| CD3G | RELB | ^23^ | Promotes |
| CD3G | ZAP70 | ^24^ | Promotes |
| CD8A | CD247 | ^28^ | Promotes |
| CD8A | CD3D | ^28^ | Promotes |
| CD8A | CD3E | ^28^ | Promotes |
| CD8A | CD3G | ^28^ | Promotes |
| CD8A | DNMT3A | ^21^ | Promotes |
| CD8A | LCK | ^26^ | Promotes |
| CTLA4 | AKT1 | ^15^ | Inhibits |
| CTLA4 | AKT2 | ^15^ | Inhibits |
| CTLA4 | AKT3 | ^15^ | Inhibits |
| CTLA4 | NF-κB1 | ^15^ | Inhibits |
| CTLA4 | NF-κB2 | ^15^ | Inhibits |
| CTLA4 | REL | ^15^ | Inhibits |
| CTLA4 | RELA | ^15^ | Inhibits |
| CTLA4 | RELB | ^15^ | Inhibits |
| DNMT3A | DNMT3A | ^29^ | Inhibits |
| DNMT3A | TCF7 | ^30,31^ | Inhibits |
| DNMT3B | DNMT3A | ^29^ | Inhibits |
| DNMT3L | DNMT3A | ^29^ | Inhibits |
| EGR1 | ID3 | ^32^ | Promotes |
| EGR1 | IL2 | ^33^ | Promotes |
| EGR1 | IL2RB | ^34^ | Promotes |
| EGR2 | GZMB | ^35^ | Inhibits |
| EGR2 | ID3 | ^35^ | Promotes |
| EGR2 | IFN-γ | ^35^ | Inhibits |
| EGR2 | LAG3 | ^36^ | Promotes |
| EGR2 | MYC | ^35^ | Promotes |
| EGR2 | PRDM1 | ^35^ | Inhibits |
| EGR2 | TBX21 | ^35^ | Inhibits |
| EGR2 | TCF7 | ^35^ | Promotes |
| EGR2 | ZEB2 | ^35^ | Promotes |
| EGR3 | GZMB | ^35^ | Inhibits |
| EGR3 | ID3 | ^35^ | Promotes |
| EGR3 | IFN-γ | ^35^ | Inhibits |
| EGR3 | MYC | ^35^ | Promotes |
| EGR3 | PRDM1 | ^35^ | Inhibits |
| EGR3 | TBX21 | ^35^ | Inhibits |
| EGR3 | TCF7 | ^35^ | Promotes |
| EGR3 | ZEB2 | ^35^ | Promotes |
| Exhaustion | ZN | ^37^ | Promotes |
| EZH2 | DNMT3A | ^38^ | Promotes |
| EZH2 | DNMT3B | ^38^ | Promotes |
| EZH2 | DNMT3L | ^38^ | Promotes |
| FOS | IFN-γ | ^39^ | Promotes |
| FOS | IL2 | ^40^ | Promotes |
| FOS | IL2RB | ^41^ | Promotes |
| FOS | PDCD1 | ^42^ | Promotes |
| FOS | PRDM1 | ^14,40^ | Promotes |
| FOXO1 | BCL6 | ^43^ | Promotes |
| FOXO1 | EZH2 | ^44^ | Inhibits |
| FOXO1 | PDCD1 | ^45,46^ | Promotes |
| GATA3 | BATF | ^11^ | Promotes |
| GATA3 | MYC | ^47^ | Promotes |
| HIF1A | BATF | ^11^ | Promotes |
| HIF1A | MYC | ^48^ | Promotes |
| HRAS | MAPK1 | ^49^ | Promotes |
| HRAS | MAPK3 | ^49^ | Promotes |
| HRAS | MAPK8 | ^49^ | Promotes |
| ICOS | MAF | ^15^ | Promotes |
| ID2 | TCF3 | ^13^ | Inhibits |
| ID3 | TCF3 | ^13^ | Inhibits |
| IFNAR1 | BCL6 | ^50,51^ | Promotes |
| IFNAR1 | MT1 | ^52^ | Promotes |
| IFNAR1 | MT2 | ^52^ | Promotes |
| IFNAR1 | PDCD1 | ^42^ | Promotes |
| IFNAR1 | PIK3CA | ^53^ | Promotes |
| IFNAR1 | PIK3CB | ^53^ | Promotes |
| IFNAR1 | PIK3CG | ^53^ | Promotes |
| IFNAR1 | PRKCQ | ^53^ | Promotes |
| IFNAR1 | STAT1 | ^54^ | Promotes |
| IFNAR1 | STAT4 | ^55^ | Promotes |
| IFNAR2 | BCL6 | ^50,51^ | Promotes |
| IFNAR2 | MT1 | ^52^ | Promotes |
| IFNAR2 | MT2 | ^52^ | Promotes |
| IFNAR2 | PDCD1 | ^42^ | Promotes |
| IFNAR2 | PIK3CA | ^53^ | Promotes |
| IFNAR2 | PIK3CB | ^53^ | Promotes |
| IFNAR2 | PIK3CG | ^53^ | Promotes |
| IFNAR2 | PRKCQ | ^53^ | Promotes |
| IFNAR2 | STAT1 | ^54^ | Promotes |
| IFNAR2 | STAT4 | ^55^ | Promotes |
| IFN-γ | EGR2 | ^35^ | Inhibits |
| IFN-γ | EGR3 | ^35^ | Inhibits |
| IFN-γ | MT1 | ^52^ | Promotes |
| IFN-γ | MT2 | ^52^ | Promotes |
| IFN-γ | PIK3CA | ^53^ | Promotes |
| IFN-γ | PIK3CB | ^53^ | Promotes |
| IFN-γ | PIK3CG | ^53^ | Promotes |
| IFN-γ | PRKCQ | ^53^ | Promotes |
| IFN-γ | SLC2A1 | ^56^ | Inhibits |
| IFN-γ | SLC2A2 | ^56^ | Inhibits |
| IFN-γ | SLC2A3 | ^56^ | Inhibits |
| IFN-γ | SLC2A4 | ^56^ | Inhibits |
| IFN-γ | SLC2A5 | ^56^ | Inhibits |
| IFN-γ | STAT1 | ^57^ | Inhibits |
| IFN-γ | STAT2 | ^57^ | Inhibits |
| IFNGR1 | EGR2 | ^35^ | Inhibits |
| IFNGR1 | EGR3 | ^35^ | Inhibits |
| IFNGR1 | MT2 | ^52^ | Promotes |
| IFNGR1 | PIK3CA | ^53^ | Promotes |
| IFNGR1 | PIK3CB | ^53^ | Promotes |
| IFNGR1 | PIK3CG | ^53^ | Promotes |
| IFNGR1 | PRKCQ | ^53^ | Promotes |
| IFNGR2 | EGR2 | ^35^ | Inhibits |
| IFNGR2 | EGR3 | ^35^ | Inhibits |
| IFNGR2 | MT2 | ^52^ | Promotes |
| IFNGR2 | PIK3CA | ^53^ | Promotes |
| IFNGR2 | PIK3CB | ^53^ | Promotes |
| IFNGR2 | PIK3CG | ^53^ | Promotes |
| IFNGR2 | PRKCQ | ^53^ | Promotes |
| IL12RB1 | BATF | ^58^ | Promotes |
| IL12RB1 | ID2 | ^59^ | Promotes |
| IL12RB1 | PRDM1 | ^59^ | Promotes |
| IL12RB1 | STAT4 | ^60,61^ | Promotes |
| IL12RB1 | TBX21 | ^62^ | Promotes |
| IL12RB2 | BATF | ^58^ | Promotes |
| IL12RB2 | ID2 | ^59^ | Promotes |
| IL12RB2 | PRDM1 | ^59^ | Promotes |
| IL12RB2 | STAT4 | ^60,61^ | Promotes |
| IL12RB2 | TBX21 | ^62^ | Promotes |
| IL2 | BCL6 | ^41^ | Inhibits |
| IL2 | IL2RB | ^41^ | Promotes |
| IL21R | BATF | ^10^ | Promotes |
| IL21R | STAT3 | ^63,64^ | Promotes |
| IL2RA | BCL6 | ^43,65^ | Inhibits |
| IL2RA | FOXO1 | ^43,66^ | Inhibits |
| IL2RA | GATA3 | ^67^ | Promotes |
| IL2RA | HIF1A | ^65^ | Promotes |
| IL2RA | ID2 | ^59^ | Promotes |
| IL2RA | PRDM1 | ^59^ | Promotes |
| IL2RB | BCL6 | ^43,65^ | Inhibits |
| IL2RB | CD160 | ^68^ | Promotes |
| IL2RB | CD244 | ^68^ | Promotes |
| IL2RB | FOXO1 | ^43,66^ | Inhibits |
| IL2RB | GATA3 | ^67^ | Promotes |
| IL2RB | HAVCR2 | ^68^ | Promotes |
| IL2RB | HIF1A | ^65^ | Promotes |
| IL2RB | ID2 | ^59^ | Promotes |
| IL2RB | LAG3 | ^68^ | Promotes |
| IL2RB | MYC | ^68^ | Promotes |
| IL2RB | PDCD1 | ^68^ | Promotes |
| IL2RB | PRDM1 | ^59^ | Promotes |
| IRF4 | BCL6 | ^10,69^ | Inhibits |
| IRF4 | PRDM1 | ^10,69^ | Promotes |
| JUN | IFN-γ | ^39^ | Promotes |
| JUN | IFN-γ R1 | ^39^ | Promotes |
| JUN | IFN-γ R2 | ^39^ | Promotes |
| JUN | IL2 | ^40^ | Promotes |
| JUN | IL2RB | ^41^ | Promotes |
| JUN | PDCD1 | ^42^ | Promotes |
| JUN | PRDM1 | ^14,40^ | Promotes |
| JUNB | IFN-γ | ^39^ | Promotes |
| JUNB | IL2 | ^40^ | Promotes |
| JUNB | IL2RB | ^41^ | promotes |
| JUNB | PDCD1 | ^42^ | Promotes |
| JUNB | PRDM1 | ^40^ | Promotes |
| JUND | IFN-γ | ^39^ | Promotes |
| JUND | IL2 | ^40^ | Promotes |
| JUND | IL2RB | ^41^ | Promotes |
| JUND | PDCD1 | ^42^ | Promotes |
| JUND | PRDM1 | ^40^ | Promotes |
| KDM6B | EZH2 | ^70^ | Inhibits |
| KRAS | MAPK1 | ^49^ | Promotes |
| KRAS | MAPK3 | ^49^ | Promotes |
| KRAS | MAPK8 | ^49^ | Promotes |
| LCK | ZAP70 | ^26^ | Promotes |
| LEF1 | BCL6 | ^71^ | Promotes |
| LEF1 | CXCR5 | ^72^ | Promotes |
| LEF1 | EOMES | ^73^ | Promotes |
| LEF1 | GZMA | ^73^ | Inhibits |
| LEF1 | GZMB | ^73^ | Inhibits |
| LEF1 | IFN-γ | ^74^ | Inhibits |
| LEF1 | KLRG1 | ^73^ | Inhibits |
| LEF1 | LEF1 | ^13^ | Promotes |
| LEF1 | MYC | ^73^ | Promotes |
| LEF1 | PRDM1 | ^13^ | Promotes |
| LEF1 | PRF1 | ^75^ | Inhibits |
| MAPK1 | EGR1 | ^76^ | Promotes |
| MAPK1 | FOS | ^26^ | Promotes |
| MAPK1 | JUN | ^26^ | Promotes |
| MAPK1 | JUNB | ^26^ | Promotes |
| MAPK1 | JUND | ^26^ | Promotes |
| MAPK3 | EGR1 | ^76^ | Promotes |
| MAPK3 | FOS | ^26^ | Promotes |
| MAPK3 | JUN | ^26^ | Promotes |
| MAPK3 | JUNB | ^26^ | Promotes |
| MAPK3 | JUND | ^26^ | Promotes |
| MAPK8 | FOS | ^26^ | Promotes |
| MAPK8 | JUN | ^26^ | Promotes |
| MAPK8 | JUNB | ^26^ | Promotes |
| MAPK8 | JUND | ^26^ | Promotes |
| MT1 | MAPK1 | ^77^ | Promotes |
| MT1 | MAPK3 | ^77^ | Promotes |
| MT1 | ZN | ^37^ | Promotes |
| MT2 | ZN | ^37^ | Promotes |
| MTOR | Glycolysis | ^4^ | Promotes |
| MTOR | RICTOR | ^78^ | Promotes |
| MTOR | RPTOR | ^78^ | Promotes |
| MYC | Proliferation | ^48^ | Promotes |
| NFATC1 | IFN-γ | ^39,79^ | Promotes |
| NFATC1 | IFNGR1 | ^39^ | Promotes |
| NFATC1 | IFNGR2 | ^39^ | Promotes |
| NFATC1 | IL2 | ^40^ | Promotes |
| NFATC1 | IL2RB | ^41^ | Promotes |
| NFATC1 | NFATC1 | ^80^ | Promotes |
| NFATC1 | NR4A1 | ^81^ | Promotes |
| NFATC1 | PDCD1 | ^42^ | Promotes |
| NFATC1 | PRDM1 | ^40^ | Promotes |
| NFATC2 | EGR2 | ^36^ | Promotes |
| NFATC2 | EGR3 | ^35^ | Promotes |
| NFATC2 | IFN-γ | ^39,79^ | Promotes |
| NFATC2 | IL2 | ^40^ | Promotes |
| NFATC2 | IL2RB | ^41^ | Promotes |
| NFATC2 | NFATC1 | ^82^ | Promotes |
| NFATC2 | PDCD1 | ^42^ | Promotes |
| NFATC2 | PRDM1 | ^40^ | Promotes |
| NF-κB1 | IL2 | ^40^ | Promotes |
| NF-κB1 | IL2RB | ^41^ | Promotes |
| NF-κB1 | IRF4 | ^83^ | Promotes |
| NF-κB1 | PDCD1 | ^42^ | Promotes |
| NF-κB1 | PRDM1 | ^40^ | Promotes |
| NF-κB2 | IRF4 | ^83^ | Promotes |
| NR4A1 | Glycolysis | ^84^ | Promotes |
| NR4A1 | IRF4 | ^85^ | Inhibits |
| NR4A2 | Glycolysis | ^86^ | Promotes |
| NR4A3 | Glycolysis | ^87^ | Promotes |
| NRAS | MAPK1 | ^49^ | Promotes |
| NRAS | MAPK3 | ^49^ | Promotes |
| NRAS | MAPK8 | ^49^ | Promotes |
| PDCD1 | AKT1 | ^88^ | Promotes |
| PDCD1 | AKT2 | ^88^ | Promotes |
| PDCD1 | AKT3 | ^88^ | Promotes |
| PDCD1 | CD247 | ^89^ | Inhibits |
| PDCD1 | CD28 | ^89^ | Inhibits |
| PDCD1 | CD3D | ^89^ | Inhibits |
| PDCD1 | CD3E | ^89^ | Inhibits |
| PDCD1 | CD3G | ^89^ | Inhibits |
| PDCD1 | CD8A | ^89^ | Inhibits |
| PDCD1 | NR4A1 | ^86^ | Inhibits |
| PDCD1 | BATF | ^90^ | Promotes |
| PDCD1 | HRAS | ^91^ | Inhibits |
| PDCD1 | KRAS | ^91^ | Inhibits |
| PDCD1 | NRAS | ^91^ | Inhibits |
| PDCD1 | PIK3CA | ^91^ | Inhibits |
| PDCD1 | PIK3CB | ^91^ | Inhibits |
| PDCD1 | PIK3CG | ^91^ | Inhibits |
| PDCD1 | PPARGC1A | ^88^ | Inhibits |
| PDCD1 | PRKCQ | ^15^ | Inhibits |
| PDCD1 | SLC2A1 | ^56^ | Promotes |
| PDCD1 | SLC2A2 | ^56^ | Promotes |
| PDCD1 | SLC2A3 | ^56^ | Promotes |
| PDCD1 | SLC2A4 | ^56^ | Promotes |
| PDCD1 | SLC2A5 | ^56^ | Promotes |
| PDCD1 | ZAP70 | ^25^ | Inhibits |
| PPARGC1A | Proliferation | ^5^ | Promotes |
| PRDM1 | BCL6 | ^13,92,93^ | Inhibits |
| PRDM1 | CXCR5 | ^13^ | Inhibits |
| PRDM1 | ID3 | ^94^ | Inhibits |
| PRDM1 | IL2 | ^94^ | Inhibits |
| PRDM1 | MYC | ^95^ | Inhibits |
| PRDM1 | NFATC1 | ^96^ | Inhibits |
| PRDM1 | PDCD1 | ^96^ | Inhibits |
| PRDM1 | TCF7 | ^13^ | Inhibits |
| PRKCQ | NF-κB1 | ^97^ | Promotes |
| PRKCQ | RELA | ^97^ | Promotes |
| REL | IRF4 | ^83^ | Promotes |
| RELA | IL2 | ^40^ | Promotes |
| RELA | IL2RB | ^41^ | Promotes |
| RELA | IRF4 | ^83^ | Promotes |
| RELA | PDCD1 | ^42^ | Promotes |
| RELA | PRDM1 | ^40^ | Promotes |
| RELB | IRF4 | ^83^ | Promotes |
| RUNX3 | BATF | ^11^ | Promotes |
| RUNX3 | IFN-γ | ^98^ | Promotes |
| SLC39A14 | ZN | ^99^ | Promotes |
| SLC39A8 | ZN | ^99^ | Promotes |
| STAT1 | BCL6 | ^50,51^ | Promotes |
| STAT1 | CXCR5 | ^50,51^ | Inhibits |
| STAT1 | PDCD1 | ^50,51^ | Promotes |
| STAT2 | BCL6 | ^50,51^ | Promotes |
| STAT3 | BATF | ^11^ | Promotes |
| STAT3 | FOXO1 | ^100^ | Promotes |
| STAT4 | IFN-γ | ^55^ | Promotes |
| STAT4 | IFNGR1 | ^55^ | Promotes |
| STAT4 | IFNGR2 | ^55^ | Promotes |
| STAT4 | STAT1 | ^101^ | Inhibits |
| STAT4 | TBX21 | ^55^ | Promotes |
| TBX21 | BCL6 | ^102^ | Inhibits |
| TBX21 | BTLA | ^103^ | Inhibits |
| TBX21 | CD160 | ^103^ | Inhibits |
| TBX21 | HAVCR2 | ^104^ | Promotes |
| TBX21 | IFN-γ | ^98^ | Promotes |
| TBX21 | IL2RB | ^105^ | Promotes |
| TBX21 | LAG3 | ^103^ | Inhibits |
| TBX21 | PDCD1 | ^103^ | Inhibits |
| TBX21 | RUNX3 | ^98^ | Promotes |
| TBX21 | ZEB2 | ^106^ | Promotes |
| TCF3 | CXCR5 | ^13^ | Promotes |
| TCF3 | TCF7 | ^13^ | Promotes |
| TCF7 | BCL6 | ^71^ | Promotes |
| TCF7 | CXCR5 | ^72^ | Promotes |
| TCF7 | EOMES | ^73^ | Promotes |
| TCF7 | GATA3 | ^74^ | Promotes |
| TCF7 | GZMA | ^73^ | Inhibits |
| TCF7 | GZMB | ^73^ | Inhibits |
| TCF7 | IFN-γ | ^74^ | Inhibits |
| TCF7 | IFNGR1 | ^74^ | Inhibits |
| TCF7 | IFNGR2 | ^74^ | Inhibits |
| TCF7 | KLRG1 | ^73^ | Inhibits |
| TCF7 | MYC | ^73^ | Promotes |
| TCF7 | PRDM1 | ^13^ | Inhibits |
| TCF7 | PRF1 | ^75^ | Inhibits |
| TET1 | DNMT3A | ^107,108^ | Inhibits |
| TET1 | DNMT3B | ^107,108^ | Inhibits |
| TET1 | DNMT3L | ^107,108^ | Inhibits |
| TET1 | PDCD1 | ^109^ | Promotes |
| TET2 | DNMT3A | ^107,108^ | Inhibits |
| TET2 | DNMT3B | ^107,108^ | Inhibits |
| TET2 | DNMT3L | ^107,108^ | Inhibits |
| TET2 | PDCD1 | ^109^ | Promotes |
| TET3 | DNMT3A | ^107,108^ | Inhibits |
| TET3 | DNMT3B | ^107,108^ | Inhibits |
| TET3 | DNMT3L | ^107,108^ | Inhibits |
| TET3 | PDCD1 | ^109^ | Promotes |
| TIGIT | MAPK1 | ^110^ | Inhibits |
| TIGIT | MAPK3 | ^110^ | Inhibits |
| TIGIT | NF-κB1 | ^110^ | Inhibits |
| TIGIT | NF-κB2 | ^110^ | Inhibits |
| TIGIT | PIK3CA | ^110^ | Inhibits |
| TIGIT | PIK3CB | ^110^ | Inhibits |
| TIGIT | PIK3CG | ^110^ | Inhibits |
| TIGIT | REL | ^110^ | Inhibits |
| TIGIT | RELA | ^110^ | Inhibits |
| TIGIT | RELB | ^110^ | Inhibits |
| TNF | NF-κB1 | ^111^ | Promotes |
| TNF | NF-κB2 | ^111^ | Promotes |
| TNF | REL | ^111^ | Promotes |
| TNF | RELA | ^111^ | Promotes |
| TNF | RELB | ^111^ | Promotes |
| TNFRSF1A | TNF | ^111^ | Promotes |
| TNFRSF1B | TNF | ^111^ | Promotes |
| TNFRSF9 | CD247 | ^112^ | Promotes |
| TNFRSF9 | CD28 | ^112^ | Promotes |
| TNFRSF9 | CD3D | ^112^ | Promotes |
| TNFRSF9 | CD3E | ^112^ | Promotes |
| TNFRSF9 | CD3G | ^112^ | Promotes |
| TNFRSF9 | CD8A | ^112^ | Promotes |
| TNFRSF9 | FOXO1 | ^113^ | Inhibits |
| ZAP70 | Adhesion | ^24^ | Promotes |
| ZAP70 | CA | ^24^ | Promotes |
| ZAP70 | HRAS | ^24^ | Promotes |
| ZAP70 | KRAS | ^24^ | Promotes |
| ZAP70 | NRAS | ^24^ | Promotes |
| ZAP70 | PRKCQ | ^26,27^ | Promotes |
| ZEB1 | IL2 | ^114^ | Inhibits |
| ZEB2 | IL2 | ^114^ | Inhibits |
| ZEB2 | TBX21 | ^115^ | Promotes |
| ZN | MT1 | ^77^ | Promotes |
| ZN | MT2 | ^77^ | Promotes |

### **Supplementary Table 2. Reduced TCE network specifications**

| **Source** | **Target** | **Interaction** | **Reference** | **Comment** |
| --- | --- | --- | --- | --- |
| 4-1BB | FAS/FASL | Promotes | ^116,117^ | Delayed, peaks at ~3 days |
| 4-1BB | TCF-1 | Promotes | ^113^ |  |
| AKT | BACH2 | Promotes | ^1^ |  |
| AKT | NFATC1,2 | Promotes | ^118-120^ | NFATs are activated by Ca^++^ signaling. AKT stops nuclear export of activated NFATs. |
| AKT | MTOR | Promotes | ^121,122^ |  |
| AKT | NF-κB | Promotes | ^123^ |  |
| AKT | FOXO1 | Inhibits | ^2^ |  |
| AKT | AP1 | Promotes | ^124^ |  |
| AP1 (JUN:FOS) | NR4A1 | Promotes | ^125,126^ | AP1 causes NR4A1 nuclear export, which activates its pro-apoptotic activity in mitochondria. See also ^127,128^ |
| AP1 (JUN:FOS) | NFATC1:AP1:IRF4:BATF | Inhibits | ^129-132^ | BATF & AP1 compete for the same sites^131^ |
| BACH2 | IFN-γ | Inhibits | ^1^ |  |
| BACH2 | BLIMP-1 | Inhibits | ^133^ |  |
| BACH2 | ID3 | Promotes | ^134^ |  |
| BATF | NFATC1:AP1:IRF4:BATF | Promotes | ^10,130,132,135^ | See also ^136^ |
| BCL6 | TCF-1 | Promotes | ^13,71,72^ |  |
| BCL6 | BLIMP-1 | Inhibits | ^137^ |  |
| BLIMP-1 | TIGIT | Promotes | ^138^ |  |
| BLIMP-1 | CD160 | Promotes | ^139,140^ |  |
| BLIMP-1 | CD244/2B4 | Promotes | ^139,140^ |  |
| BLIMP-1 | BCL6 | Inhibits | ^141^ |  |
| BLIMP-1 | ID3 | Inhibits | ^142^ |  |
| BTLA | CD3, TCR, CD8, CD28 | Inhibits | ^143^ |  |
| CD137L | 4-1BB | Promotes | ^144^ |  |
| CD3, TCR, CD8, CD28 | 4-1BB | Promotes | ^145^ |  |
| CD3, TCR, CD8, CD28 | RAS | Promotes | ^15,146^ |  |
| CD3, TCR, CD8, CD28 | EZH2/PRC2 | Promotes | ^147^ | PRC2 has a dual role^148^ |
| CD3, TCR, CD8, CD28 | PI3K | Promotes | ^15^ |  |
| CTLA4 | Tregs | Promotes | ^149^ |  |
| CTLA4 | CD3, TCR, CD8, CD28 | Inhibits | ^149^ |  |
| E2A | CXCR5 | Promotes | ^150^ |  |
| EGR2/3 | TBET:ZEB2 | Inhibits | ^35,151^ |  |
| EGR2/3 | LAG3 | Promotes | ^152^ |  |
| EGR2/3 | TCF-1 | Promotes | ^35^ |  |
| EGR2/3 | ID3 | Promotes | ^35^ |  |
| EGRs | FAS/FASL | Promotes | ^153,154^ |  |
| EOMES | GZMA/B | Promotes | ^155^ |  |
| EOMES | PRF1 | Promotes | ^155^ |  |
| EOMES | IFN-γ | Promotes | ^155^ |  |
| EZH2/PRC2 | EOMES | Inhibits | ^44^ |  |
| EZH2/PRC2 | TCF-1 | Inhibits | ^44^ |  |
| EZH2/PRC2 | BACH2 | Inhibits | ^44^ |  |
| EZH2/PRC2 | FOXO1 | Inhibits | ^44^ |  |
| FAS/FASL | FAS/FASL | Promotes | ^156^ |  |
| FOXO1 | PGC1A | Promotes | ^5,157-160^ |  |
| FOXO1 | TCF-1 | Promotes | ^161^ |  |
| HIF1A | MYC | Promotes | ^48^ |  |
| HIF1A | PPARa | Inhibits | ^162,163^ |  |
| ID3 | FAS/FASL | Inhibits | ^164^ |  |
| ID3 | IL2R | Inhibits | ^165^ |  |
| ID3:Tnaïve | E2A action on CXCR5 | Inhibits | ^166^ |  |
| ID3:Tstim | CXCR5 | Promotes | ^165^ | See their Fig. 4e, f |
| IFN-γ | FAS/FASL | Promotes | ^167^ | IFN-γ signaling is ***required*** for FASL expression |
| IL12R | TBET:ZEB2 | Promotes | ^168,169^ |  |
| IL12R | ID2 | Promotes | ^165^ |  |
| IL12R | BATF | Promotes | ^58^ |  |
| IL12R | FOXO1 | Inhibits | ^64^ |  |
| IL15R | TIGIT | Promotes | ^170^ |  |
| IL21R | ID2 | Promotes | ^165^ |  |
| IL21R | IRF4 | Promotes | ^10^ |  |
| IL21R | TBET:ZEB2 | Promotes | ^171^ |  |
| IL21R | BATF | Promotes | ^10^ |  |
| IL2R | TIGIT | Promotes | ^170^ |  |
| IL2R | BLIMP-1 | Promotes | ^59^ |  |
| IL2R | FOXO1 | Inhibits | ^172^ | Possible mutual repression^100^ |
| IL2R | IRF4 | Promotes | ^173^ |  |
| IL2R | ID2 | Promotes | ^165^ |  |
| IRF4 | NFATC1:AP1:IRF4:BATF | Promotes | ^10,132^ |  |
| LAG3 | Tregs | Promotes | ^174^ |  |
| MAPK | EGR2/3 | Promotes | ^76^ | See their Fig. 2 |
| MAPK | AP1 | Promotes | ^175^ |  |
| MTOR | MYC | Promotes | ^176^ | Reviewed in ^177^ |
| MTOR | HIF1A | Promotes | ^176^ | Reviewed in ^177^. HIF1A protein reaches peak levels by 24hr p.i.^178^.  At later time points (~3 days p.i.^179^), HIF1A is jointly activated by IRF4 + BATF + NFAT^135^. |
| MTOR | IRF4 | Promotes | ^179^ | Reviewed in ^177^ |
| MYC | Glycolysis | Promotes | ^180^ | Reviewed in ^177^ |
| MYC | Proliferation | Promotes | ^180^ | Reviewed in ^177^ |
| MYC | MT biogenesis | Promotes | ^181,182^ |  |
| MYC | FAS/FASL | Promotes | ^154,183,184^ |  |
| NFAT | FAS/FASL | Promotes | ^154,185^ |  |
| NFATC1 | TIM3 | Promotes | ^36^ | NFATC1 and NFATC2 bind the same motif^186^ |
| NFATC1 | LAG3 | Promotes | ^36^ | NFATC1 and NFATC2 bind the same motif^186^ |
| NFATC1 | CTLA4 | Promotes | ^36^ | NFATC1 and NFATC2 bind the same motif^186^ |
| NFATC1 | PD-1 | Promotes | ^36,187^ | NFATC1 and NFATC2 bind the same motif^186^ |
| NFATC1 | NFATC1 | Promotes | ^80,188^ | NFATC1 and NFATC2 bind the same motif^186^ |
| NFATC1 | IRF4 | Promotes | ^135^ | NFATC1 and NFATC2 bind the same motif^186^ |
| NFATC1 | NFATC1:AP1:IRF4:BATF | Promotes | ^135^ |  |
| NFATC1:AP1:IRF4:BATF | MYC | Inhibits | ^189^ |  |
| NFATC1:AP1:IRF4:BATF | NFATC1 | Promotes | ^135^ |  |
| NFATC1:AP1:IRF4:BATF | BCL6 | Inhibits | ^69^ |  |
| NFATC2 | CTLA4 | Promotes | ^190^ | NFATC1 and NFATC2 bind the same motif^186^ |
| NFATC2 | PD-1 | Promotes | ^36^ |  |
| NFATC2 | NFATC1 | Promotes | ^80^ | See also ^191^ |
| NFATC2 | IL12R | Inhibits | ^192^ |  |
| NFATs | BCL6 | Promotes | ^193^ | Indirect, via Bob1 |
| NF-κB | BTLA | Promotes | ^193^ | Indirect, via Bob1 |
| NF-κB | BCL6 | Promotes | ^193^ | Indirect, via Bob1 |
| NR4A1 | FAS/FASL | Promotes | ^144^ |  |
| PD-1 | CD3, TCR, CD8, CD28 | Inhibits | ^89,194^ |  |
| PGC1A | MT biogenesis | Promotes | ^5,195,196^ |  |
| PI3K | AKT | Promotes | ^197^ |  |
| RAS | MAPK | Promotes | ^198^ |  |
| RUNX3 | GZMA/B | Promotes | ^155^ |  |
| RUNX3 | PRF1 | Promotes | ^155^ |  |
| RUNX3 | IFN-γ | Promotes | ^155^ |  |
| RUNX3 | EOMES | Promotes | ^155^ |  |
| TBET | ZEB2 | Promotes | ^115^ |  |
| TBET:ZEB2 | GZMA/B | Promotes | ^115^ |  |
| TBET:ZEB2 | IFN-γ | Promotes | ^115^ |  |
| TBET:ZEB2 | PD-1 | Inhibits | ^103,115^ |  |
| TCF-1 | KLRG1 | Inhibits | ^73^ |  |
| TCF-1 | BCL6 | Promotes | ^199,200^ |  |
| TIGIT | degranulation | Inhibits | ^201^ |  |
| TIGIT | Tregs | Promotes | ^202^ |  |
| TIM3 | CD3, TCR, CD8, CD28 | Inhibits | ^203^ |  |
| ZEB2 | ID3 | Inhibits | ^115^ | Repression may be mutual^165^ |
| ZEB2 | TBET | Promotes | ^115^ |  |

##### **Supplementary Table 3. Timings of TCE network interactions**

| **Source** | **Target** | **Interaction** | **Reference** | **Comment** |
| --- | --- | --- | --- | --- |
| 4-1BB | FAS/FASL | Promotes | ^204,205^ | 4-1BB expression is delayed, peaking at about 1-5 days p.i.^116,117^, and its action on FAS signaling is gated by IFN-γ^167^. |
| AKT | FOXO1 | Inhibits | ^64,206^ | FOXO1 protein is high in naïve cells and down-regulated over 2–3 days post stim. |
| EZH2 | FOXO1, TCF-1 | Inhibits | ^207^ | EZH2 expression peaks at 1 day post stim in CD8+ cells |
| EZH2 | TCF-1 | Inhibits | ^44^ | H3K27me3 of TCF-1 locus peaks at ~ 10 days post-infection. |
| DNMT3A | Downstream of EZH2 activity | Inhibits | ^21^ | DNMT3A is activated by TCR stim. and peaks at ~24 hours. |
| DNMTs | TCF-1 | Inhibits | ^30,208^ | TCF-1 locus is methylated after day 4 and by ~ day 8 post-infection. |

#### **Supplementary References (for the Supplementary Tables)**

54.	Au-Yeung, N., Mandhana, R. & Horvath, C. M. Transcriptional regulation by STAT1 and STAT2 in the interferon JAK-STAT pathway. *JAKSTAT* **2,** e23931 (2013).

127.	Tullai, J. W., Tacheva, S., Owens, L. J., Graham, J. R. & Cooper, G. M. AP-1 is a component of the transcriptional network regulated by GSK-3 in quiescent cells. *PLoS One* **6,** e20150 (2011).

189.	Mognol, G. P., Carneiro, F. R., Robbs, B. K., Faget, D. V. & Viola, J. P. Cell cycle and apoptosis regulation by NFAT transcription factors: new roles for an old player. *Cell Death Dis.* **7,** e2199 (2016).
