## Supplemental Figures for "Integrative network modeling reveals mechanisms underlying T cell exhaustion"

**Supplementary Fig. 1.** Diverse mechanisms can lead to a failure of $CD8^+$ T cells to respond to tumors. In the scenarios to the left of the figure, $CD8^+$ T cells are either not activated (e.g. due to a dearth of antigens or spatial exclusion of T cells), or they may be unable to respond due to extrinsic suppression. In senescence, anergy and exhaustion on the other hand, $CD8^+$ T cells are activated but hypofunctional.

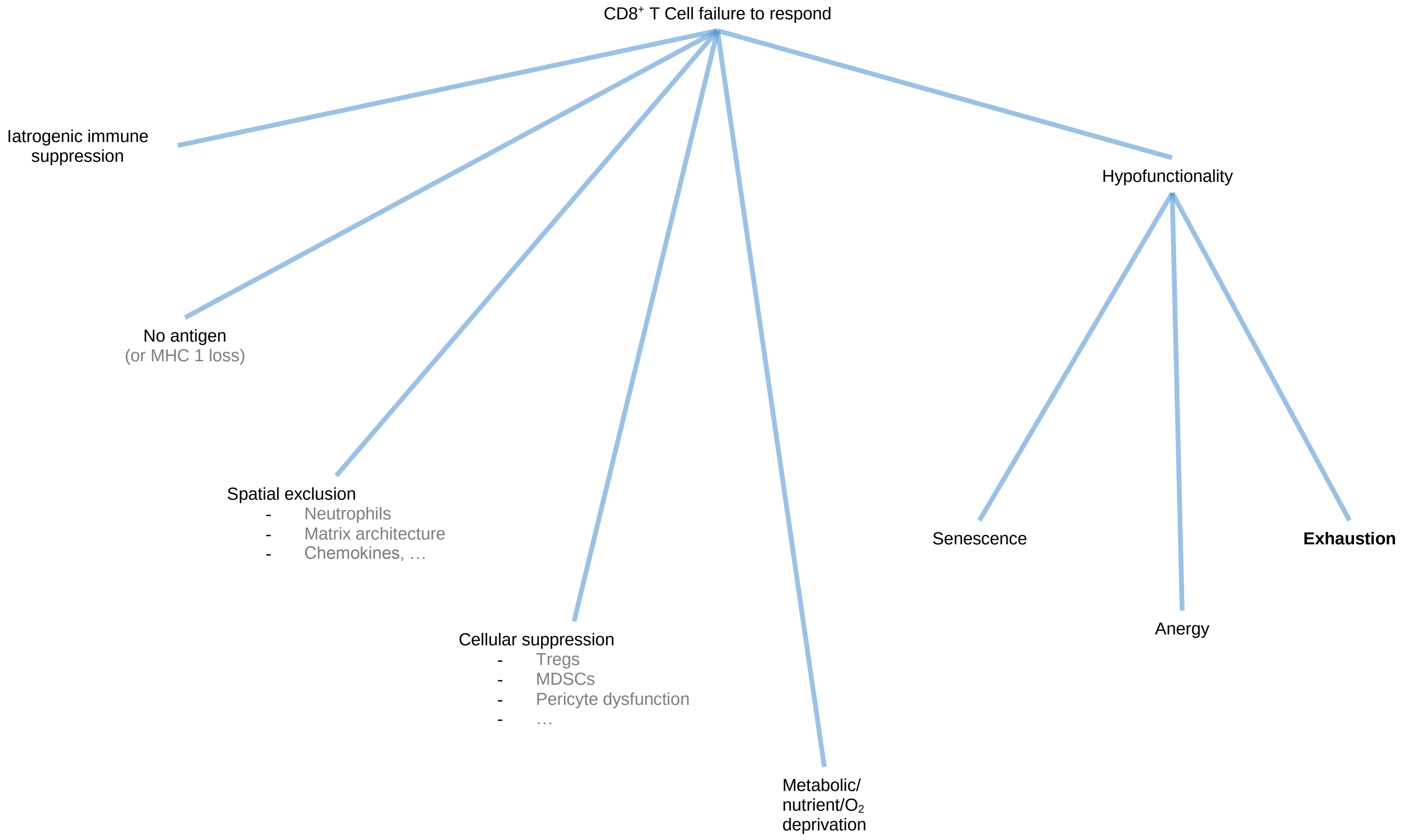

**Supplementary Fig. 2.** Literature-based network of molecular interactions underlying CD8$^{+}$ T cell exhaustion (TCE), comprising 505 interactions and 113 genes (119 nodes). For simplicity, each node in the network represents the overall activity of a gene *and* its product(s). Edges (connecting lines) represent reported interactions. Edges ending in a blue arrowhead indicate that the source node 'promotes' the target node. Edges ending in red bars denote repression. Orange labels and bounding boxes mark groups of related genes/gene products.

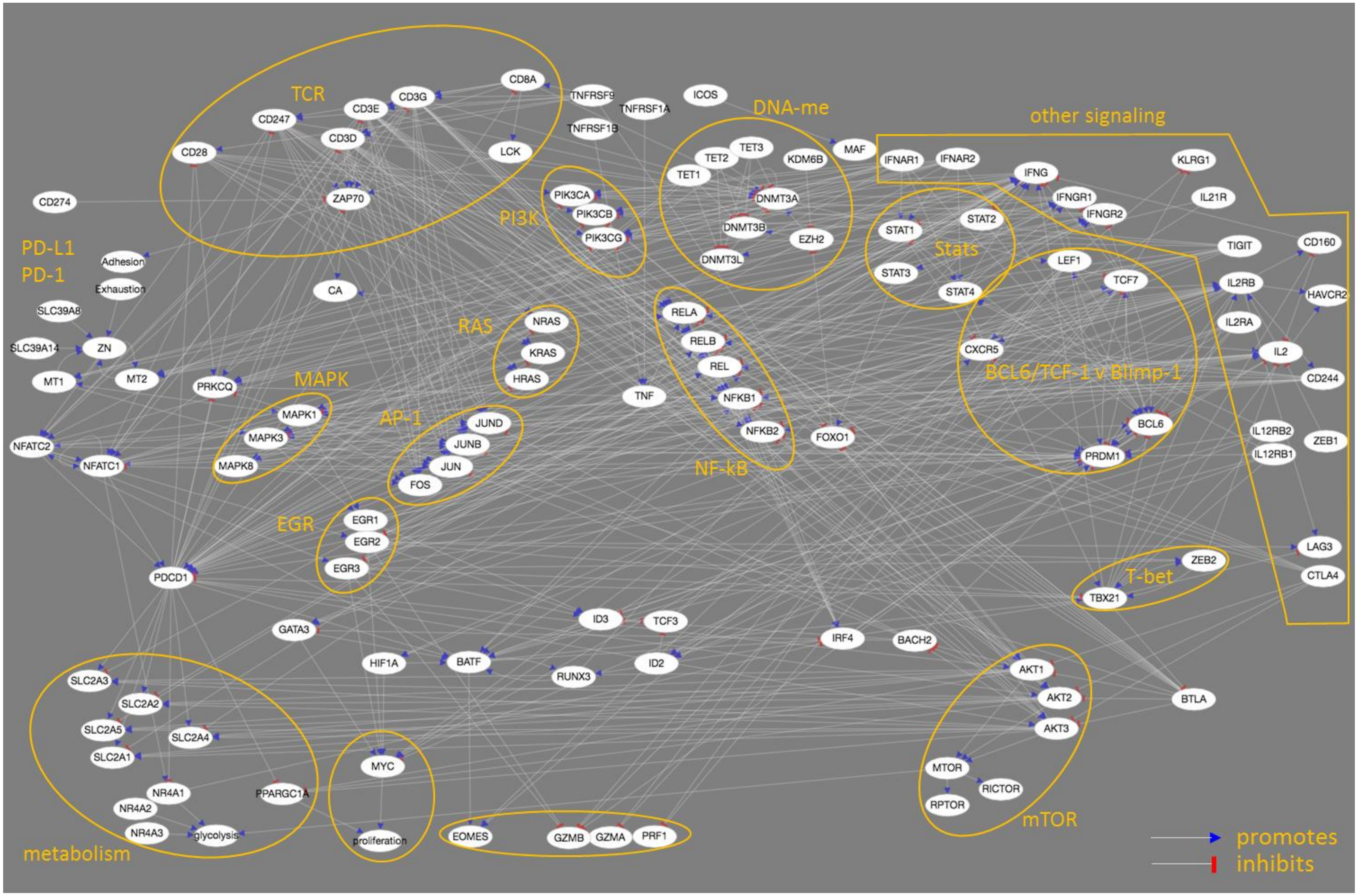

**Supplementary Fig. 3.** Superposition of expression data onto the TCE network highlights time/condition dependence of interactions. Node colors indicate mRNA expression relative to naïve CD8$^{+}$ cells (see color bar at top left). Red edges are inhibitory. Blue edges are promoting. Edge thickness indicates the fraction of replicates in which the source and target gene expression are concordant with the sense of edge (see key at bottom right). Day 5, acute infection data from GSE89307, Schietinger lab, 2017.

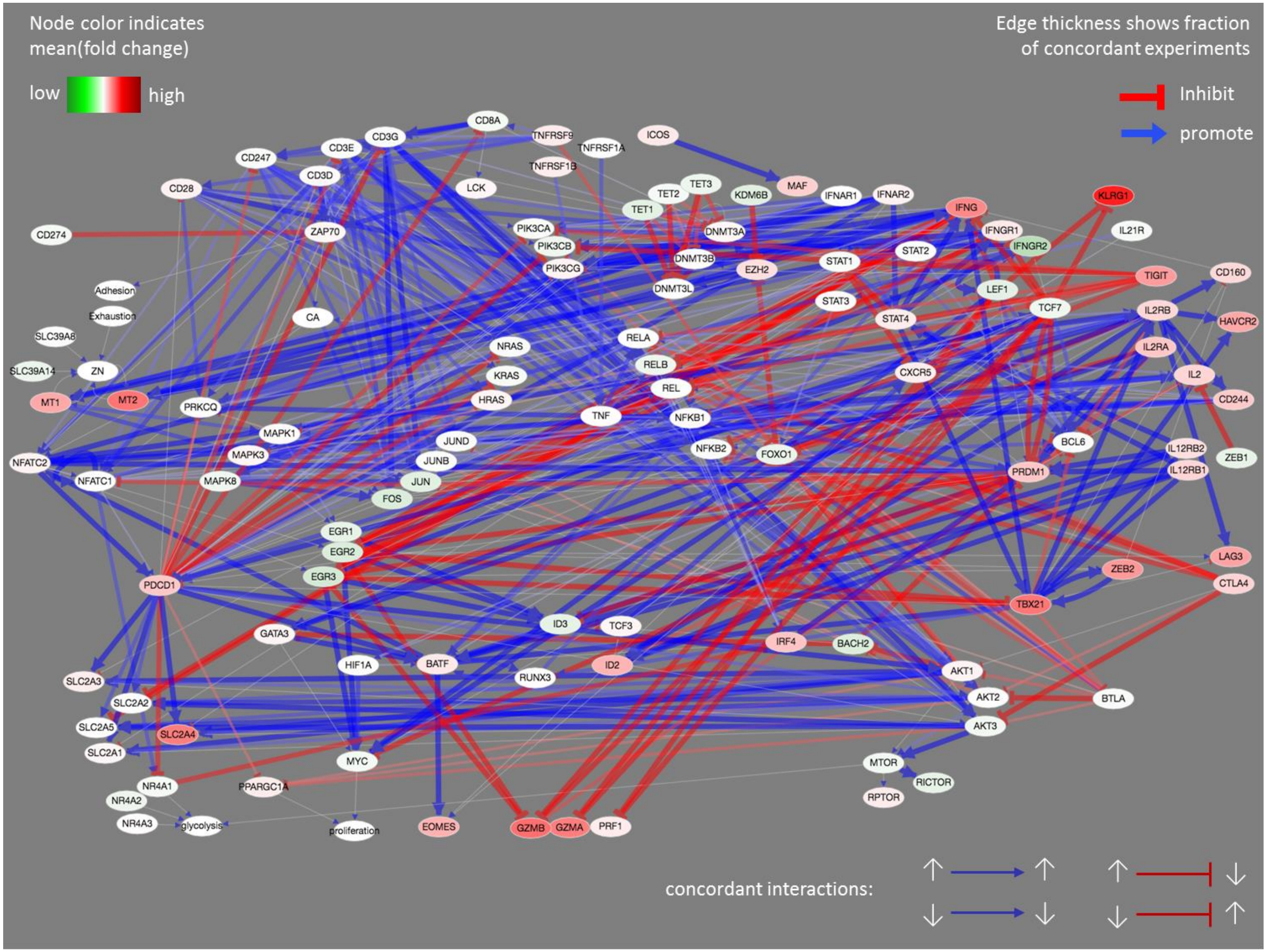

**Supplementary Fig. 4**. Superposition of expression data onto the TCE network highlights time/condition dependence of interactions. Node colors indicate mRNA expression relative to naïve CD8$^+$ cells (see color bar at top left). Red edges are inhibitory. Blue edges are promoting. Edge thickness indicates the fraction of replicates in which the source and target gene expression are concordant with the sense of edge (see key at bottom right). Day 7, acute infection data from GSE89307, Schietinger lab, 2017.

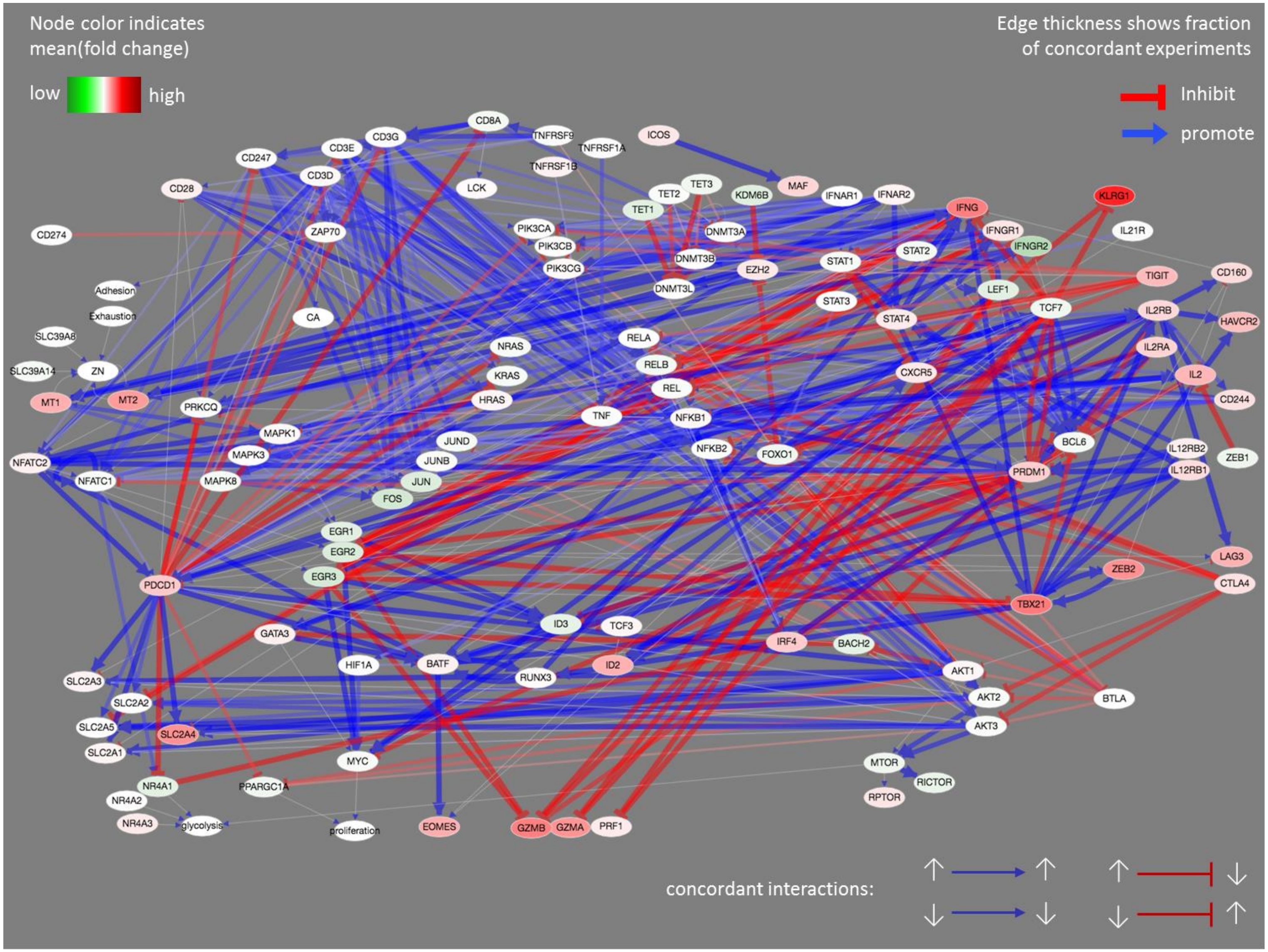

**Supplementary Fig. 5**. Superposition of expression data onto the TCE network highlights time/condition dependence of interactions. Node colors indicate mRNA expression relative to naïve CD8$^{+}$ cells (see color bar at top left). Red edges are inhibitory. Blue edges are promoting. Edge thickness indicates the fraction of replicates in which the source and target gene expression are concordant with the sense of edge (see key at bottom right). Day 5, tumor data from GSE89307, Schietinger lab, 2017.

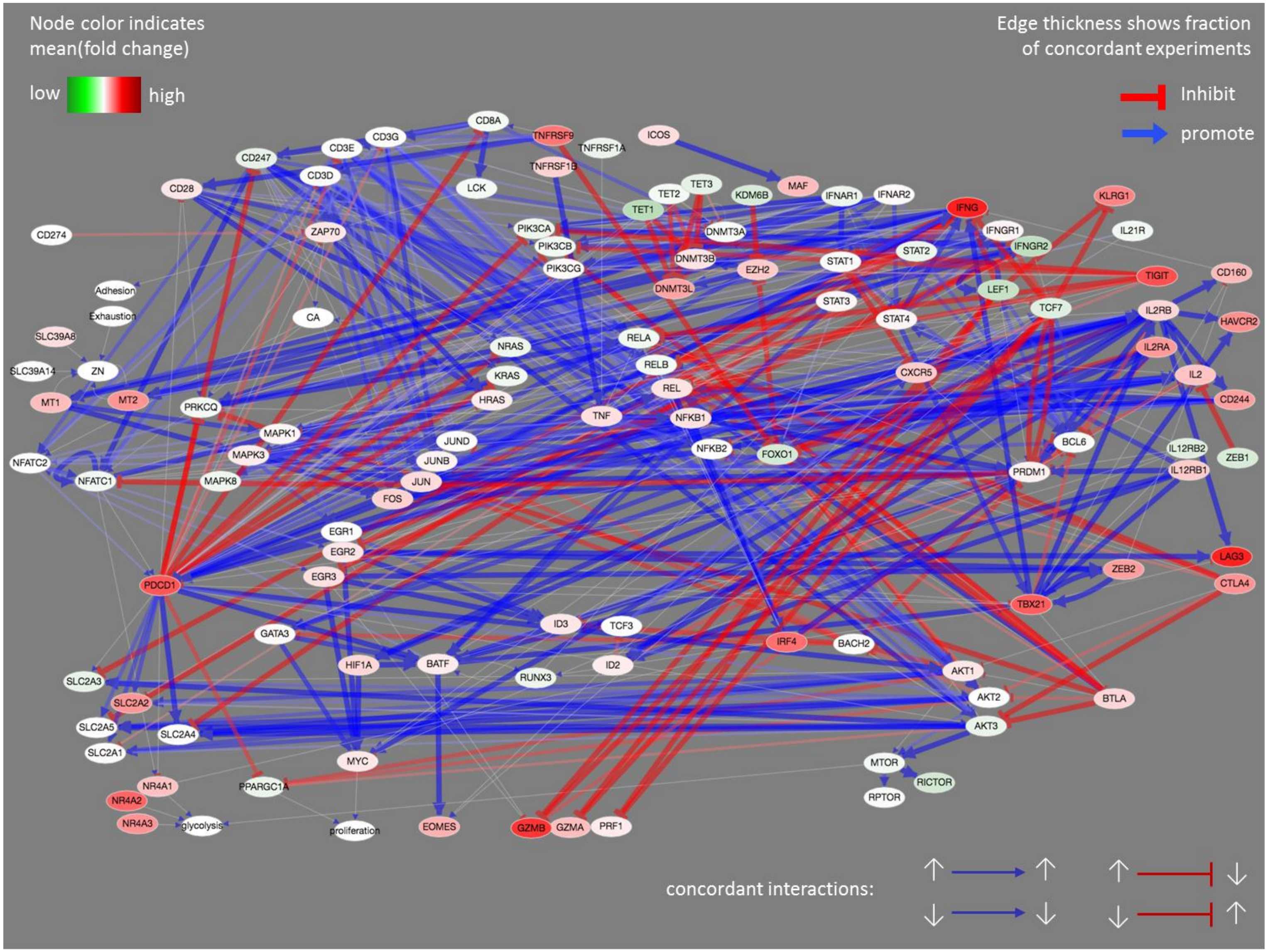

**Supplementary Fig. 6**. Superposition of expression data onto the TCE network highlights time/condition dependence of interactions. Node colors indicate mRNA expression relative to naïve CD8$^{+}$ cells (see color bar at top left). Red edges are inhibitory. Blue edges are promoting. Edge thickness indicates the fraction of replicates in which the source and target gene expression are concordant with the sense of edge (see key at bottom right). Day 7, tumor data from GSE89307, Schietinger lab, 2017.

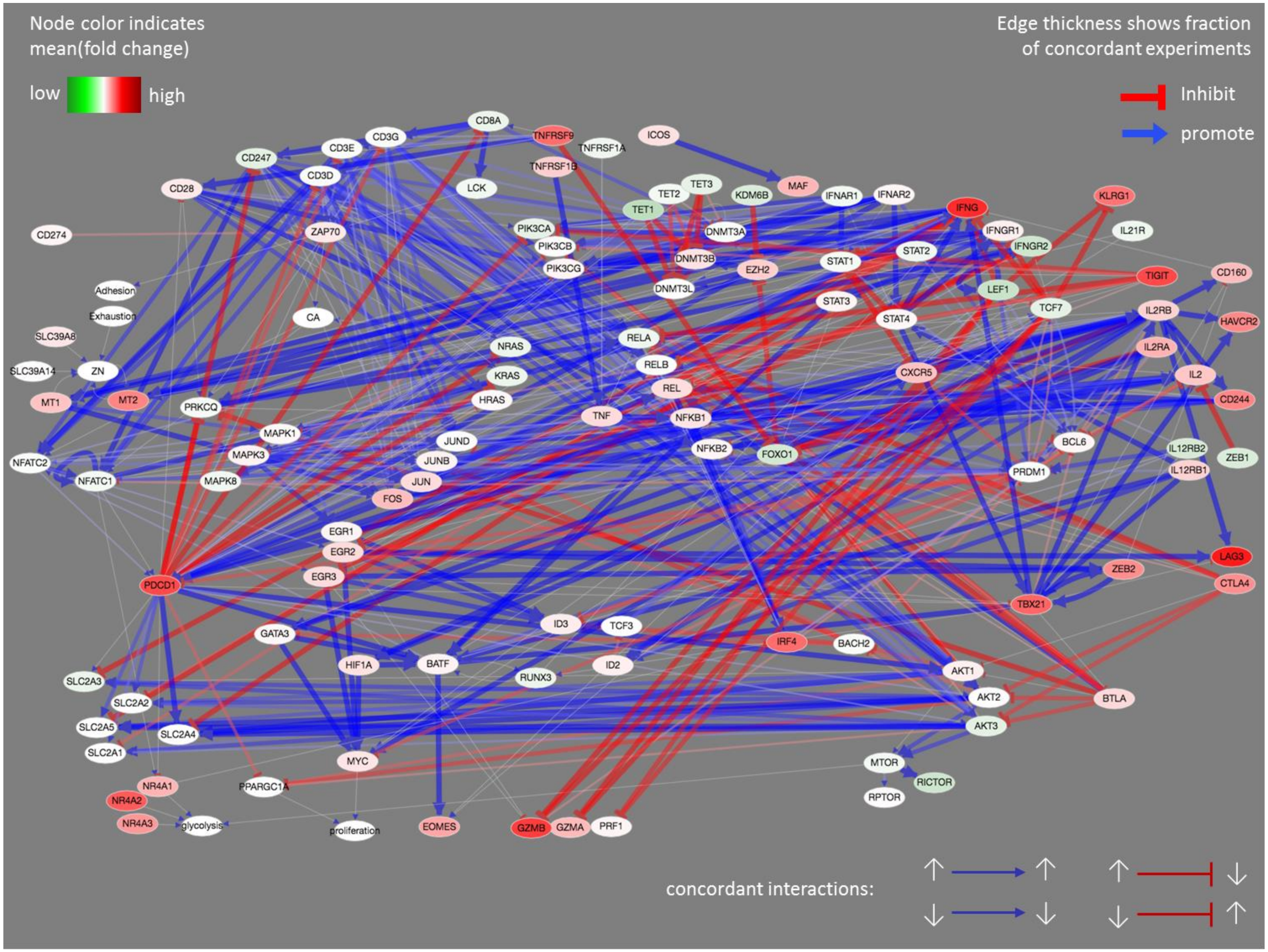

**Supplementary Fig. 7**. Superposition of expression data onto the TCE network highlights time/condition dependence of interactions. Node colors indicate mRNA expression relative to naïve CD8$^{+}$ cells (see color bar at top left). Red edges are inhibitory. Blue edges are promoting. Edge thickness indicates the fraction of replicates in which the source and target gene expression are concordant with the sense of edge (see key at bottom right). Day 14, tumor data from GSE89307, Schietinger lab, 2017.

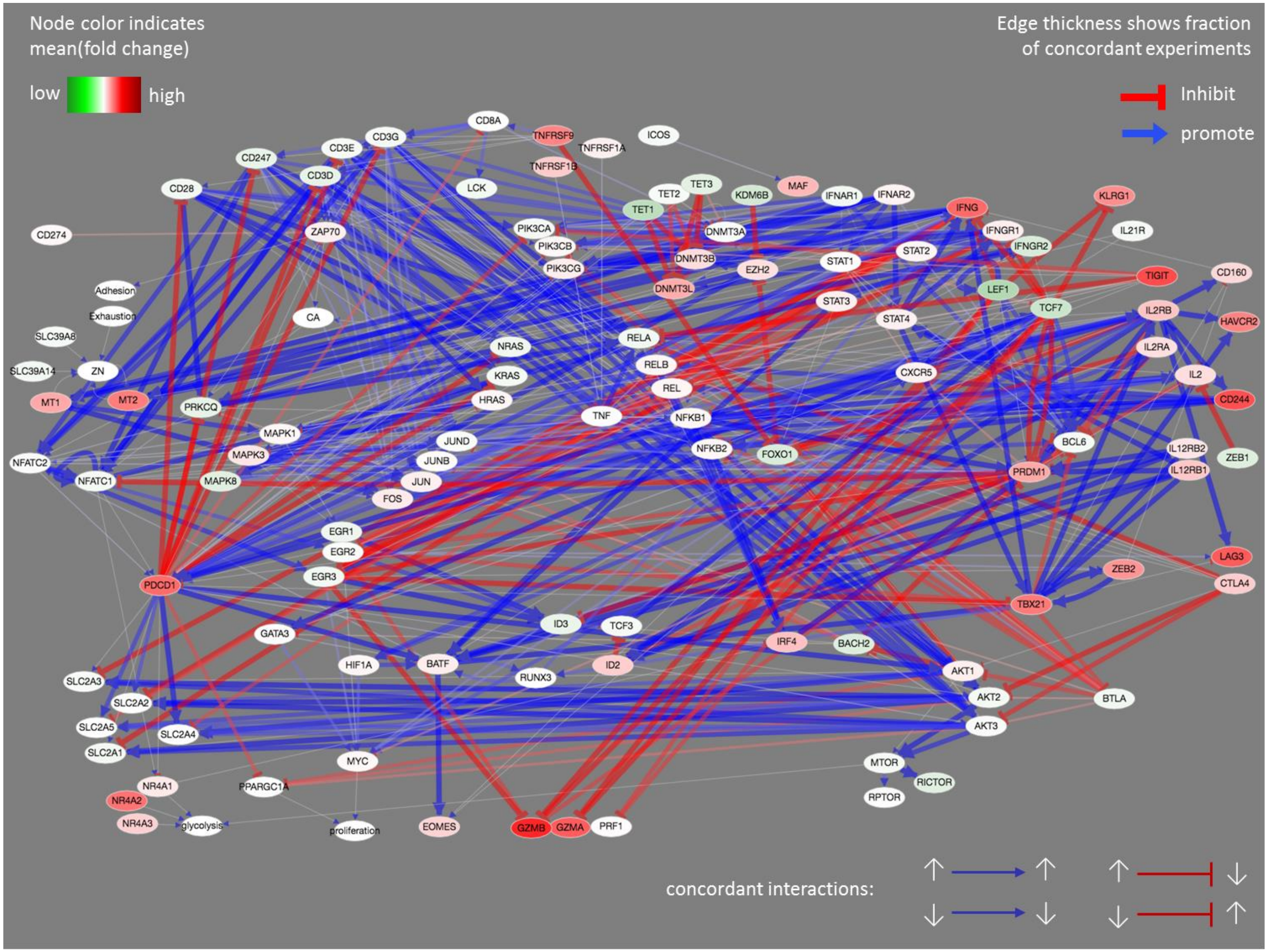

**Supplementary Fig. 8**. Superposition of expression data onto the TCE network highlights time/condition dependence of interactions. Node colors indicate mRNA expression relative to naïve CD8$^{+}$ cells (see color bar at top left). Red edges are inhibitory. Blue edges are promoting. Edge thickness indicates the fraction of replicates in which the source and target gene expression are concordant with the sense of edge (see key at bottom right). Day 21, tumor data from GSE89307, Schietinger lab, 2017.

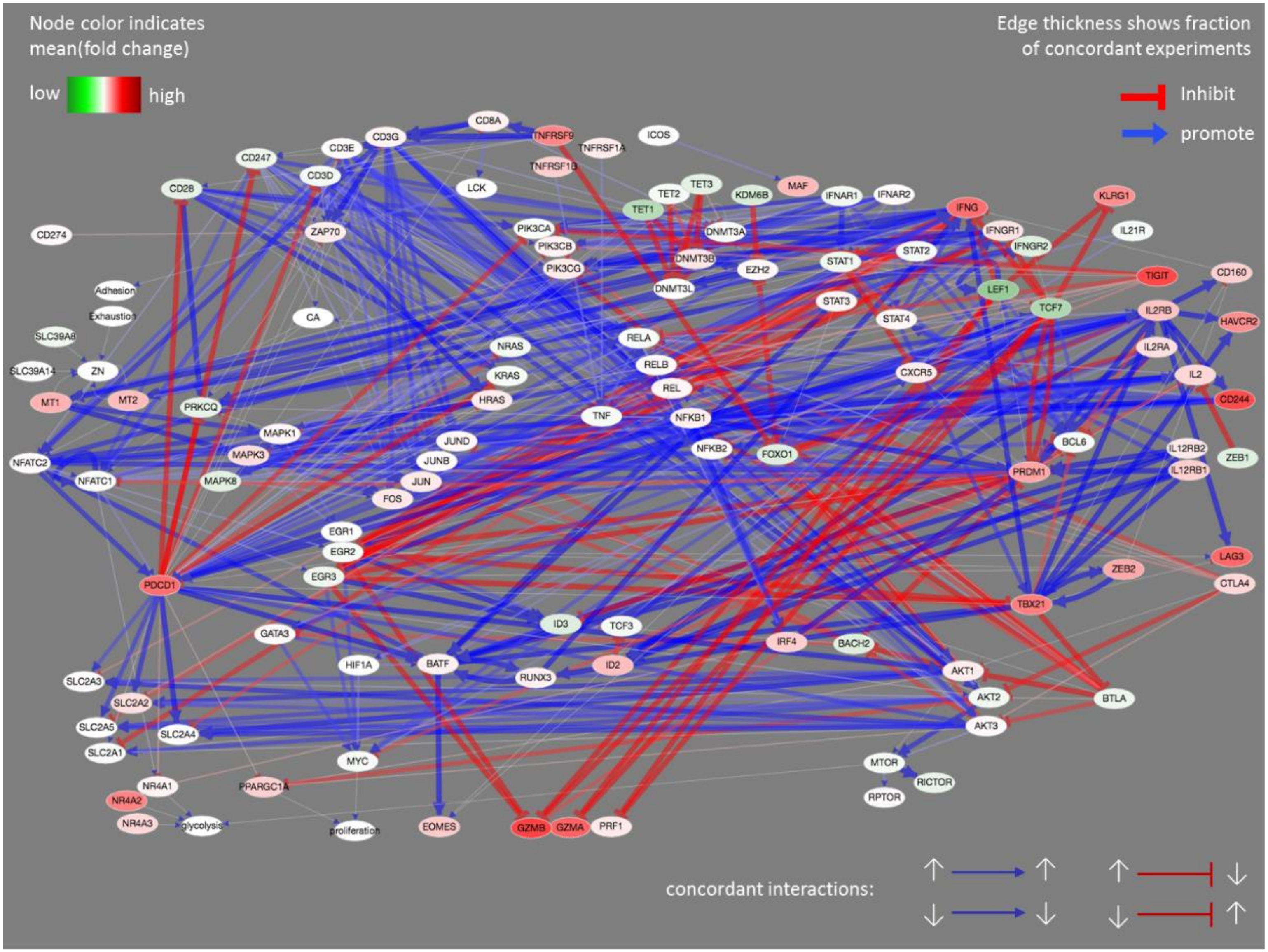

**Supplementary Fig. 9**. Genes and interactions in our TCE network are consistent with experimental data. The heatmap at left shows edges (rows) in which expression fold change with respect to naïve cells in GSE89307 is concordant (red) or not (blue) with the sense of the edge. 17 TCE-network edges (3.6% of 478 gene-gene edges) were not concordant in any condition, listed below in the format ‘source:target:edgeSense’:

```
 [1] AKT1:MTOR:1        BATF:IRF4:-1        CD160:IFNG:-1       CD160:IL2:-1
 [5] CTLA4:AKT1:-1      DNMT3A:DNMT3A:-1    FOXO1:PDCD1:1       IFNG:SLC2A4:-1
 [9] LEF1:EOMES:1       TBX21:CD160:-1      TBX21:LAG3:-1       TBX21:PDCD1:-1
[13] TCF7:EOMES:1       TCF7:IFNGR2:-1      TET1:PDCD1:1        TET3:PDCD1:1
[17] ZEB2:IL2:-1
```

Right panel: using the same data and 500,000 randomly assigned edges, concordance scores of 1 occur at a False Discovery Rate of ~15%.

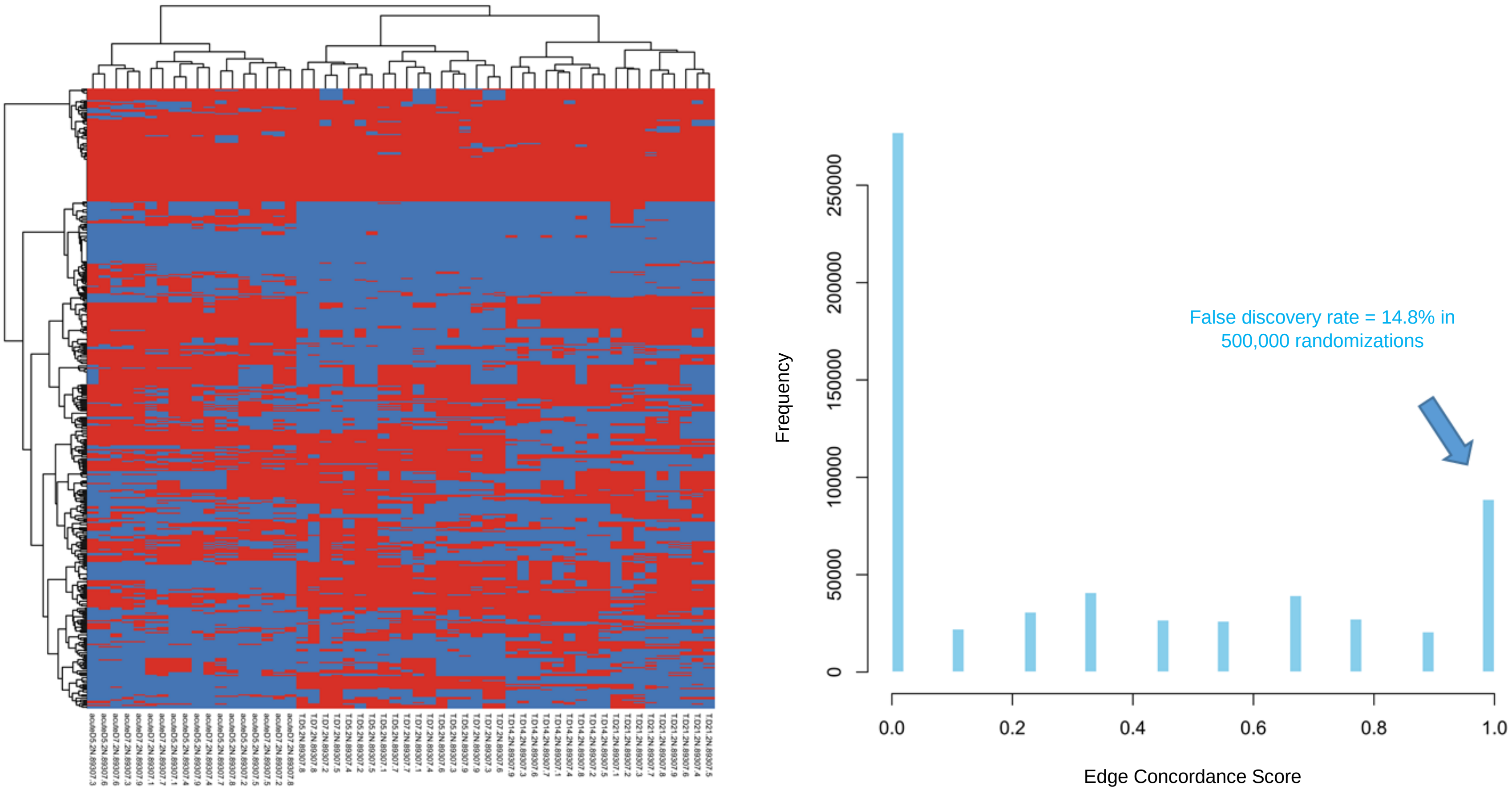

**Supplementary Fig. 10**. The TCE network genes track $CD8^+$ evolution during both acute and chronic stimulation. Example PCA plots showing data from GSE89307 (mouse liver tumor $CD8^+$ tumor-infiltrating lymphocytes). The top row shows principal component analysis (PCA) plots using the TCE network genes (left) and 2 published gene sets, as indicated. In all 3 plots, acute (shades of blue) and chronic (yellow to red) state transitions have similar trajectories and are monotonic with respect to time. The lower 3 plots confirm these observations using 3 alternate gene sets, as indicated.

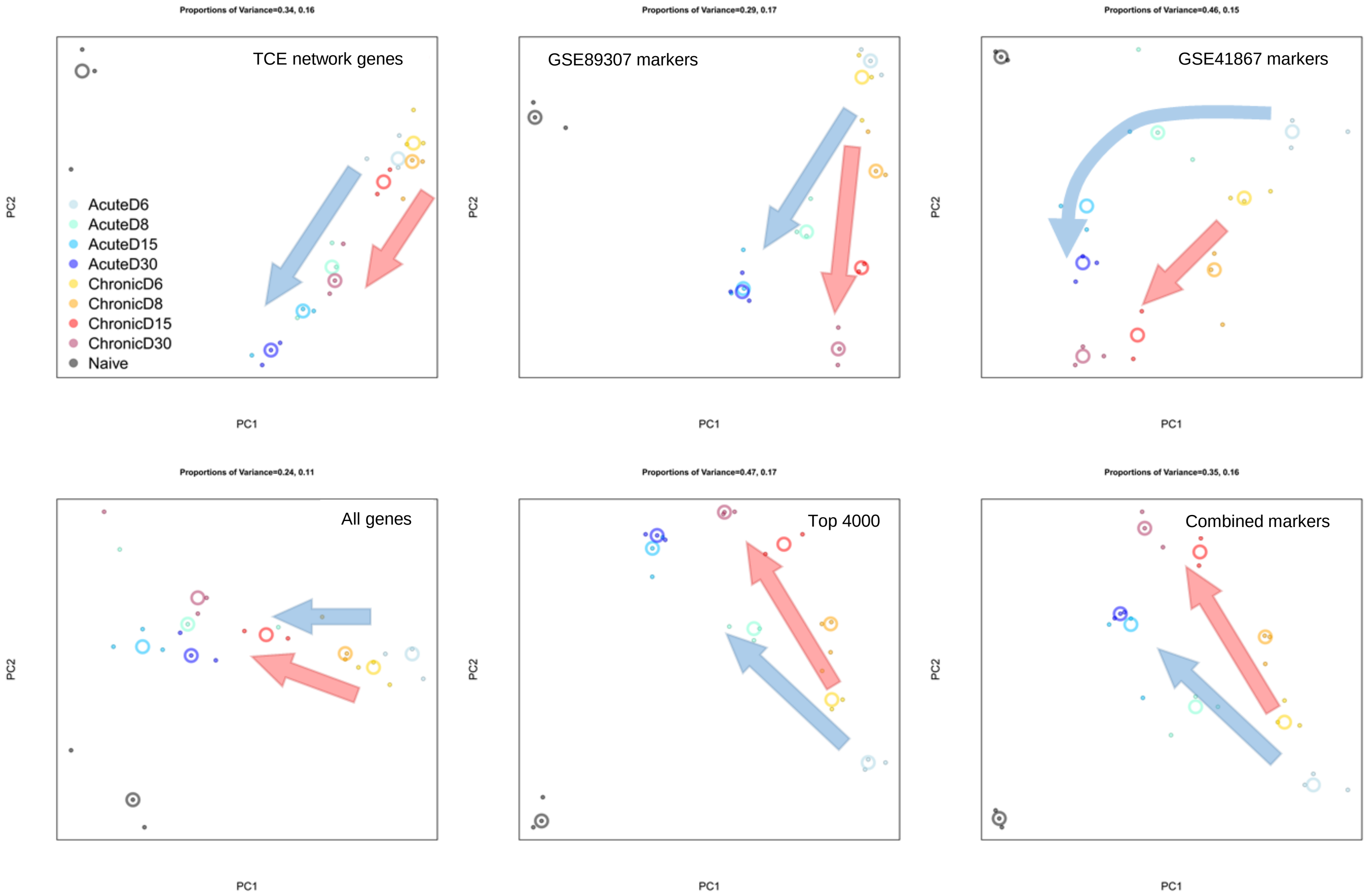

**Supplementary Fig. 11**. Acute and chronic stimulation of CD8⁺ cells show similar metabolic profiles. Metabolic pathway activity scores were generated as described in Methods. The heatmap on the left is for GSE89307 (mouse liver tumor CD8⁺ tumor-infiltrating lymphocytes). Columns are CD8⁺ cells in various states: N = naïve. E5 and E7 = acute infection at days 5 and 7. TDx = tumor CD8⁺ cells at day 'x', as indicated. Areas overlaid in gray show memory cell states. The heat map on the right shows metabolic pathway activities of acutely stimulated CD8⁺ cells for GSE15907 (the Immunological Genome Project). 'Nve' marks naïve cells. 'VSV' = vesicular stomatitis virus. 'Lis' = *Listeria monocytogenes*. Days and hours post-infection are indicated by 'd' and 'hr'.

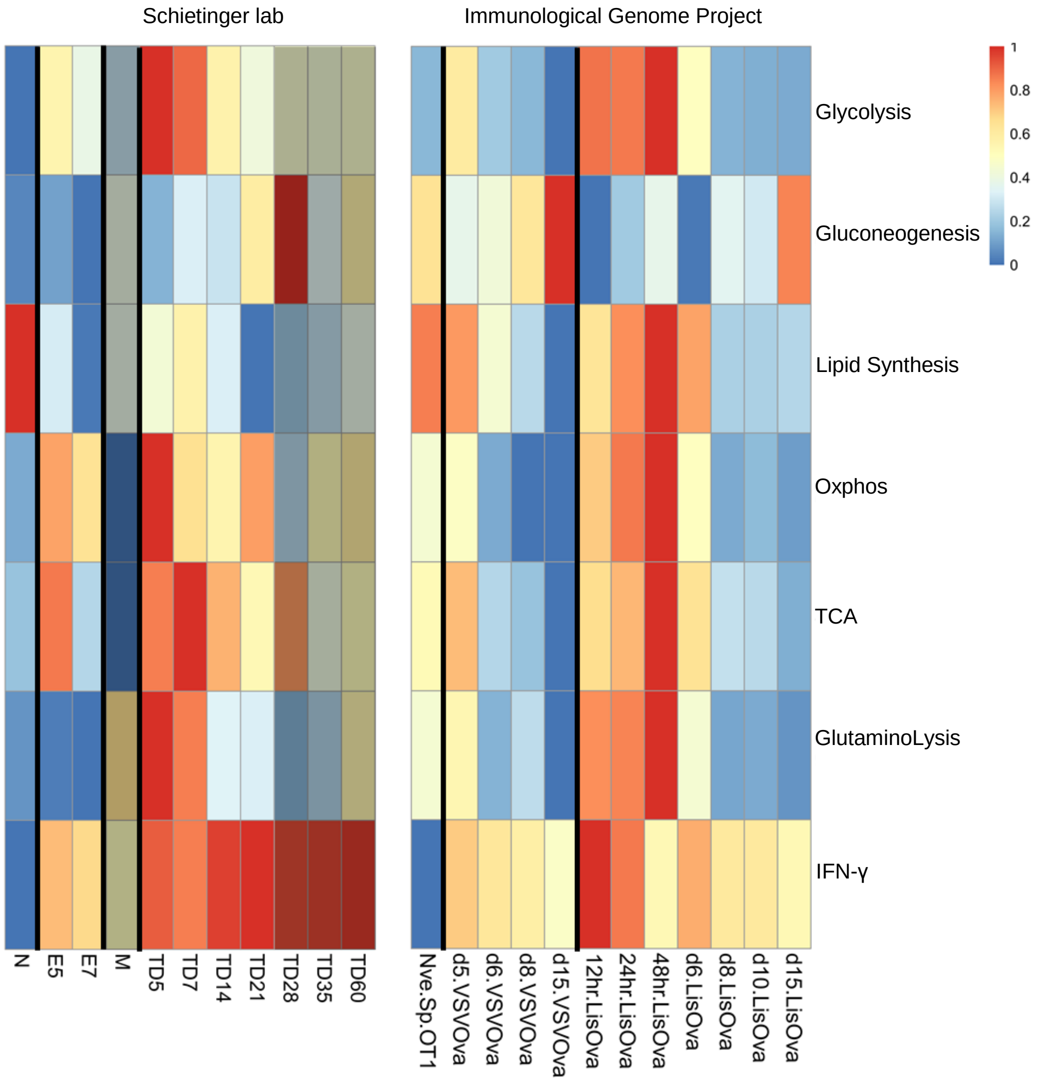

**Supplementary Fig. 12**. Changes in metabolic gene activity track CD8+ TCE network state changes during both acute and chronic CD8+ T cell stimulation. Principal component analysis plots for 2 data sets are shown as examples. Arrows indicate the direction of change over time. See Methods for a list of the metabolic marker genes used.

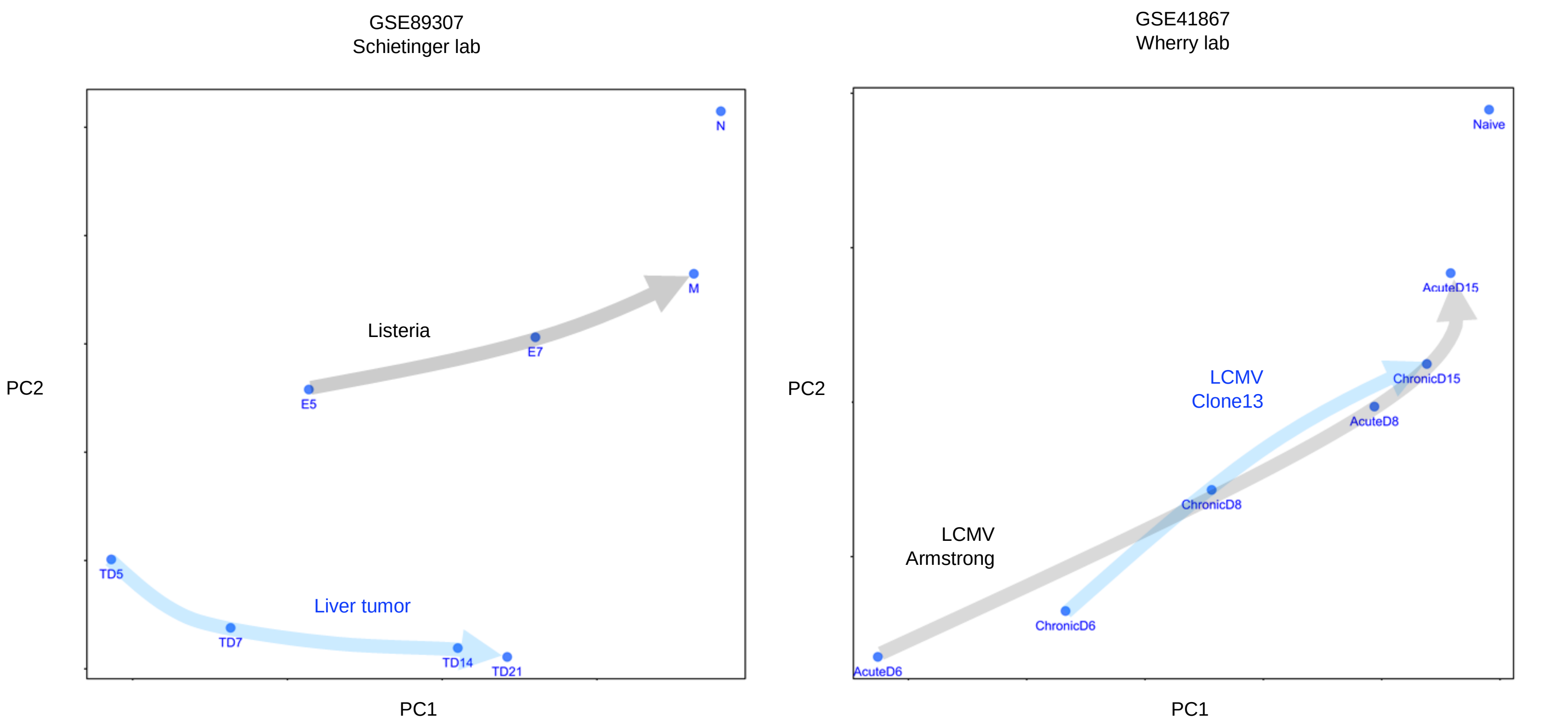

**Supplementary Fig. 13**. Co-expression gene clusters for GSE89307 (Schietinger lab mouse liver tumor CD8$^+$ T cells). Example clusters showing up(down)-regulation between days 7 and 14 following tumor initiation ('early' and 'late' activity phases) are highlighted in blue (yellow) background.

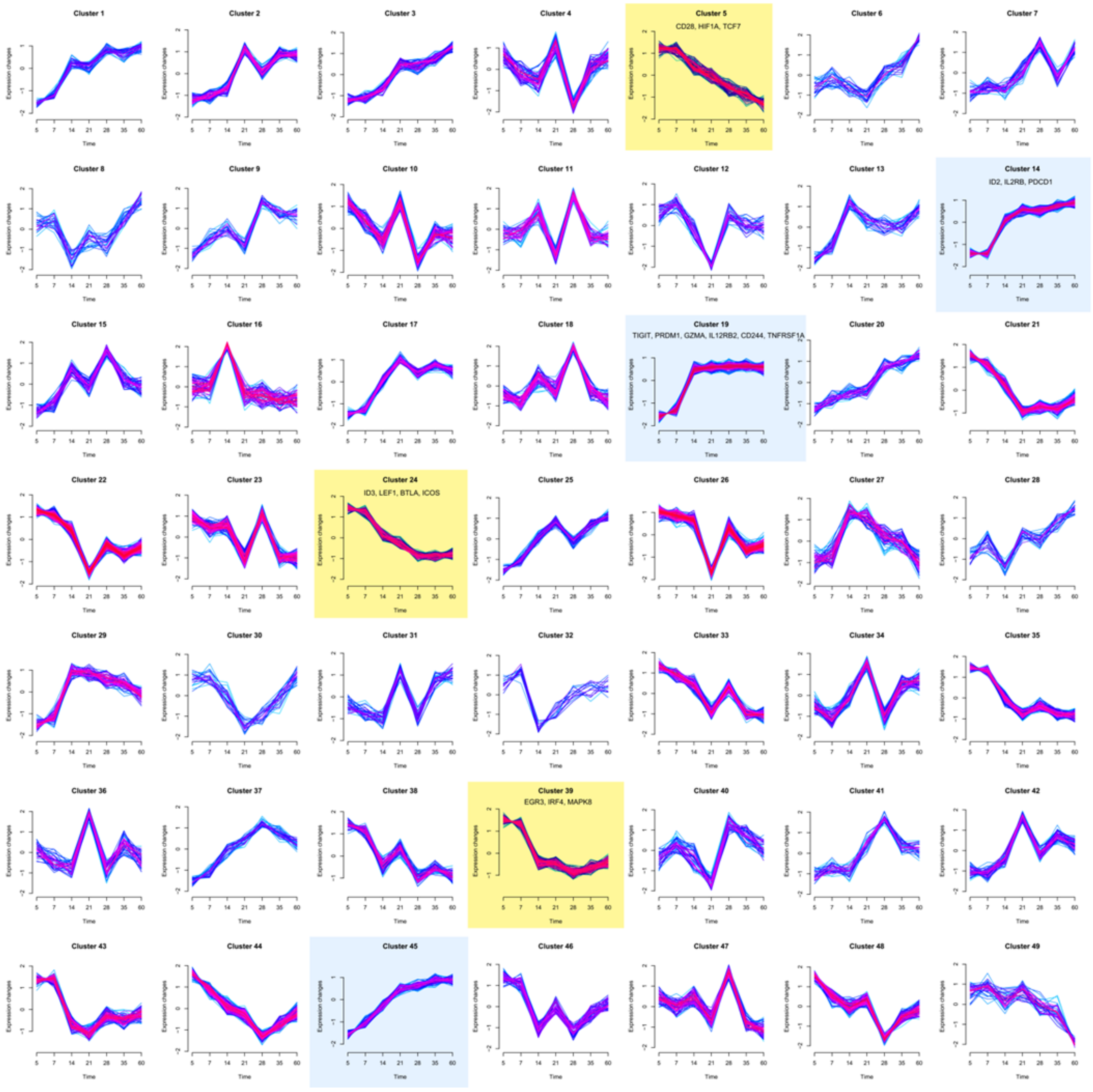

**Supplementary Fig. 14**. Expression clusters of *metabolic* genes in tumor-infiltrating lymphocytes (GSE89307, Schietinger lab, 2017). Green background marks genes changing between days 7 and 14 ('early' and 'late' phases). Clusters containing the 3 genes discussed in Supplementary Fig. 16 are marked in red.

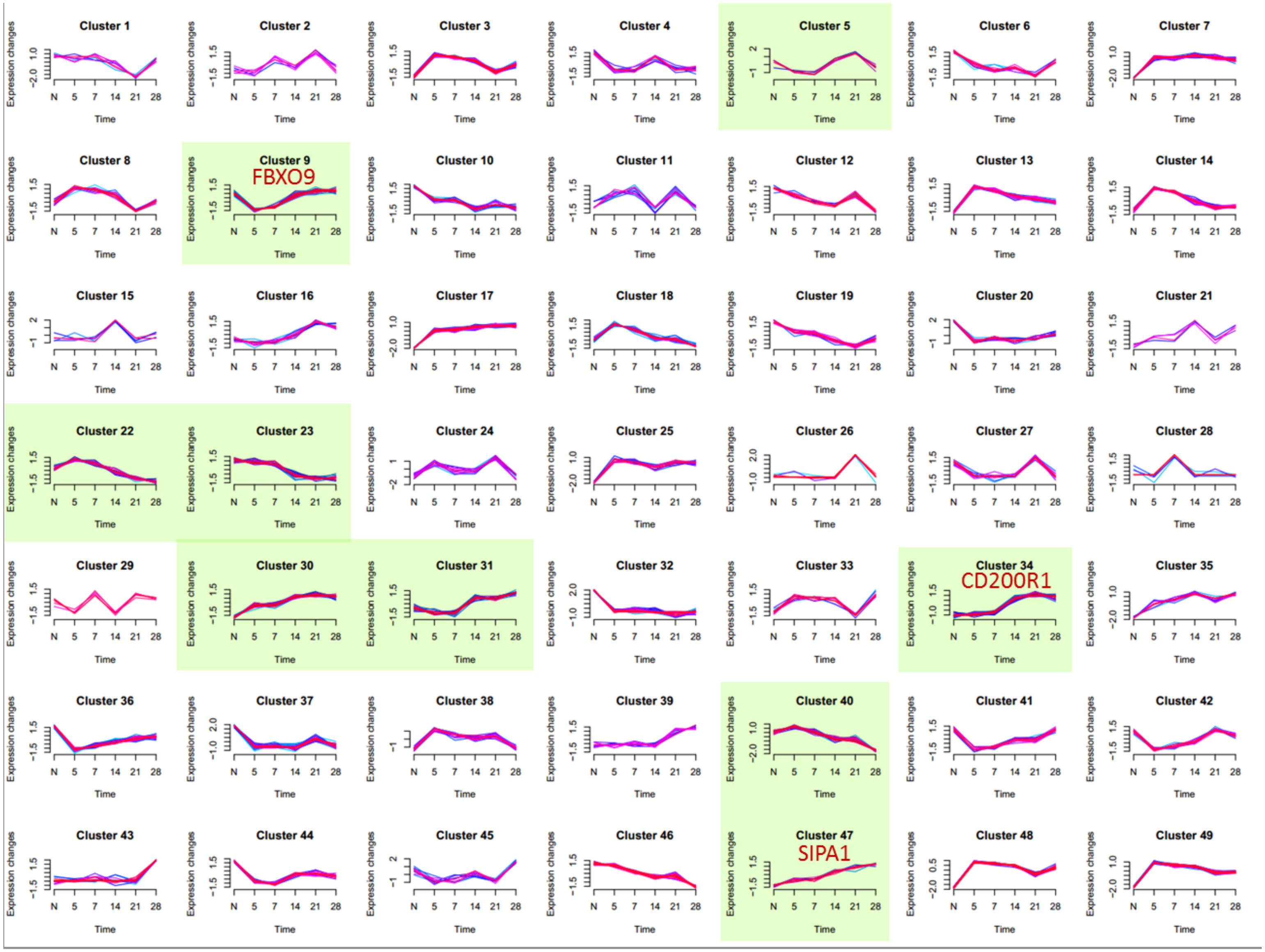

**Supplementary Fig. 15.** Expression clusters of *metabolic* genes in chronic LCMV (GSE41867, Wherry lab, 2012). Green background marks genes changing between days 6 and 15 (‘early’ and ‘late’ phases). Clusters containing the 3 genes discussed in Supplementary Fig. 16 are marked in red.

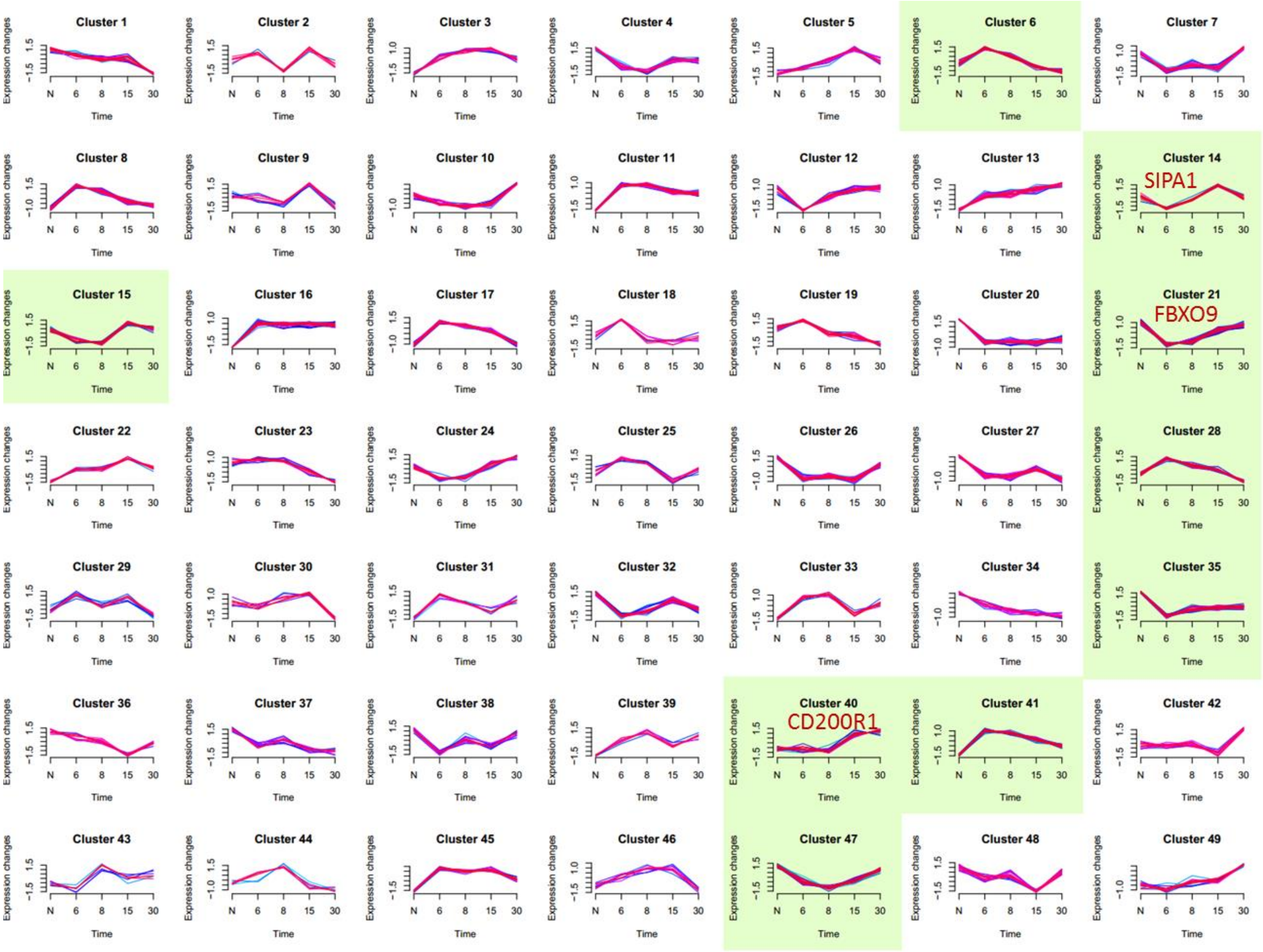

**Supplementary Fig. 16**. Active-late, chronic-only, metabolic-cluster genes that are differently clustered in acute response (all 3 genes are TP53 regulated). CD200R1 is known to repress ERK1/2 and IFN□[1], and confers tolerance[2]. FBXO9 suppresses mTORC1 and cell proliferation[3]. SIPA1 suppresses RAS signaling and proliferation[4]. Of the 3 genes, only CD200R1 shows high absolute fold changes in both chronic infection and tumor settings, but not in acute infection (right hand panels).

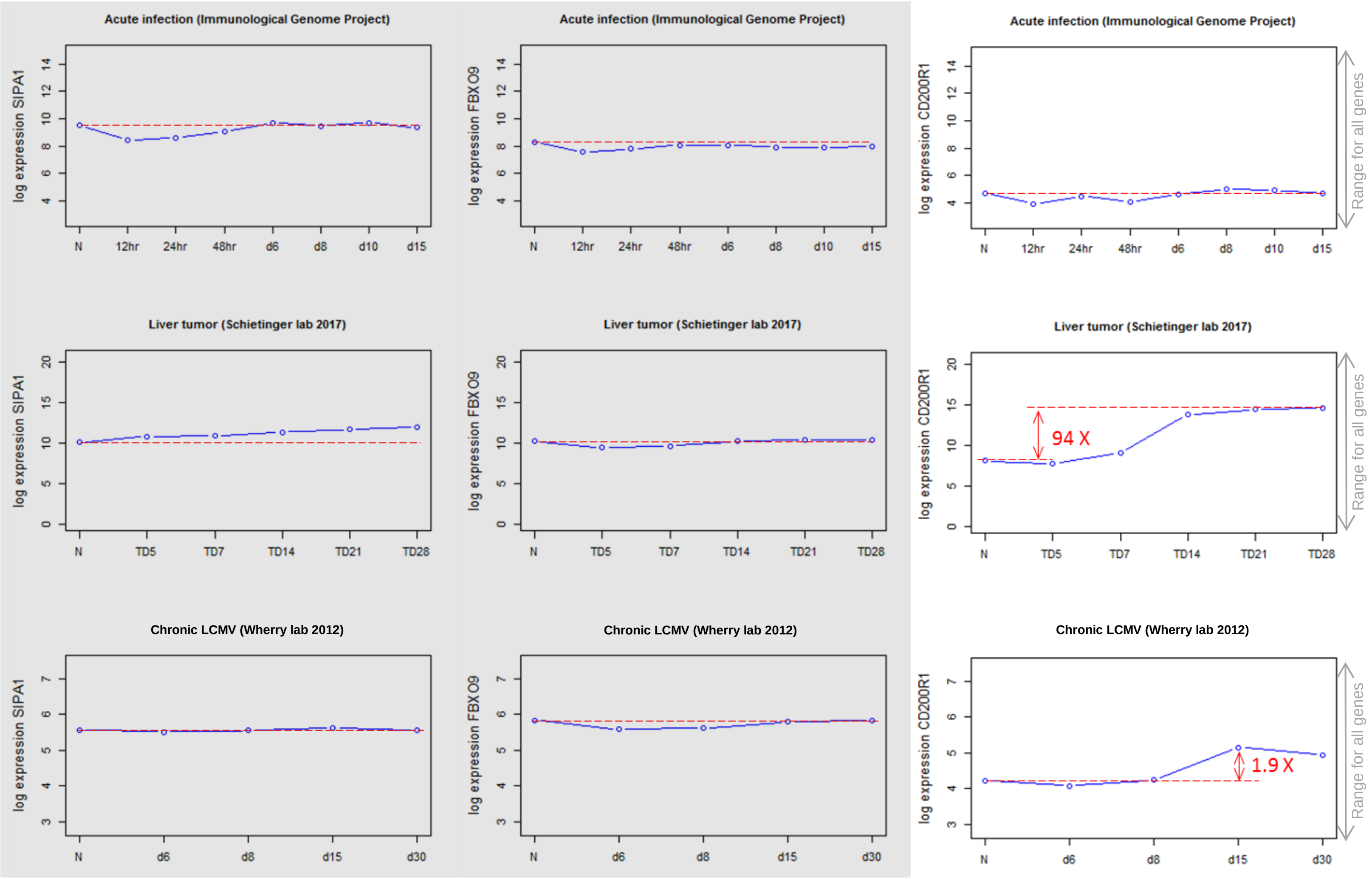

**Supplementary Fig. 17**. Initial (starting) state of an example Boolean logic model (see Methods for model equations). Red nodes are on. The remaining nodes are off.

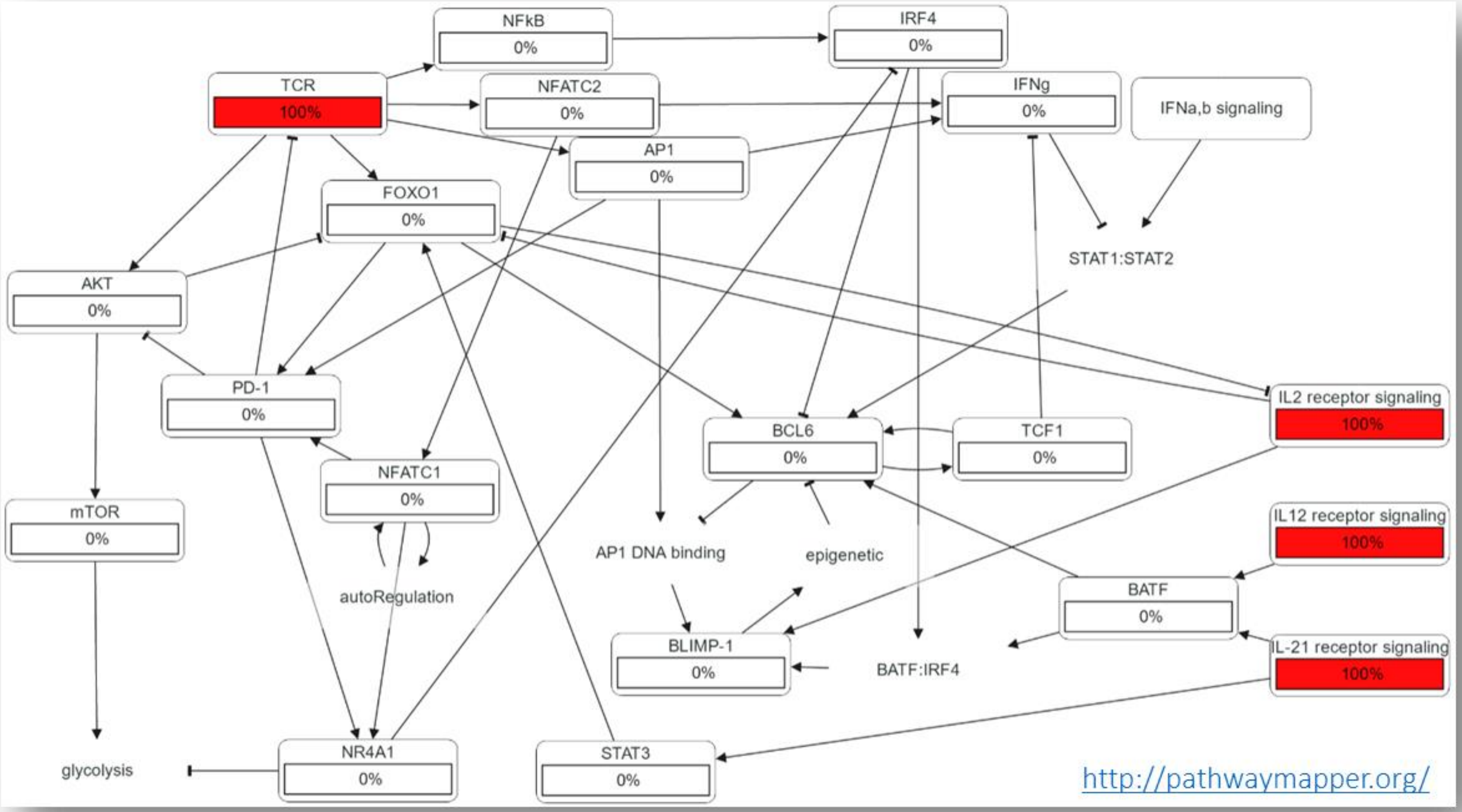

**Supplementary Fig. 18**. Early acute response state of the example Boolean logic model. Red nodes are on. The remaining nodes are off.

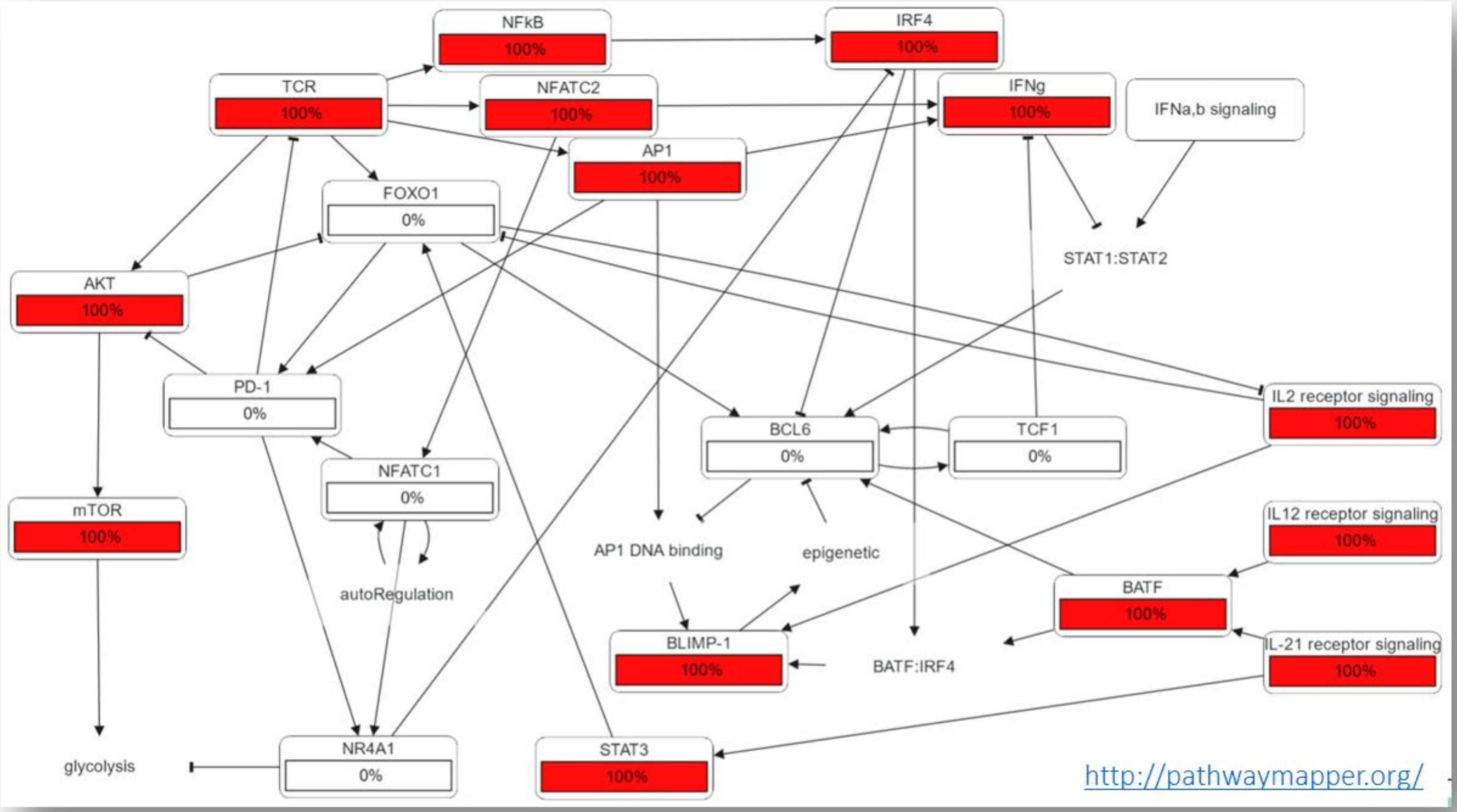

**Supplementary Fig. 19**. Late acute response state of an example Boolean logic model. Red nodes are on. The remaining nodes are off.

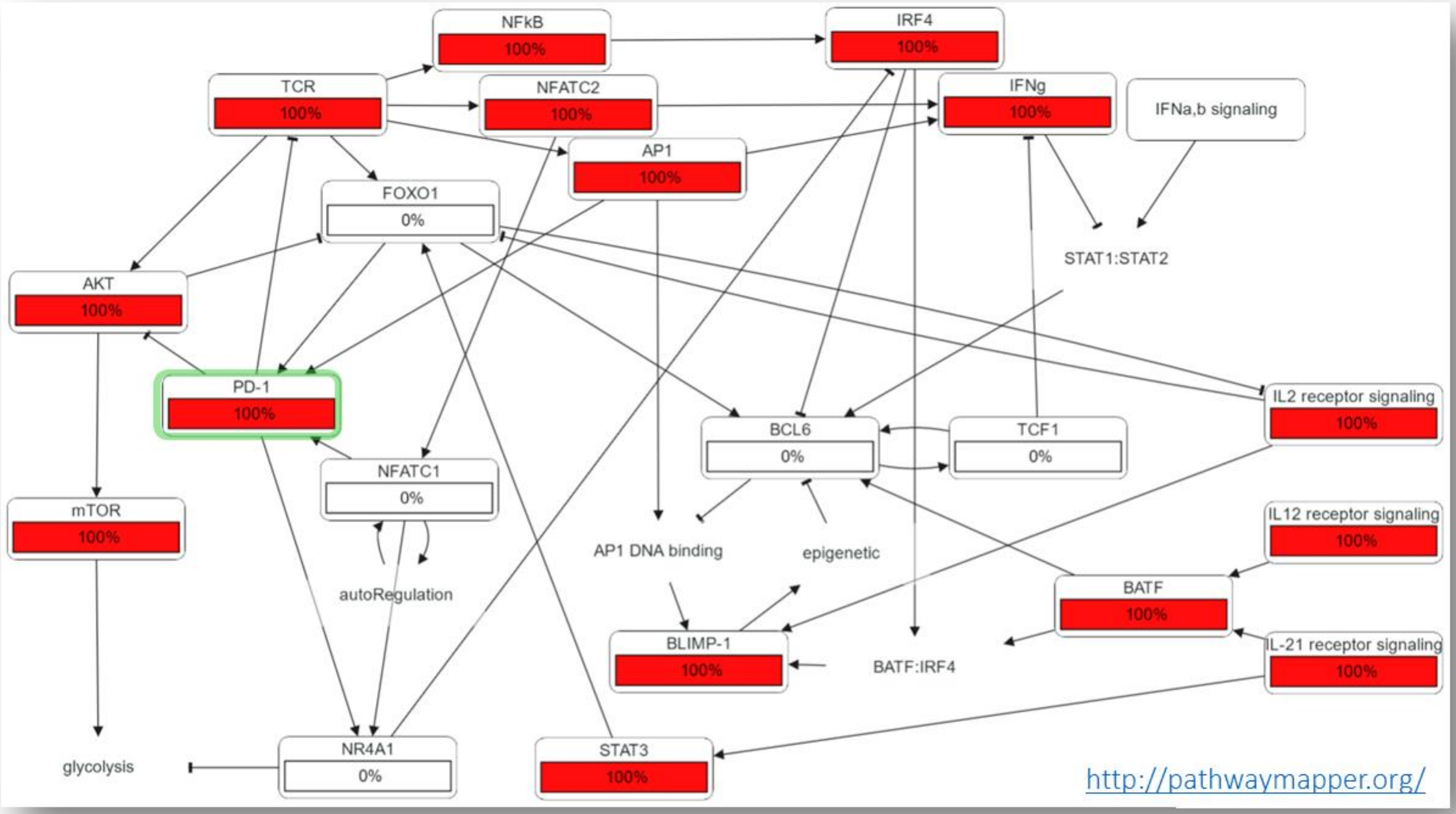

**Supplementary Fig. 20**. Terminal exhaustion state of an example Boolean logic model. Red nodes are on. The remaining nodes are off.

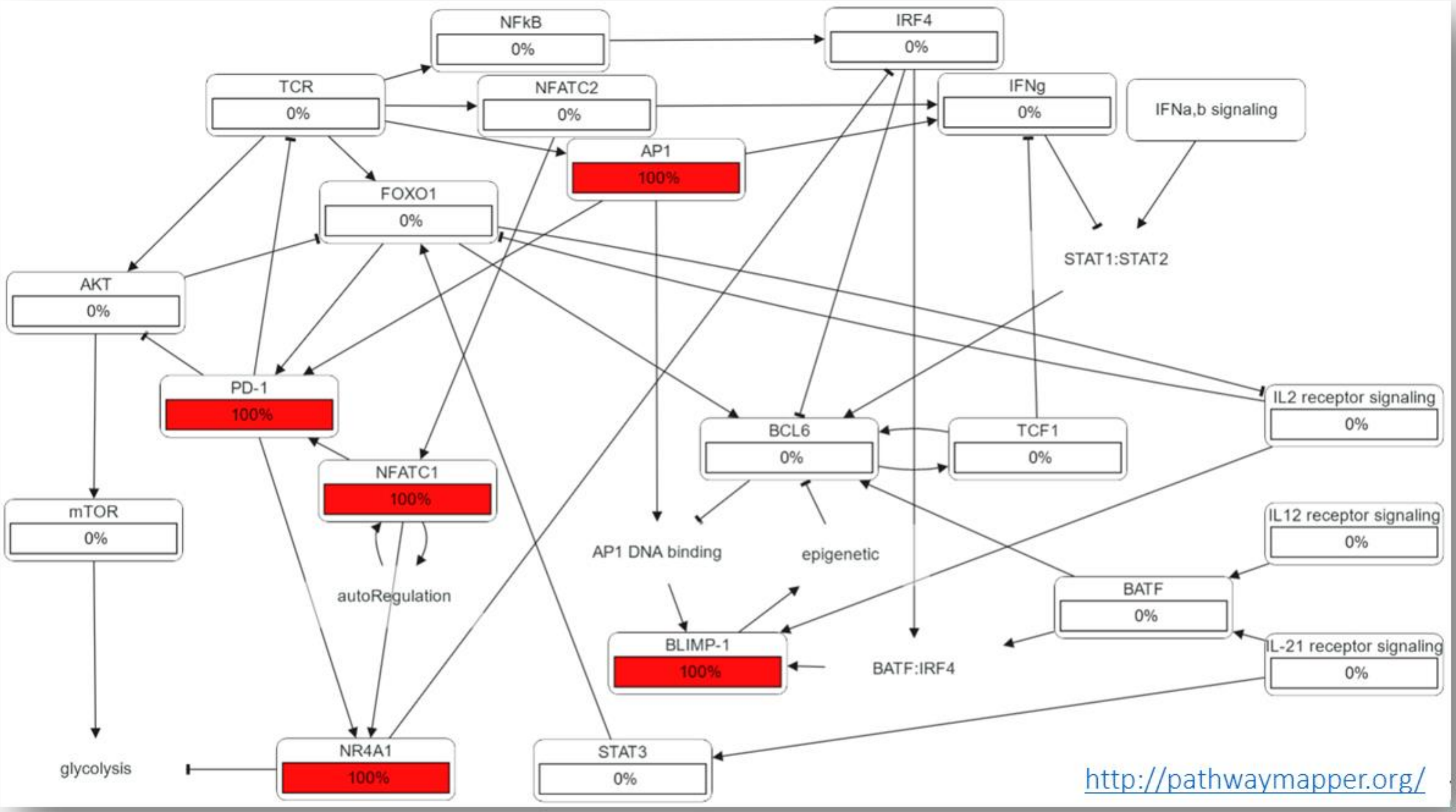

**Supplementary Fig. 21**. Four functional network motifs common in T cell exhaustion: positive feedback, negative feedback, and feed-forward loops are defined at left and schematically presented at right. Mutual inhibition (bottom row) is a special case of positive feedback, exemplified in the TCE network by the mutual inhibition between the BCL6/TCF-1 axis and BLIMP-1. The schematic presented here shows a tube resting on a fulcrum at its middle. If the tube is partially filled with a liquid, it becomes bistable. Any shift of the liquid to one side of the tube will lower the corresponding end of the tube, causing more liquid to flow in the same direction.

***Positive feedback*** refers to any situation where a <u>downstream</u> quantity ***reinforces*** the effects of <u>upstream</u> events

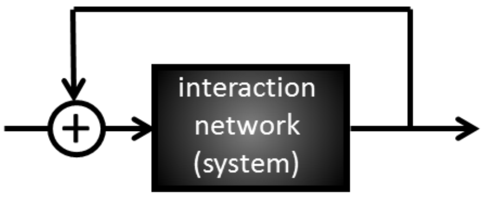


***Negative feedback*** refers to any situation where a <u>downstream</u> quantity ***counteracts*** the effects of <u>upstream</u> events

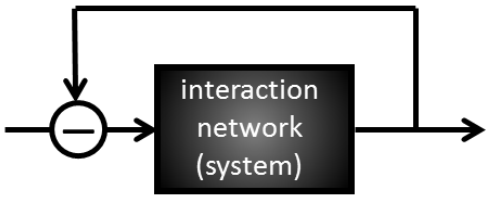


A ***feed-forward loope*** refers to any situation where an <u>upstream</u> quantity regulates <u>downstream</u> events

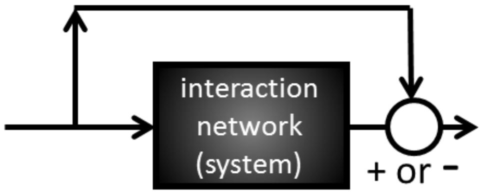


***Mutual inhibition:***

BCL6, TCF-1 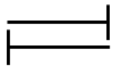 BLIMP-1

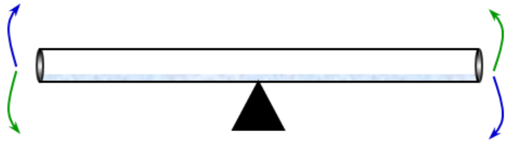

**Supplementary Fig. 22**. The simplified literature-based network after grouping uninformative isoforms and chains. Note that some gene families/isoforms have been collapsed into single nodes (e.g. NF-κB) unless isoforms are known to play distinct roles (e.g. ID2, ID3) during acute/chronic T cell responses.

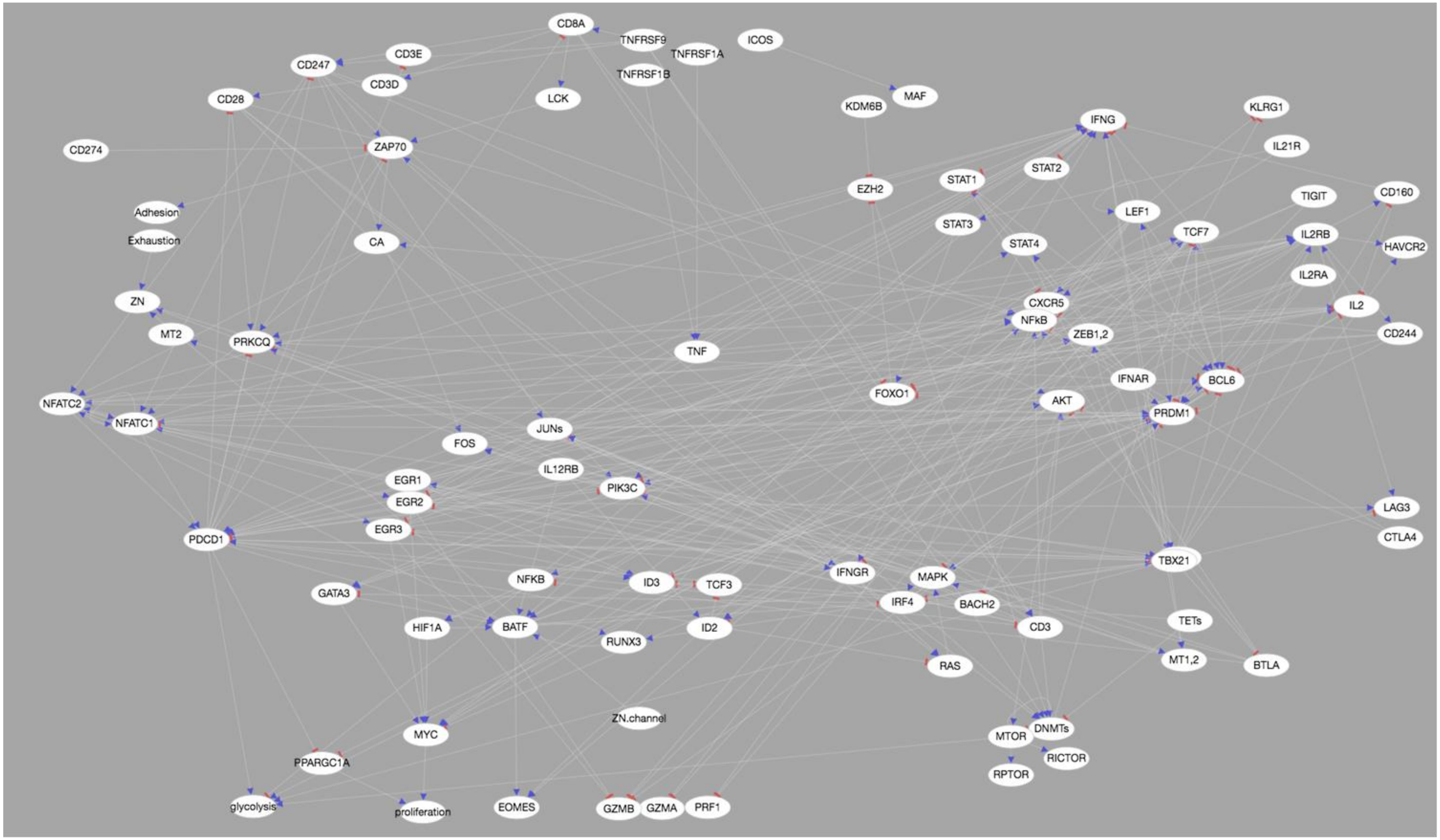

**Supplementary Fig. 23**. Single-cell RNA-seq data suggest the mutual repression of *BCL6* and BLIMP-1 (*PRDM1*) is all-or-nothing (bistable). Panel (**a**) shows a theoretical phase portrait for bistable mutual exclusion (adapted from Bolouri[5]). Red arrows show the direction of change in BCL6 and BLIMP-1 for any given pair of values. Example state trajectories are shown in cyan. Note that all trajectories end at one of two possible steady states (at top-left and bottom-right). Inset shows example time courses for BLIMP-1 (blue) and BCL6 (red) activity levels corresponding to the black trajectory in the main figure. (**b**) In 4,482 of 5,063 single CD8$^{+}$ T cells (89%), either BCL6 or BLIMP-1 mRNA is not detected. Inset shows that only 6% of cells have > 4 BCL6 & PRDM1 reads at the same time. Data from Zheng et al[6].

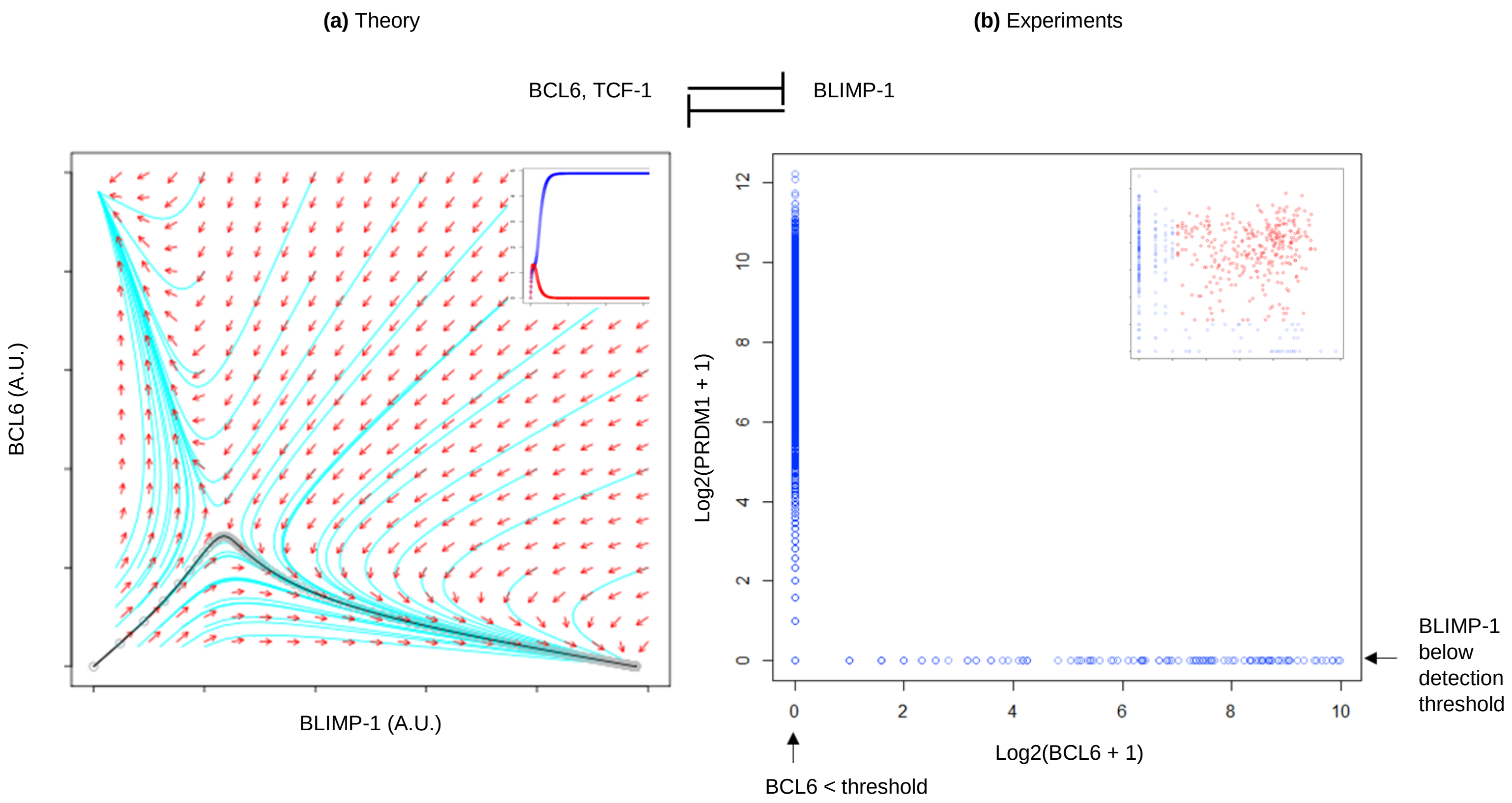

**Supplementary Fig. 24**. Compared to simple direct regulation (**a**), negative feedback can enable faster responses, and more precise response to stimuli (**b**). '$t_{1/2}$' is the time it takes for a nominal gene (*x*) to reach 50% activity. '*b*' is an activating input. The plots in (**a**, **b**) show how the activity of '*x*' varies with the level of the input '*b*' under direct and negative-feedback regulation. In (**a**), the steady state value of *x* (denoted '$x_{ss}$') is a linear function of '*b*'. With negative feedback (**b**), $x_{ss}$ (the point at which the rate of production of '*x*' - blue and green curves - crosses its loss rate - red line) varies non-linearly with '*b*'. As a result, a large change in '*b*' can produce a much smaller change in $x_{ss}$. (Rosenfeld et al[7]).

**(a)** Simple/direct regulation:

$$\frac{dx}{dt} = b - ax$$

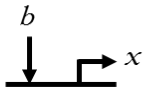


$$x(t) = \frac{b}{a}(1 - e^{-at})$$

$$0 = b - ax_{ss} \Rightarrow x_{ss} = \frac{b}{a} \Rightarrow 0.5\frac{b}{a} = \frac{b}{a}(1 - e^{-at}) \Rightarrow t_{1/2} = \frac{ln2}{a}$$

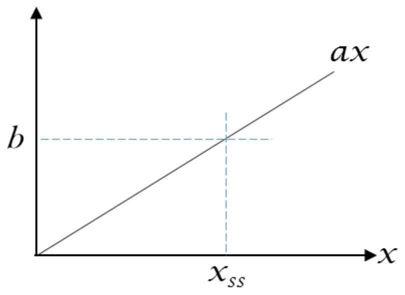


**(b)** Negative feedback:

$$\frac{dx}{dt} = \frac{b}{\left(1 + \left(\frac{x}{K}\right)^n\right)} - ax$$

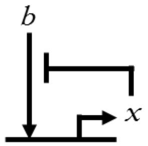


Simplest case: $n = 1$

$$x_{ss} = \frac{\sqrt{K^2 + 4K\,b/a} - K}{2} \approx \sqrt{K\,b/a}$$

$$t_{1/2} \approx \frac{K}{2b}$$ ⇒ Faster response with higher b

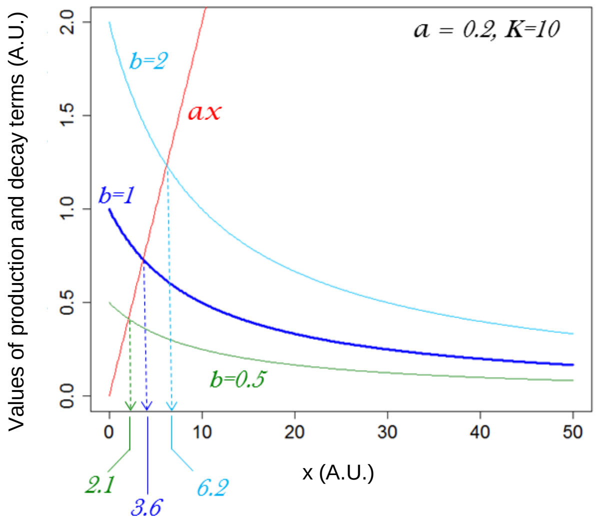

**Supplementary Fig. 25**. A simplified simulation model of negative feedback by inhibitory receptors on CD8⁺ T cells demonstrates the potential benefits of the Negative Feedback functional motif. (**a**) and (**b**) show simplified network diagrams. (**c**) Example simulation results demonstrating the speed gain and robustness properties of negative feedback. Model ODE has the form: $y' = ((11.175+x)/(1+y)) - 0.1*y$, where the parameter values are selected arbitrarily for purely illustrative purposes.

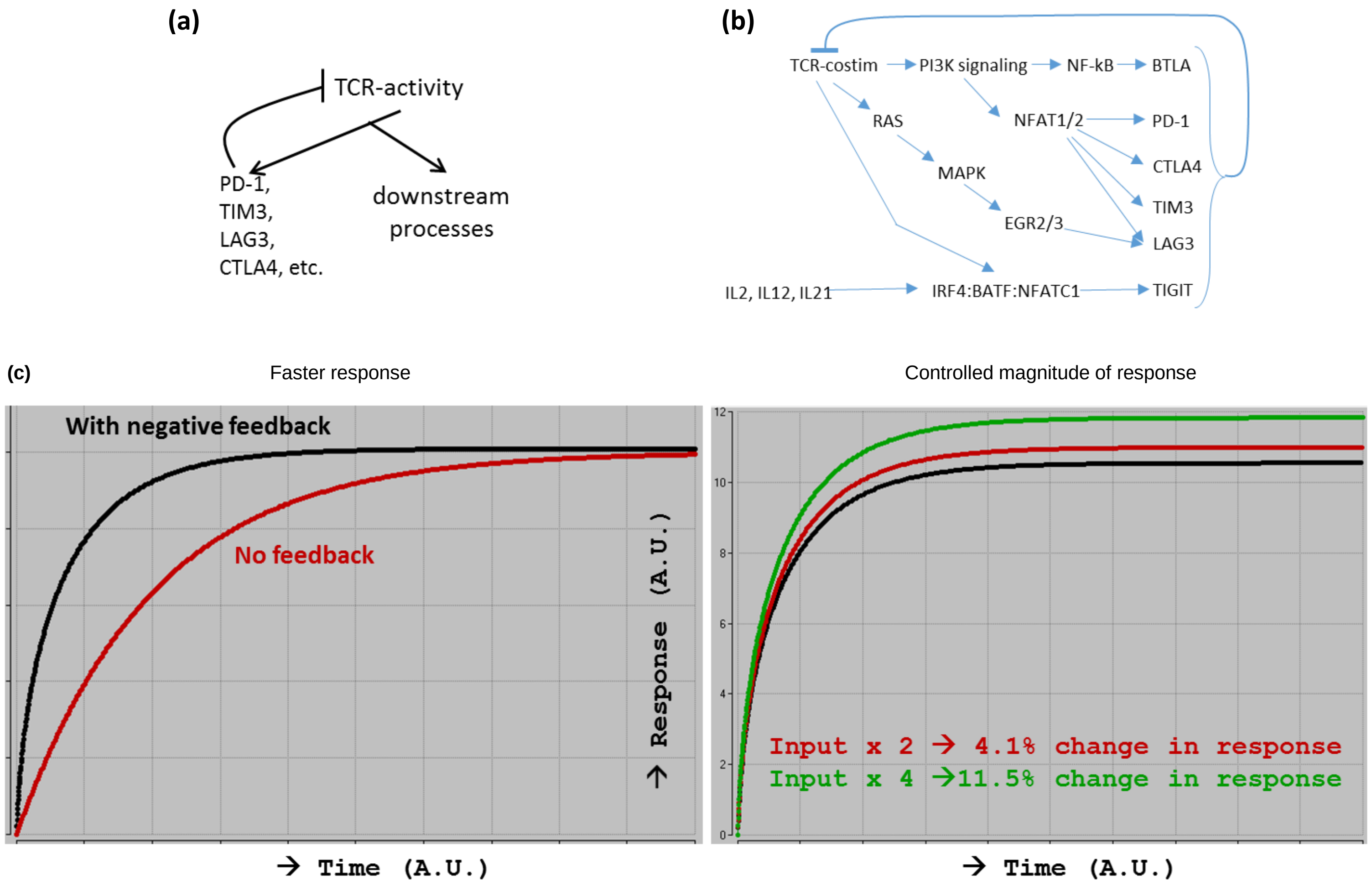

**Supplementary References (for the Supplementary Figures)**


1. Misstear, K. et al. Suppression of antigen-specific T cell responses by the Kaposi's sarcoma-associated herpesvirus viral OX2 protein and its cellular orthologue, CD200. *J. Virol.* **86,** 6246–6257 (2012).
2. Rosenblum, M. D. et al. CD200 is a novel p53-target gene involved in apoptosis-associated immune tolerance. *Blood* **103,** 2691–2698 (2004).
3. Fernández-Sáiz, V. et al. SCFFbxo9 and CK2 direct the cellular response to growth factor withdrawal via Tel2/Tti1 degradation and promote survivial in multiple myeloma. *Nat. Cell. Biol.* **15,** 72–81 (2013).
4. Kurachi, H. et al. Human SPA-1 gene product selectively expressed in lymphoid tissues is a specific GTPase-activating protein for Rap1 and Rap2. Segregate expression profiles from a rap1GAP gene product. *J. Biol. Chem.* **272,** 28081–28088 (1997).
5. Bolouri, H. *Computational Modeling of Gene Regulatory Networks – a Primer* (Imperial College Press, London, 2008).
6. Zheng, C. et al. Landscape of infiltrating T cells in liver cancer revealed by single-cell sequencing. *Cell* **169,** 1342–1356.e16 (2017).
7. Rosenfeld, N., Elowitz, M. B. & Alon, U. Negative autoregulation speeds the response times of transcription networks. *J. Mol. Biol.* **323,** 785–793 (2002).
